# Cell material state determines high-frequency cell deformation and microbubble-induced permeabilization

**DOI:** 10.64898/2026.04.23.720380

**Authors:** S Sloan, O Pattinson, Ben Issa A, E Stride, S Tilley, J Kanczler, D Carugo, F Pierron, ND Evans

**Affiliations:** Centre for Human Development, Stem Cells and Regenerative Medicine, Bone and Joint Research group, University of Southampton; Bioengineering Sciences Group, Institute for Life Sciences, University of Southampton; Botnar Research Centre, Nuffield Department of Orthopaedics, Rheumatology and Musculoskeletal Sciences (NDORMS), University of Oxford; Trauma and Orthopaedics, University Hospital Southampton; MatchID NV, Ghent, Belgium

**Author notes:** Equal contribution (S Sloan, O Pattinson).

**Keywords:** mechanobiology, microbubbles, sonoporation, viscoelasticity, digital image correlation

## Abstract

Mechanical properties influence how cells withstand and transmit force, but their role in clinically relevant, high-frequency therapeutic perturbations remains poorly understood. Ultrasound-stimulated microbubbles can locally deform cells and tissues to enhance drug delivery, yet therapeutic responses vary markedly across mechanical microenvironments. How cellular material properties govern microbubble-cell interactions at megahertz loading rates - well beyond the range accessible to conventional mechanobiology - remains unresolved. Here we combine ultra-high-speed imaging with digital image correlation to map microbubble-induced deformation in living cells at ultrasound frequencies. We show that oscillating microbubbles generate harmonic deformation waves whose spatial decay defines a micrometre-scale attenuation length governed by cytoskeletal organisation and intracellular viscoelastic dissipation. Pharmacological softening or stiffening alters both deformation propagation and wave speed, and these mechanical changes predict permeabilisation efficiency in tumour-derived and primary human bone-marrow stromal cells. These findings establish cellular viscoelasticity as a determinant of microbubble bioeffects and suggest that tissue mechanical state can be used as a design parameter for ultrasound-mediated therapeutic delivery.

## Introduction

Cells and tissues are viscoelastic materials whose mechanical properties influence how they deform, dissipate energy and respond biologically to applied forces ^1^. Most experimental approaches in cell mechanobiology probe these behaviours at quasi-static or low-frequency timescales, yet many biomedical interventions impose mechanical loading at far higher rates ^2,3^. One example of this is therapeutic ultrasound, where acoustic excitation occurs at megahertz frequencies, generating sub-microsecond mechanical perturbations that are beyond the range routinely accessible to conventional cell-mechanics assays ^4^. How living cells transmit and dissipate mechanical energy under these conditions, and whether their material state determines biological outcome, remains poorly understood.

Ultrasound-stimulated microbubbles provide both a clinically relevant actuator and a tractable model for studying this problem. Microbubbles are micrometre-sized, surfactant-stabilised, gas-filled particles routinely used as contrast agents in ultrasound imaging ^5,6^. Beyond diagnostics, they are being developed for targeted drug delivery ^7^ and ultrasound-accelerated thrombolysis ^8,9^. Acting as cavitation nuclei, microbubbles convert incident acoustic energy into mechanical motion ^10^. As an ultrasound pressure wave propagates through fluid, oscillation of the compressible gas core generates local pressure fields and flow that mechanically perturb nearby cells and tissues ^11^. These perturbations can induce transient membrane permeabilisation (eg via pore formation, known as sonoporation), endothelial barrier modulation and extracellular matrix disruption, which have been exploited to deliver macromolecules into cells ^12,13^ and tissues ^11^ , and to accelerate clot breakdown ^14^. Microbubble-enabled ultrasound therapies have progressed to clinical evaluation, including early-phase studies in oncology for minimally invasive, targeted delivery of chemotherapeutics ^15,16^ and disruption of thromboembolism ^17^.

Despite clinical progress, the therapeutic outcomes of ultrasound and microbubble exposure are not uniform, and vary substantially with tissue microenvironmental context ^18,19^. Emerging *in vivo* work suggests that tumour mechanical state - manifesting as stiffness and the perfusion constraints that imposes - can markedly change the effectiveness of ultrasound-microbubble delivery ^18,20^. This raises a fundamental biophysical question - when an oscillating microbubble applies high-frequency mechanical loading to a nearby cell, is the resulting deformation governed only by bubble dynamics, or also by the material properties of the cell itself?

There are strong reasons to expect the latter. Cell deformation under dynamic loading depends on cytoskeletal organisation, membrane–cortex coupling and the frequency-dependent viscoelastic response of the cytoplasm and cortex. ^21^. Indirect evidence supports a role for cell mechanics in the effects of microbubbles, with Fan *et al.* reporting higher permeabilisation efficiency during S phase than in G1 or G2/M, which they attributed to stiffness differences across the cell cycle and altered resistance to cyclic loading ^22,23^. At the tissue scale, microbubble activity in microvessels can transmit deformation tens of micrometres into the surrounding parenchyma ^24^, implying that tissue viscoelasticity can determine both the magnitude and spatial distribution of microbubble-induced strain. Because viscoelasticity varies substantially between tissues and is altered in many disease states by more than an order of magnitude ^1^, the mechanical properties of the microenvironment are likely to influence the clinical efficacy of ultrasonic-microbubble-mediated drug delivery. Consistent with this hypothesis, *in vivo* studies have reported that the efficacy of microbubble-mediated ultrasound delivery can depend strongly on the tumour microenvironment ^18^. To date, however, direct quantitative measurements linking cell mechanical state to microbubble-driven deformation fields and downstream phenotypic effects remain limited, largely because conventional cell mechanobiology assays probe much lower frequencies than ultrasound.

Answering this question quantitatively, however, is challenging because microbubble oscillation occurs at ultrasound frequencies (∼1 MHz), so time-dependent effects must be resolved at sub-microsecond timescales. Ultra-high-speed (UHS) video microscopy at >5LJMfps has therefore been used to visualise cell–microbubble interactions, demonstrating membrane motion near oscillating bubbles ^25^, and the dependence of sonoporation on bubble–cell contact ^26^. These observations support the prevailing view that microbubble-driven membrane strain and strain rate are key determinants of permeabilisation ^27,28^. While UHS imaging has transformed our qualitative understanding, it cannot on its own provide the full-field deformation maps needed to relate spatial strain patterns to cellular mechanics. Digital image correlation (DIC) is a full-field technique for recovering displacement and strain from patterned images ^29^ that can quantify the spatial distribution and magnitude of microbubble-driven deformation, but has not to date been applied to microbubble-induced cell deformations.

Here, we combine UHS imaging with DIC to quantify microbubble-induced cell deformation at ultrasound frequencies. By extracting spatially and temporally-resolved deformation amplitudes, we test whether cytoskeletal mechanics regulate deformation transmission and predict microbubble-induced permeabilisation efficiency. Specifically, we quantify (i) the spatial decay of deformation away from the oscillating bubble and (ii) the propagation speed of the deformation field, then relate these to cytoskeletal perturbations that soften or stiffen cells and to permeabilisation measured by uptake of a cell-impermeant dye. In doing so, we establish a link between cell mechanical state and sonoporation efficiency, providing a mechanistic basis for how tissue- and disease-specific differences in stiffness may govern therapeutic response. More broadly, this framework offers a route to predict and optimise ultrasound–microbubble delivery, and suggests that microbubbles could act not only as therapeutic actuators but also as probes of local mechanical state in living tissues.

## Results

### Sub-microsecond quantification of microbubble-induced cell displacement using DIC

In previous work we developed a custom acoustic chamber enabling imaging of cell-microbubble interactions under ultrasound stimulation at time resolutions exceeding 1LMfps ^30^. Using this platform, we observed dynamic, spatially heterogeneous deformations within the cytoplasm surrounding oscillating microbubbles. We hypothesised that the magnitude and spatial distribution of these deformations may reveal cell mechanical/material properties, and also may relate to the likelihood of biological effects such as sonoporation. Although approaches such as microscopic shear-wave elastography ^31^ and a variety of techniques have been applied to determine cell mechanical properties ^2^, none currently resolve the mechanical response at the MHz frequencies of microbubble oscillation.

To quantify microbubble-induced cell deformation, we combined ultra-high-speed imaging with DIC, exploiting the intrinsic speckle pattern of mammalian cells in bright-field microscopy. MG63 cells were seeded in the device’s fluidic chamber (Figure 1a). After chamber inversion to promote bubble-cell contact, frequent cell-microbubble interactions were observed. Under ultrasound stimulation, microbubbles oscillated synchronously with the driving acoustic frequency, imparting visible displacement to the cell body and cytoplasm detectable by video microscopy and grey-scale fluctuation analysis (Figure 1b, Video 1 and Supplementary Figure 1). Because bright-field images of cells contain intrinsic speckle, DIC could be applied without exogenous fiduciary markers. Using MatchID software, directional and absolute displacements were calculated across the entire cell for each frame of the UHS recordings (Video 2, Video 3 and Video 4).

**Figure 1:**
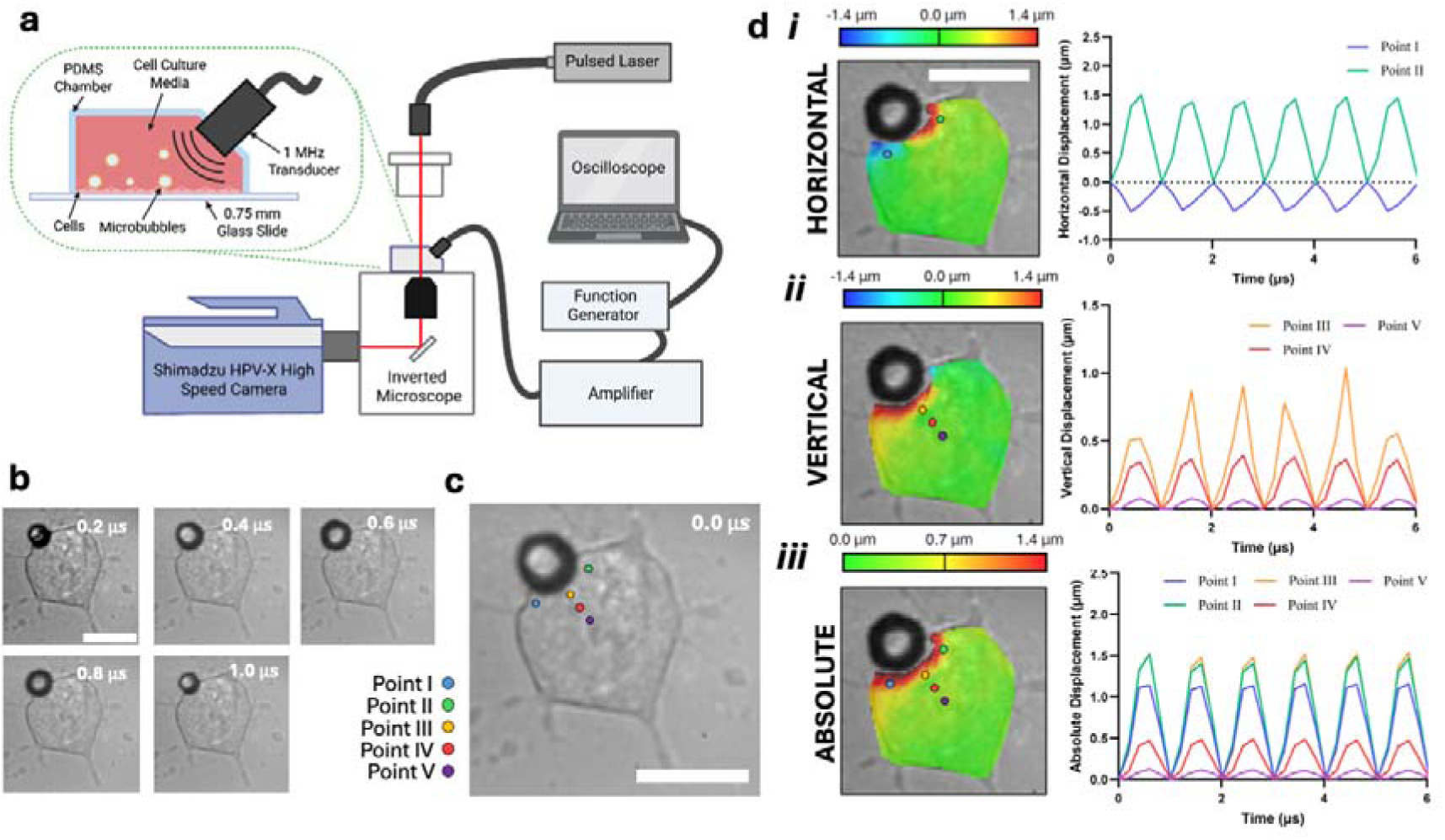
Sub-microsecond quantification of microbubble-induced cell displacement using DIC. **a**) Schematic of the acoustic device developed to induce cell-microbubble interactions under an ultrasound field, placed within the high-speed imaging microscopy apparatus. **b**) The first 5 frames of the Shimadzu HPV-X images captured at 5 million FPS showing an example of a cell-microbubble interaction, with three distinct regions in the cell identified (Video 1, scale bar = 10 µm) **c**) Reference image of the cell-microbubble interaction (scale bar = 50 µm) with 5 points defined within the cell around the microbubble, at different orientations and distances. **d**) DIC spatial maps in the region of interest showing the **i**) horizontal displacement (Video 2), **ii**) vertical displacement (Video 3), and **iii**) absolute displacement (Video 4) (scale bar = 25 µm).

To quantify the magnitude of cell displacement with respect to the microbubble, we examined a series of individual subsets within the cell body that varied in distance and distribution from the microbubble edge (Figure 1c). Horizontal displacement in subsets on approximately opposite sides of the microbubble was similar in magnitude but opposite in direction (Figure 1d(i)). The peak vector displacement during each oscillation cycle was -0.5 ± 0.02 µm, on the left side of the cell and 1.5 ± 0.09 µm on the opposite side (points I and II; Figure 1d(i)). Vertical displacement declined as a function of distance perpendicular to the microbubble periphery, from 0.9 ± 0.2 µm at point III to 0.4 ± 0.02 µm at point IV to 0.1 ± 0.02 µm at point V (Figure 1d(ii)). Absolute displacement oscillated at the same frequency as the microbubble and showed comparable amplitudes at points equidistant from the bubble boundary, but varied with radial angle (magnitude of displacement at 1.21 ± 0.01 µm, 1.49 ± 0.09 µm, 1.5 ± 0.08 µm, 0.49 ± 0.02 µm, and 0.17 ± 0.02 µm for points I, II, III, IV and V, respectively; Figure 1d(iii)). Together, these results demonstrate that DIC can quantify spatial distributions of cell deformation during microbubble oscillation without exogenous markers, providing a framework for quantitative analysis of microbubble-cell mechanical interactions.

### Microbubble-induced dynamic cell displacement is sinusoidal and its amplitude decays exponentially with distance

As ultrasound at MHz frequencies induces sinusoidal oscillation in microbubbles, we hypothesised that adjacent cells would undergo similar deformation. To confirm this, microbubble edge-detection data were fitted to a sinusoidal function, Eq. 1 (Fig. 2a(i–ii)). The fit showed strong agreement (R² = 0.9720) with an oscillation frequency of 0.99 MHz, matching the driving ultrasound frequency, and a recovered oscillation amplitude of 1.03 µm representing the mean radial expansion and contraction over the video sequence. Next, the spatial pattern of the induced deformation oscillation amplitude throughout the entire cell was determined by performing FFTs for every subset of data generated via MatchID (Figure 2b). Points near the microbubble exhibited harmonic oscillations at 1 MHz with peak displacement of 1.06 µm, comparable to the bubble oscillation (Figure 2b(i)). The amplitude of the 1 MHz FFT component was therefore extracted for all subsets and mapped to reconstruct the spatial deformation field across the cell (Figure 2b(ii-iv)). In materials under dynamic loading, spatial decay of deformation away from the loading point reflects attenuation of a propagating mechanical wave and can reveal damping properties ^32^. Quantifying the displacement around the microbubble with respect to, for example, microbubble size or amplitude of oscillation, may offer the opportunity to determine the rheological properties of complex cell structures at high frequency, which has been challenging in the past ^2^.

**Figure 2:**
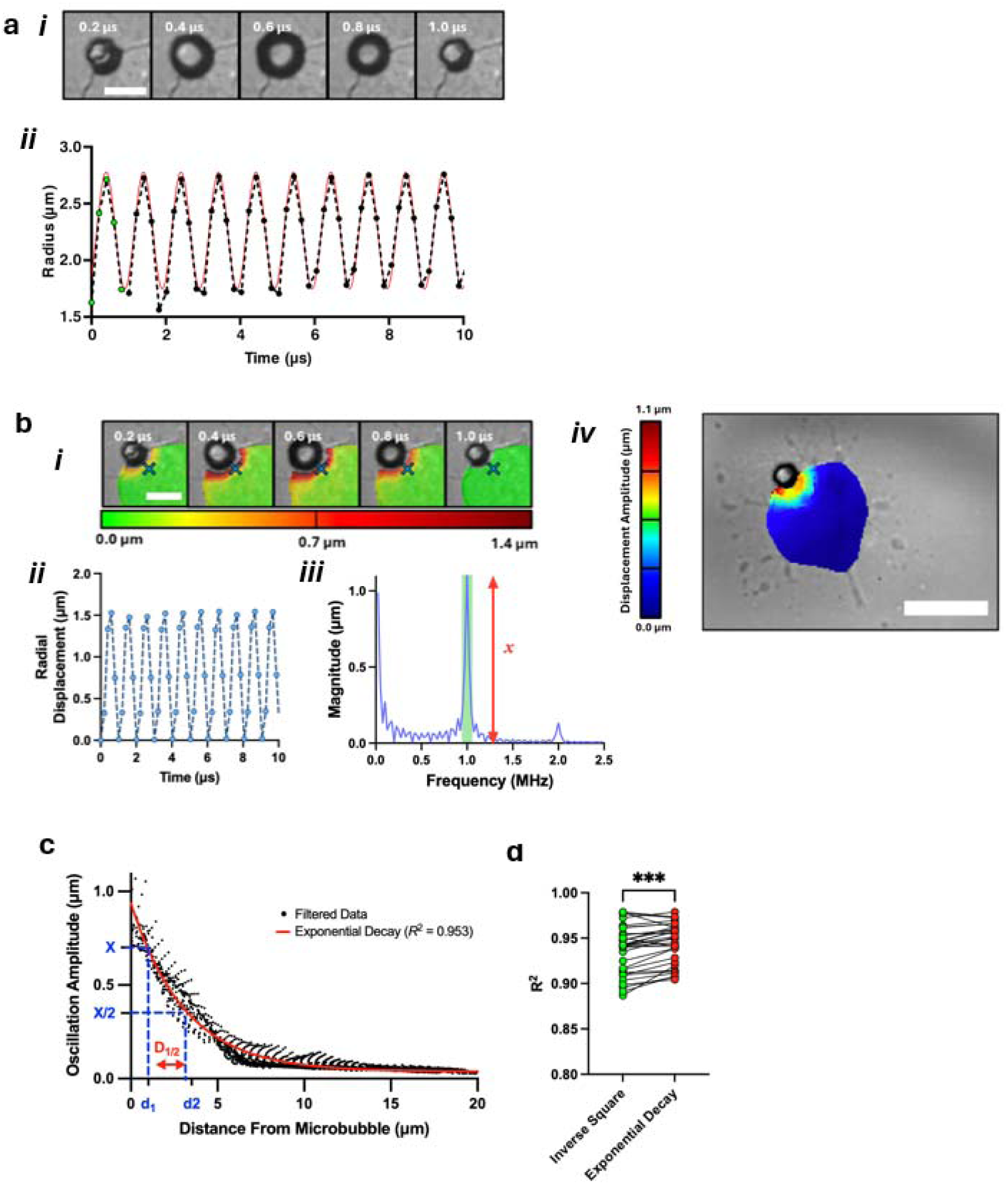
Microbubble-induced dynamic cell displacement is sinusoidal and its amplitude decays exponentially with distance. a. **i**) First 5 frames showing a single microbubble oscillation cycle (scale bar = 10 µm). **ii**) The first 10 µs of the microbubble oscillation measurements with a fitted sine function. The first 5 green data points correspond to the frames shown in **i**. **b**) **i**) 5 frames showing the absolute displacement over a single oscillation cycle in the DIC map (top, scale bar = 5, (Video 4). The cross refers to a point near the microbubble, where **ii**) an example plot of radial deformation against time is shown. From this, **iii)** an FFT of the radial deformation is produced, where **iv)** the magnitude at 1MHz is mapped across the cell’s surface, showing the deformation induced by the microbubbles’ oscillation (scale bar = 25 µm). **c**) Induced displacement throughout the cell with distance from the microbubble, with inverse square and exponential decay models fit. **d**) R^2^ values for each model compared across all data sets. (* = p < 0.05, ** = p < 0.01, *** = p < 0.001, **** = p < 0.0001).

We therefore examined the relationship between displacement amplitude and distance from the microbubble boundary. In the example cell (Fig. 2b), displacement decreased smoothly from ∼1 µm at the bubble interface to ∼0 µm at ∼25 µm distance (Fig. 2c). Two models were tested to describe this decay: exponential (Eq. 2) and inverse-square (Eq. 3). Across datasets, the exponential model provided a significantly better fit (R² = 0.944 ± 0.022 vs 0.937 ± 0.026; p < 0.001; n = 31) (Fig. 2d). Using this model we defined the half-decay distance, hereafter which we name “attenuation length scale”, D_1/2_ (Eq. 4), yielding a mean value of 2.85 ± 0.69 µm (n = 31), (Figure 2c; note that all decay data is provided in Supplementary Information 1). Exponential attenuation is characteristic of wave propagation in viscoelastic media, where internal friction dissipates mechanical energy and produces spatial decay of deformation ^33,34^. The cytoplasm is commonly described as a complex viscoelastic composite comprising cytoskeletal filaments embedded in viscous cytosol. Relative motion between filaments, crosslinkers, organelles and surrounding fluid generates frequency-dependent damping. While cellular viscoelasticity is typically measured by microrheology at frequencies below ∼100 kHz ^3^, our measurements infer dissipation from deformation decay under MHz excitation. These results therefore suggest that intracellular architecture acts as a lossy viscoelastic transmission medium. If cytoskeletal organisation dominates dissipation and load transfer, we next hypothesised that perturbing it should systematically shift the attenuation length scale D_1/2_, a parameter rarely accessible for living cells under ultrasound-rate (MHz) loading.

### Cell material properties determine D_1/2_ and microbubble-induced permeabilisation

To test this hypothesis, cells were treated with agents that disrupt cytoskeletal structure and reduce stiffness (cytochalasin D, latrunculin A, blebbistatin and Y-27632) or with agents that promote actin polymerisation and increase stiffness (calyculin A and nocodazole). Confocal imaging of phalloidin-stained cells confirmed a reduction or increase in filamentous actin in cells treated with softening or stiffening agents, respectively (Figure 3a and Supplementary Figure 2).

**Figure 3:**
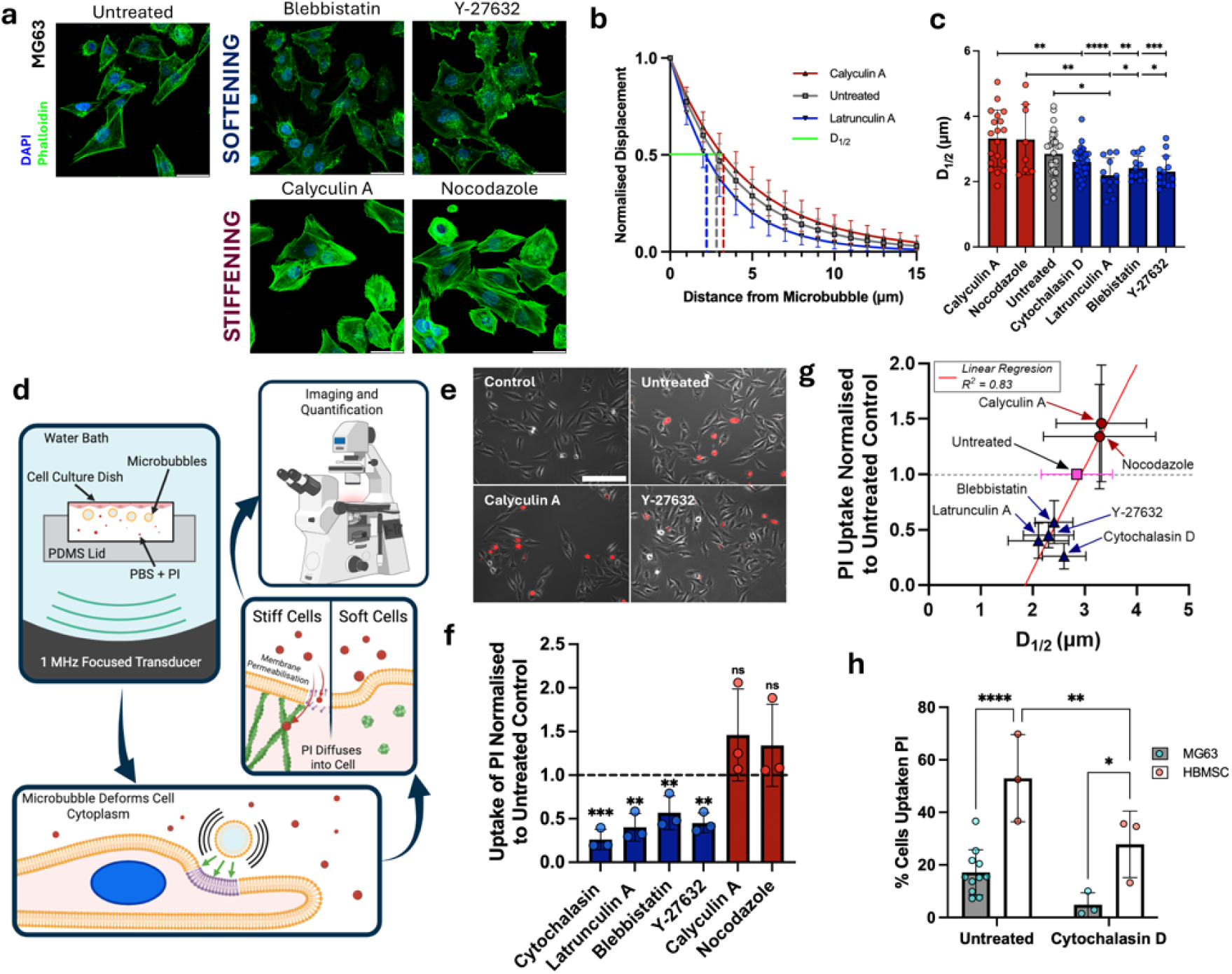
Disruption of the cell cytoskeleton reduces D_1/2_ and protects against microbubble-induced permeabilisation. **a**) Confocal images with DAPI-stained nuclei (blue) and phalloidin-stained actin (green) for untreated, blebbistatin, Y-27632-treated (softening agents) and calyculin A or nocodazole-treated (stiffening agents) MG63 cells (scale bar = 50 µm). **b**) D_1/2_ was reduced in softened cells and increased in stiffened cells, as exemplified by comparison of decay curves for untreated and latrunculin or calyculin A-treated cells **c**) Comparison of D_1/2_ across all treatment groups. **d**) A schematic showing the experimental approach to test ultrasound microbubble-induced permeabilisation as a function of treatment with softening or stiffening agents and **e**) PI uptake in control, untreated, Y27632 and calyculin A-treated cells, with **f**) showing the significant decrease in permeablisation in softened cells relative to untreated or stiffened cells. **g)** Normalised permeabilsation correlates with D_1/2_ modulated with softening or stiffening agents compared to controls. **h**) Effect of cytochalasin D on permeabilisation efficiency in HBMSCs compared with the MG63 cells, showing a consistent effect in both cell types but a significantly high permeabilisation effect in HBMSCs relative to MG63s. (* = p < 0.05, ** = p < 0.01, *** = p < 0.001, **** = p < 0.0001).

To quantify the effect of these stiffness modifications on deformation propagation, we calculated mean values of the attenuation length scale, *D_1/2_*. Normalised decay curves revealed clear differences between groups, with stiffening agents increasing *D_1/2_*and softening agents decreasing it (Figure 3b). All individual groups are shown in Fig. 3c and statistical analyses in Supplementary Table 1. Pooled analysis showed that *D_1/2_* was significantly larger in cells treated with stiffening agents (calyculin A and nocodozole) than in cells treated with softening agents (cytochalasin D, latrunculin A, blebbistatin and Y-27632; 3.31 ± 0.92 µm vs. 2.44 ± 0.46 µm; *p* < 0.0001). Untreated cells showed intermediate behaviour, differing significantly from both softened (p = 0.010) and stiffened (p = 0.019) groups.

These data indicate that cytoskeletal organisation does regulate microbubble-driven cell deformation. In viscoelastic media, changes in attenuation length reflect shifts in how mechanical energy is transmitted and dissipated through the material, with the decay length capturing the balance between loss and load-bearing pathways in the cytoplasm ^3,35^. That these differences remain resolvable under ultrasound-rate (MHz) stimulation demonstrates that intracellular dissipation pathways determine deformation propagation even at the extreme strain rates imposed by oscillating microbubbles. The direction of the effect - more rapid attenuation in softened conditions - mirrors trends often reported at the tissue scale in shear-wave elastography, where softer biological materials commonly exhibit stronger attenuation ^36,37^ Although the material properties of viscoelastic systems have rarely been measured directly at MHz frequencies, theory suggests that the bulk response becomes increasingly important in this regime ^33,35^. Nevertheless, our data demonstrate that cell stiffness modulates deformation attenuation even at these much higher frequencies.

We next asked whether the cellular mechanical state also regulates microbubble-induced permeabilisation. While indirect evidence suggests that cell stiffness influences microbubble-induced permeabilisation ^22,23,38^ this has not (to the best of our knowledge) been tested directly. MG63 cells were treated with the same panel of softening or stiffening agents and exposed to ultrasound in the presence of microbubbles and the cell-impermeant dye propidium iodide (PI) (Figure 3d). Compared with untreated cells (17.14 ± 8.62 %), softening treatments significantly reduced permeabilisation efficiency: cytochalasin D, 4.82 ± 4.58 % (p < 0.01), latrunculin A, 5.13 ± 0.64 % (p < 0.01), blebbistatin, 9.04 ± 1.27 % (p < 0.05), and Y-27632, 8.86 ± 3.49 % (p < 0.01) (Figure 3e,f). Although stiffening treatments produced a non-significant increase, when permeabilization was plotted as a function of *D_1/2_* there was a clear correlation, with an R^2^ = 0.83 (Figure 3g). No significant PI uptake occurred in control experiments with drug treatment alone or ultrasound in the absence of microbubbles (Supplementary Figure 3).

Tumour-derived cell lines such as MG63 are typically softer than primary stromal cells such as HBMSCs, which may influence their susceptibility to microbubble-induced permeabilisation ^39^. Consistent with this, cytochalasin D treatment significantly reduced PI uptake in primary human bone marrow stromal cells (HBMSCs; 53.0 ± 13.5 % vs 27.8 ± 10.3 % (p < 0.01), similar to that found in MG63 cells (Figure 3h). Significant differences were also observed between MG63 and HBMSC populations in both untreated (p < 0.0001) and cytochalasin-treated (p < 0.05) groups. Controls again showed no PI uptake from drug treatment alone or ultrasound without microbubbles (Supplementary Figure 3).

Together, these data indicate that cellular mechanical state influences microbubble-induced permeabilisation, with pharmacological softening reducing PI uptake in parallel with shortening the attenuation length. This is possibly due to a reduced area of the membrane experiencing stretching, and a similarly reduced area active for microbubble-induced permeabilisation, hence the reduction in uptake. Another possible explanation involves cytoskeletal control of membrane–cortex coupling and prestress. During microbubble oscillation, radial bubble expansion produces deformation normal to the cell surface, generating membrane strain parallel to the membrane. In untreated cells, the actin cytoskeleton forms an interconnected network anchored to the membrane via integrins, constraining membrane motion so that applied forces generate higher local stresses and increase susceptibility to rupture. Disruption of the actin network reduces this prestress and structural constraint, allowing the membrane to deform more and absorb energy without failure. Future experiments using membrane-tension probes or additional fluorescent markers may help resolve these mechanisms ^40–42^.

### Spatial phase variation reflects the propagation of the displacement wave through the cell

For deformation to propagate through the cell, mechanical energy must be transmitted as a travelling deformation wave. We therefore examined the phase spectrum of the FFT of the radial component of the deformation (expressed in polar coordinates about the microbubble centre) and mapped the phase at 1 MHz, across the cell. A gradual phase shift with distance from the microbubble was observed (Figure 4a), consistent with propagation of a deformation wave. Plotting against distance from the microbubble revealed an approximately linear relationship (Figure 4b), allowing the wave propagation speed to be calculated from the phase gradient using Eq 5.

**Figure 4:**
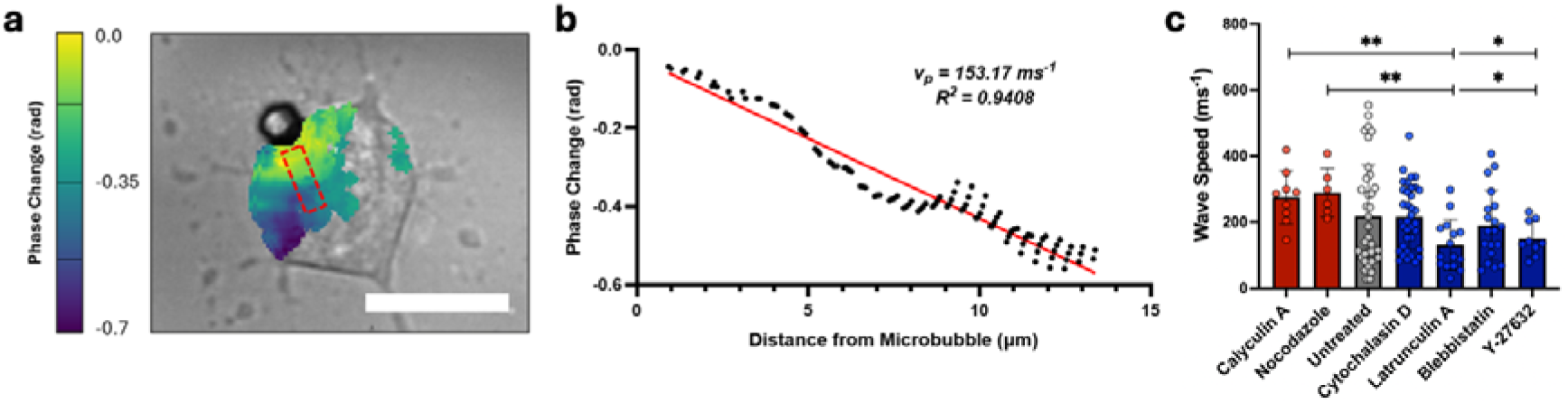
Mechanical waves can be observed transiting MG63 cells and wave speed depends on material properties of the cell. **a**) Spatial colour map of the phase change. A region of interest has been highlighted where a gradual change in phase was observed with distance from the microbubble (scale bar = 25 µm). **b**) Phase change plotted against distance from the microbubble for the highlighted region of interest. A linear regression (Red line) is applied to the data to recover wave velocity. **c**). Wave velocity is affected by chemicals that modify the cytoskeleton, with slower velocities recorded in softened compared to stiffened cells.

Across all untreated microbubble-cell interactions, the average propagation speed was 253 ± 216 ms^-1^ (n = 31). This velocity is substantially lower than compression wave speeds in soft tissues (1450 to 1600 ms^-1^) but much higher than typical shear-wave speeds 1 and 10 ms^-1^ ^36^. The intermediate velocity may suggest the possible existence of non-classical mechanical wave modes, such as surface or Lamb waves, including possible contributions from the glass substrate. At MHz frequencies, theoretical and experimental treatments suggest that shear-wave behaviour in viscoelastic soft materials becomes strongly attenuated ^43^, such that the measured response may increasingly reflect bulk/compressional contributions and geometry-dependent guided modes.

Experimental factors may also influence the measured speeds. Widefield microscopy collects light from above and below the focal plane (≈0.5□µm), and deformation waves generated by the microbubble–cell interaction may propagate obliquely relative to the imaging plane. Both effects would lead to an underestimation of the true propagation velocity and an overestimation of the attenuation length. Mechanically, propagation speeds were higher in cells treated with stiffening agents (calyculin A and nocodazole) than in cells treated with softening agents (cytochalasin D, latrunculin A, blebbistatin and Y-27632; 281.3□±□75.2 vs 182.3□±□97.6□m□s□¹; p□=□0.0066). Speeds were also greater in stiffened than untreated cells (281.3□±□75.2 vs 217.8□±□157.6□m□s□¹; p□=□0.0270), while softened cells showed a non-significant reduction relative to untreated cells. Together with the shorter attenuation length observed in softened cells, these results are consistent with increased mechanical dissipation and reduced load transfer following cytoskeletal disruption (Figure 4c). It is also worth noting that there was a larger variance observed in untreated cells compared to those treated with pharmacological inhibitors. This may reflect greater heterogeneity in cell cycle state in these populations due to differences in stages of cell cycle (which are unlikely present in treated cells), as cell cycle stage can significantly affect cell stiffness and has been implicated as a factor affecting microbubble-induced permeabilisation ^22,23^.

The reduced wave speed indicates increased damping associated with changes in effective shear or bulk modulus, depending on the wave mode involved. Although experimental data remain limited, Grasland-Mongrain *et al.* measured optical shear-wave propagation in isolated murine oocytes and reported a lower shear modulus in cytochalasin-treated cells compared with controls ^31,44^ consistent with increased damping in softer cells. However, the excitation frequencies in shear-wave elastography (∼15□kHz) are orders of magnitude lower than those in our experiments (∼1□MHz). Because shear-wave behaviour is strongly frequency-dependent, attenuation of shear waves in our experiments is significant, and we remain uncertain as to whether the observed displacements are caused by shear waves. Nonetheless, the differences observed here are most plausibly explained by cytoskeletal structural changes that alter energy transmission and dissipation within the cytoplasm.

In summary, our results show that the response of cells to oscillating microbubbles is governed not only by bubble dynamics but also by the material properties of the cell. By combining ultra-high-speed imaging with digital image correlation we quantified microbubble-induced deformation fields at ultrasound frequencies and identified measurable descriptors of this response - the attenuation length scale (D□/□) and the propagation speed of the deformation wave. Both parameters were modulated by cytoskeletal organisation, demonstrating that intracellular viscoelasticity determines how mechanical energy introduced by microbubble oscillation is transmitted and dissipated within the cell.

Importantly, these mechanical changes were accompanied by corresponding changes in microbubble-induced permeabilisation, linking MHz-rate deformation physics directly to sonoporation outcome. These findings therefore suggest that tissue mechanical state may be an important determinant of the heterogeneous therapeutic response observed in ultrasound–microbubble delivery. More broadly, the approach presented here raises the possibility that a better knowledge of tissue mechanical properties may enable better tuned parameters for maximising the chance of microbubble-induced permeabilsiation, including sonoporation, and therefore therapeutic success. In addition, oscillating microbubbles could serve not only as therapeutic actuators but also as local mechanical probes. Extending similar measurements to microvessels or intact tissues, coupled with appropriate models of wave propagation and boundary conditions, may ultimately enable recovery of effective tissue mechanical properties during ultrasound-mediated therapies.

## Methods

### Acoustic Device Manufacture

The design and manufacture of the acoustic device have been described previously ^30^. Briefly, a polydimethylsiloxane (PDMS) manifold was prepared by mixing PDMS precursor and curing agent (Sylgard 184, Farnell, Leeds, UK) at a 10:1 weight ratio, followed by degassing in a desiccator connected to a vacuum pump. A 3-D-printed poly(lactic acid) (PLA) mould was made using an Ultimaker S5 printer and filled with liquid PDMS, which was then cured for 48-72 hours. The solid PDMS manifold was subsequently plasma-bonded to a 170 µm thick (length × width: 75 × 25 mm) glass cover-slide, to create a water-tight chamber. The manifold included an inlet into which a 10 mm element-size transducer (Precision Acoustics, Dorchester, UK) could be inserted to deliver acoustic stimulation through the fluid-filled chamber. These devices had reproducible architectures over multiple manufactured replicas. Over the course of the study, a new device was fabricated each time a different type of cell-microbubble interaction was captured.

### Microbubble Production

Air-filled microbubbles were produced using a 2-stage sonication technique adapted from a previously described method ^45^. Briefly, 1,2-distearoyl-sn-glycero-3-phosphocholine (DSPC) and polyoxyethylene (40) stearate (PEG-40s) were first combined at a molar ratio of 9:1 from stock concentrations of 10 mg/mL (in chloroform). The organic solvent was left to evaporate overnight, and the obtained dry lipid film was hydrated in Dulbecco’s phosphate-buffered saline (DPBS). The lipid dispersion was then sonicated using a Fisherbrand™ Model 120 Sonic tip sonicator (tip diameter: 3.175 mm, Fisher Scientific, Leicestershire, UK). A first sonication stage at 40% intensity for 2.5 minutes at the base of the suspension was followed by a sonication at 70% intensity for 30 s with the tip positioned at the liquid-air interface. The resulting microbubble suspension was subsequently placed in ice until use.

### Cell Culture

MG-63 cells were cultured in Dulbecco’s modified Eagle’s medium (DMEM) with 10% (v/v) fetal bovine serum (FBS), 50 IU/mL penicillin and 50 μg/mL streptomycin sulphate in T75 culture flasks. Primary human bone marrow stromal cells (HBMSC) were isolated from patients undergoing hip replacement surgery at Southampton General Hospital and Spire Southampton Hospital (with local ethics committee approval from the University of Southampton and Research Governance Office (ERGO 31875.A12) and the Northwest-Greater Manchester East Research Ethics Committee (18/NW/0231)). The bone marrow was washed multiple times in Minimum Essential Medium α (α-MEM) and poured through a 0.6 µm cell strainer to remove fat and bone fragments. HBMSCs were cultured in α-MEM with 10% fetal bovine serum (FBS) and 50 IU/mL penicillin and 50 μg/mL streptomycin sulphate. Once at 80% confluence, the cells were split and seeded into the manufactured acoustic devices at seeding densities of 30,000 cells per cm^2^. The devices were filled with 2 mL of the required culture medium, and the cells within the manufactured devices were kept in a 5% CO_2_ incubator for 24 hours. Cells were also seeded at a density of 40,000 cells/cm^2^ for microbubble-induced permeabilisation experiments in a 35 mm glass Ibidi cell culture dish (Thistle Scientific, Glasgow, UK).

### Modulation of cytoskeletal integrity

To investigate the effect of the material properties of cells on microbubble-induced cell strain, cells were treated with various cytoskeletal inhibitors that are reported to reduce or increase cell stiffness (as measured by quasi-static techniques, like atomic force microscopy). To reduce cell stiffness MG63 cells were incubated in DMEM for 30 minutes at 37 °C with either 2 µM cytochalasin-D (Merck, Dorset, UK), or 1 µM latrunculin-A (Merck, Dorset, UK) to disrupt actin polymerisation; 7 µM blebbistatin (Merck, Dorset, UK) to disrupt myosin-II; or 5 µM Y-27632 (Tocris, Bristol, UK) to disrupt Rho-associated, coiled-coil-containing protein kinase (ROCK). To stiffen cells, MG63 cells were incubated in DMEM for 15 minutes at 37 °C with 2 nM Calyculin A (Merck, Dorset, UK), or 1 µM Nocodazole (Merck, Dorset, UK), which promotes actin polymerisation. HBMSCs were treated in α-MEM for 30 minutes at 37 °C with 2 µM cytochalasin-D (Merck, Dorset, UK) to disrupt actin polymerisation and soften the cells. Cells were washed in DPBS, and they were imaged with microbubbles immediately after treatment.

### Ultra-High Speed Imaging of Cell-Microbubble Interactions

To image cell-microbubble interactions, the acoustic device containing cultured cells was filled with 8 mL of DPBS, supplemented with 100 µL of microbubble suspension (∼1×10^7^ microbubbles per mL). The device was sealed by inserting an immersion transducer into the dedicated port of the PDMS manifold and was inverted for up to 5 minutes to induce the natural adhesion between microbubbles and cells, then reinverted to leave approximately 1-2 microbubbles adhered per cell. The device was placed on the stage of an Olympus IX 71 microscope (Olympus, Tokyo, Japan). A Hypervision HPV-X camera (Shimadzu, Kyoto, Japan) was attached to the microscope’s camera port, and pulsed laser illumination was provided by a Cavilux pulsed diode laser (640 nm wavelength, 400 W, Cavitar Ltd, Tampere, Finland). A single 100-cycle pulse ultrasound wave was generated by a 1 MHz transducer (Precision Acoustics, Dorchester, UK), driven by a power amplifier fed by a signal generator (Aim-TTi, Cambridgeshire, UK). A HS3 Handyscope oscilloscope (TiePie, Sneek, The Netherlands) was employed to monitor the electrical signal driving the transducer. The acoustic stimulation also triggered the camera to image at 5 million FPS under an objective of magnification 80×, capturing the dynamic interaction of stimulated microbubbles and cells. Candidate cells with microbubbles in close proximity were identified by light microscopy imaging, and UHS videos of 128 frames were acquired by the camera, with each frame at an image resolution of 400 × 250 pixels. To analyse the images prior to DIC application, the grey level (an intensity value between 0 and 65536, indicating the light level of each pixel) of specific regions within images were compared spatially and temporally to quantify dynamic changes within the images.

### Digital Image Correlation

For measurements of microbubble-induced displacement of the cell, MatchID software (MatchID 2D, 2024.2.5, Gent, Belgium) was used to perform digital image correlation (DIC) using the speckle pattern naturally present in the cells. The initial frame of each UHS video was used as the reference frame, and a region of interest was defined, encompassing the cell to be studied. DIC was performed using a zero-normalised sum of squared differences (ZNSSD) correlation with an affine shape function, with a subset size of 17 and a step size of 1 (see full DIC reporting table in Supplementary Table 2). The subset size determined the size in pixels of a region of the image which was analysed as a single point, and each subset generated a single value of displacement data for each frame, with the step size determining the spacing between analysed subsets and, therefore, the final density of the grid of data points. Directional displacements and strains were generated at each data point across the region of interest and for every subsequent frame in the UHS video. During this study, a total of 153 UHS videos were collected in which microbubble-induced cell displacement was sufficient to support the application of DIC and the convergence of the MatchID software, with no significant or obvious erroneous data.

### Digital Image Correlation Data Processing

To extract data relevant to the microbubble-induced displacement, processing was carried out using Python 3.12.0. The displacement data in the horizontal and vertical directions were obtained from the DIC and transformed from Cartesian to polar coordinates, with the origin at the centre of the microbubble. It was initially assumed that the displacement induced in the cell by the harmonic motion of the microbubble would also be harmonic. Therefore, for each data point across the region of interest (ROI), a Fourier transform was performed on the radial component of the deformation using the SciPy fast Fourier transform (FFT) function. The absolute magnitude of the peak at the driving frequency, 1 MHz, could then be taken as the magnitude of the deformation induced by the microbubble at each point across the ROI. By considering the angle of the complex output from the FFT ^46^, the phase spectrum for each point within the ROI could be plotted, and therefore, the phase at 1 MHz (∅_ROI_ ) of each oscillating point across the ROI could also be mapped.

### Bubble Oscillation

To determine microbubble oscillation amplitude, an edge-detection method was implemented, as previously described ^30^, to confirm the harmonic oscillation of the microbubble. The microbubble oscillations were assumed to be approximately spherical, and so the radius and centre of the microbubble could be tracked through time. Using the SciPy curve fit function, Equation 1 was fitted to the radial oscillations to find the oscillation magnitude:

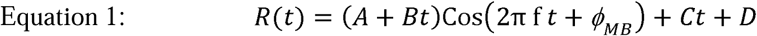

where R(t) is the radius of the microbubble at time *t*, *f* is the frequency of oscillation, *A* corresponds to the amplitude, *B* is the temporal change in amplitude, ∅_MB_ , the phase of microbubble oscillation, *C* the temporal offset, and *D* the absolute offset.

### Fitting Data to a Regression Model

To determine and quantify the mechanical decay of microbubble-induced displacement as a function of distance from the microbubble, data collected from UHS videos were fit to three regression models. An exponential function (Equation 2) was examined, as it has been demonstrated to fit the attenuation of a mechanical wave in other viscoelastic materials and also in biological cells ^2,32^. An inverse-square function (Equation 3) was also examined, as it describes the contact between two elastic bodies and the resulting deformation pattern in the well-studied Hertzian contact model.

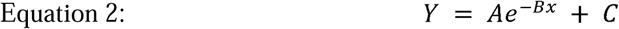

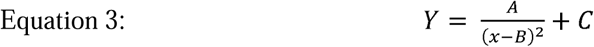

In equations (2) and (3), Y represents the absolute displacement of the cell at a specific distance from the microbubble boundary, *x* represents the radial distance from the microbubble boundary, and *A*, *B* and *C* are coefficients to be determined. By calculating the R^2^ value for each decay fit function, it was found that exponential decay best represented the data. Using the regression models, descriptors could be applied to compare the difference in displacement responses between interactions. This was defined as the distance from the bubble boundary at which the displacement is half that at the bubble/cell boundary. This is called the ‘half decay distance’, defined by the following equation:

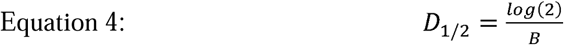

where *D_1/2_* represents the half decay distance and *B* the exponential decay coefficient found from Equation 2.

### Propagation Speed using Phase Change

After calculating the corresponding phase from the FFT, the phase change with distance from the microbubble boundary represents a temporal delay deformation through the cell. To account for any asymmetries in the microbubbles’ oscillation and to account for variations in cell morphology and heterogeneity, a narrow radial slice of the ROI was selected where there was an evident gradual decay in ∅*_ROI_* with distance from the microbubble. A linear regression was then performed on ∅*_ROI_* with distance *x* from the microbubble boundary and the gradient was then used to determine the propagation speed using Equation 5.

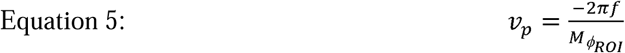

where *V_p_* is the wave speed in the cell, f is the driving frequency of 1 MHz, and M_∅ROI_ is the spatial gradient of the phase provided by the linear regression (A complete derivation can be found in Supplementary Information 2).

### Assessment of microbubble-induced cell permeabilisation

MG63 cells were cultured on a 35 mm glass Ibidi cell culture dish (Thistle Scientific, Glasgow, UK) seeded at 40,000 cells/cm^2^, 24 hours ahead of time. Cells were treated as previously described with calyculin A, nocodazole, cytochalasin D, latrunculin A, blebbestatin, and Y-27632, each with their own corresponding untreated control to account for minor variations in microbubble batch production. Propidium iodide (PI) was used as a marker for successful microbubble-induced permeabilisation (Fisher Scientific, Leicestershire, UK). An acoustically compatible PDMS lid ^47^ was then attached to the cell culture dish and filled with 10 mL of DMEM, 10 μL of 4 mg/mL PI, and 100 μL of microbubble suspension (∼1×10^7^ microbubbles per mL), resulting in a final concentration of 4 μg/mL of PI. The dish was then inverted, allowing the microbubbles to float in contact with the cell culture area. The dish was then suspended in a 0.3 L water bath, and was ultrasonically stimulated at a frequency of 1 MHz, a peak acoustic pressure of 300 kPa, a 30% duty cycle, and a 1000 kHz pulse repetition frequency using a 0.9 MHz focused transducer (Precision Acoustics, Dorchester, UK), which was driven by the same power amplifier fed from a signal generator (Aim-TTi, Cambridgeshire, UK), and was monitored by the HS3 Handyscope oscilloscope (TiePie, Sneek, The Netherlands). Cells were then removed from the water bath and washed 3× with PBS before being imaged.

### Fluorescent Microscopy

To determine the number of cells permeabilised as a proportion of total cells, phase contrast and fluorescence imaging was performed using a Zeiss Observer inverted microscope (Zeiss AG, Oberkochen, Germany) at 10× magnification using the Cy5 channel (ex λ = 628 – 648 nm, em λ = 673 – 712 nm) for imaging PI uptake. The number of cells taking up PI and the total number of cells in each image were counted using ImageJ and were taken as a percentage for comparison.

### Confocal Microscopy

For assessing the cytoskeletal structure, MG63 cells were fixed with 4% (w/v) PFA for 20 minutes at room temperature and permeabilised with 0.5% (v/v) Triton X-100 in PBS for 20 minutes, after which they were washed and incubated with 1 µg/mL Alexa Flour 488 phalloidin (Fisher Scientific, Leicestershire, UK) for 30 minutes at room temperature. Cells were then washed and stained with 4′,6-diamidino-2-phenylindole (DAPI, Fisher Scientific, Leicestershire, UK) at room temperature for 5 minutes (1:1000). Confocal images were acquired using a Leica TCS SP5 confocal microscope (Leica Microsystems, Milton Keynes, UK). Z-stacks were collected at 0.5 µm intervals using a 63× objective. DAPI and Alexa Fluor 488 phalloidin were imaged using excitation wavelengths of λ = 405 nm and λ = 488 nm, respectively.

### Statistical Analysis

All fitting data functions were described using the coefficient of determination, R^2^. Subsequently, statistical analysis was performed using GraphPad Prism 8.3 (GraphPad Software, La Jolla, CA, USA). Figure 2E uses a Wilcoxon matched-pairs test. Figure 3C and 4C uses a one-way ANOVA with Tukey’s multiple comparison test. Figures 3F use a one-way ANOVA with Dunnett’s post hoc test. Figure 3H use a two-way ANOVA with Fisher’s LSD post hoc test.

## SUPPLEMENTARY MATERIALS

Video files 1-4 have been submitted in addition to the manuscript

**Supplementary Figure 1:**
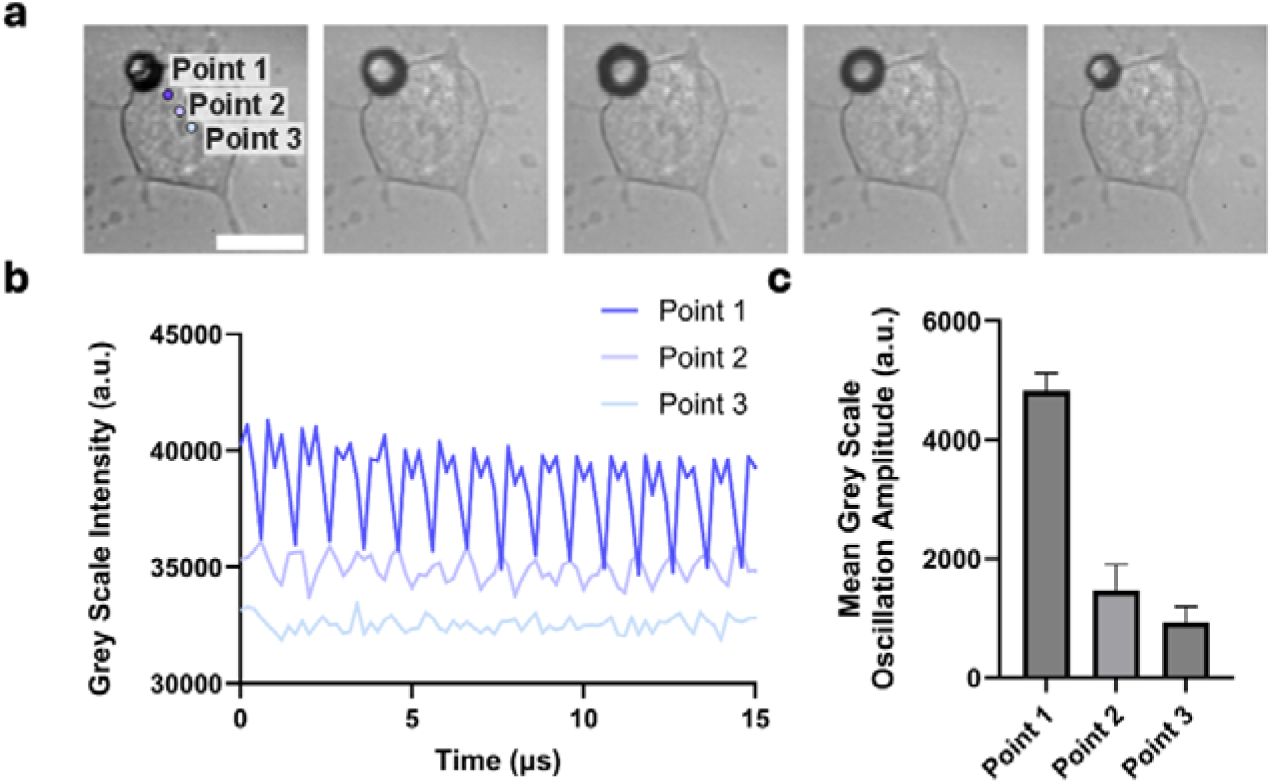
Grey scale oscillations. Figure 1: Using fluctuations in the grey scale from the high-speed imaging to infer possible deformation in the cell cytoplasm. **a)** *5 frames from* Video 1 showing ultrasound microbubble-induced displacement on the cell. Points 1, 2 and 3 have been selected at increasing distance, and b) the oscillation in the grey scale values through the first 10 μs is shown. c) The mean amplitude of these oscillations is then plotted for each point, showing a clear decrease with distance from the microbubble.

**Supplementary Figure 2:**
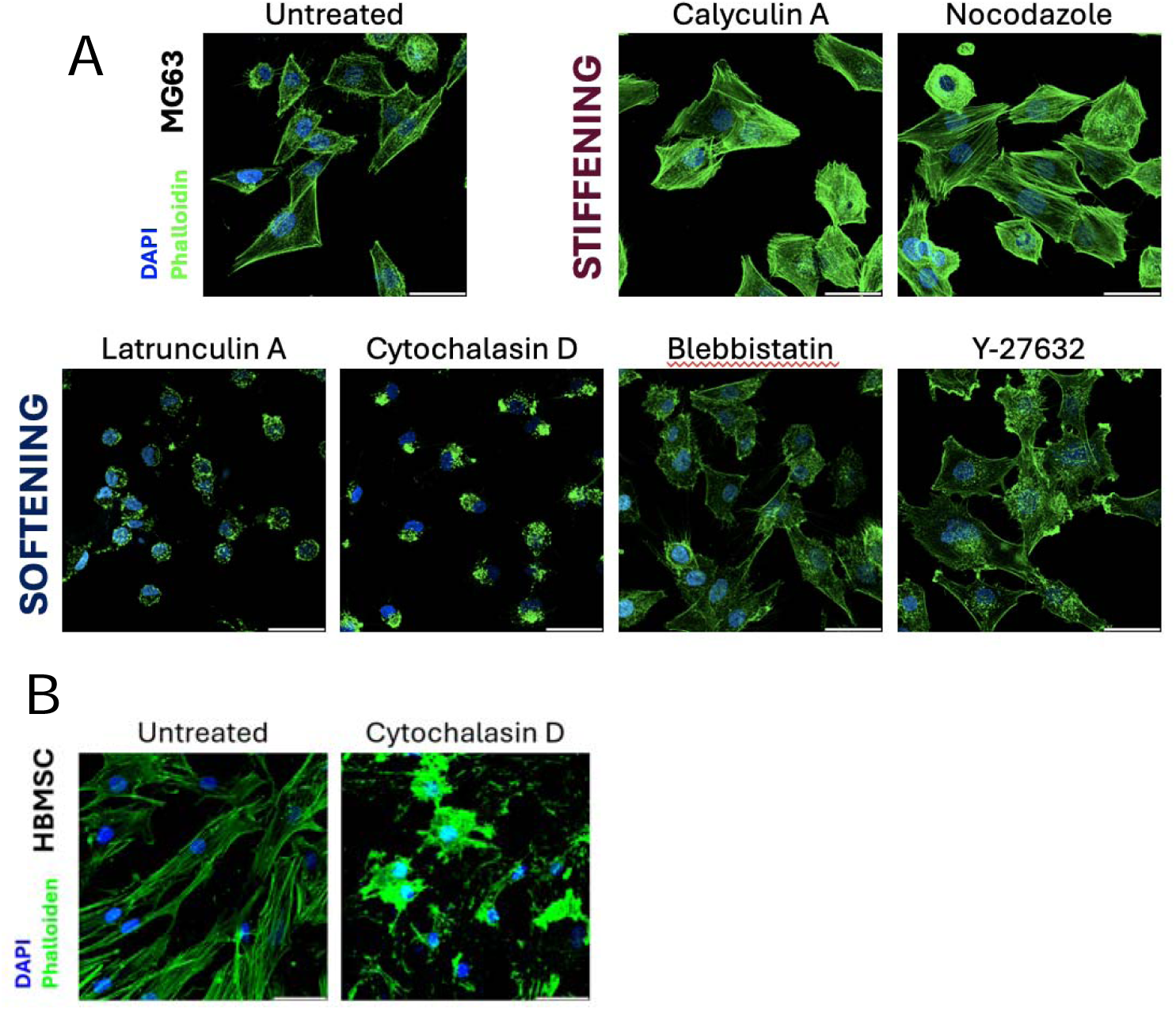
Additional images of cells treated with cytoskeletal disrupting agents. Figure 2: Confocal images with DAPI-stained nuclei (blue) and phalloidin-stained actin (green) for untreated, calyculin or nocodazole-treated (stiffening agents) and latrunculin A, cytochalasin D, blebbistatin, or Y-27632-treated (softening agents) in (A) MG63 cells and (B) BMSCs (scale bar = 50 µm).

**Supplementary Figure 3:**
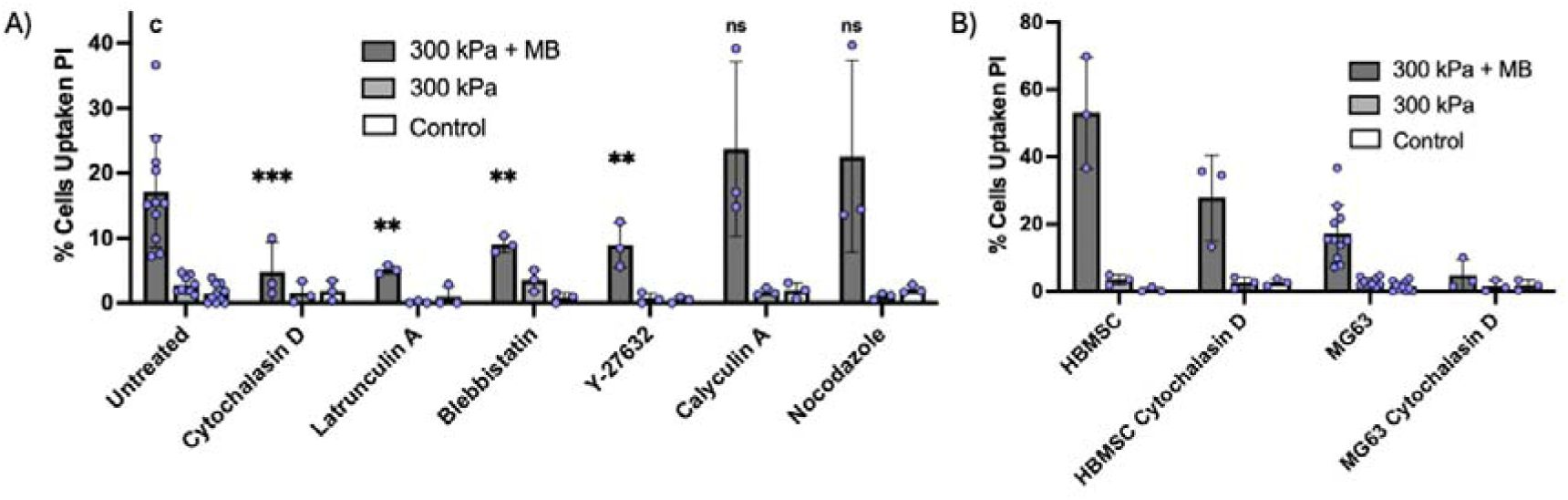
Sonopermeablisation data with controls (Figure 3f & h) Figure 3: Sonopermeabilisation data showing % PI uptake in; a) untreated MG63 cells or treated with cytochalasin D, latrunculin A, blebbistatin, Y-27632, calyculin A, and nocodazole and b) HBMSCs compared with MG63 cells untreated and treated with cytochalasin D. Both figures contain microbubble free (300 kPa) and a microbubble and ultrasound-free (control) group.

**Supplementary Table 1:**
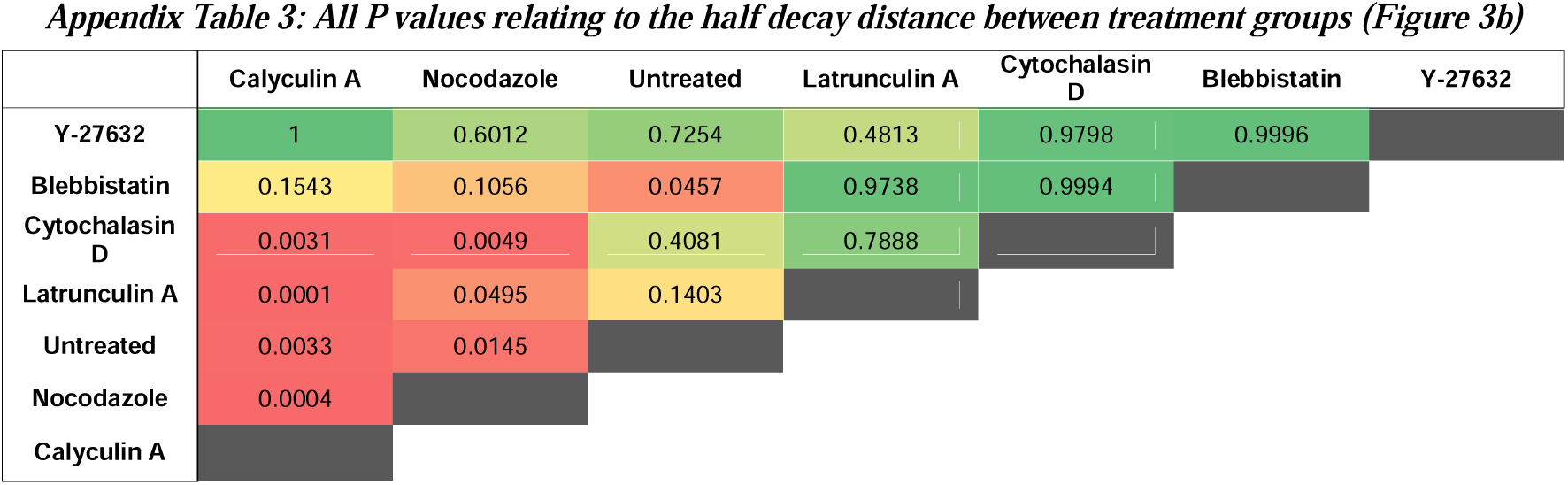
Statistical analysis of the effect of cytoskeletal inhibitors on D1/2.

**Supplementary Table 2:**
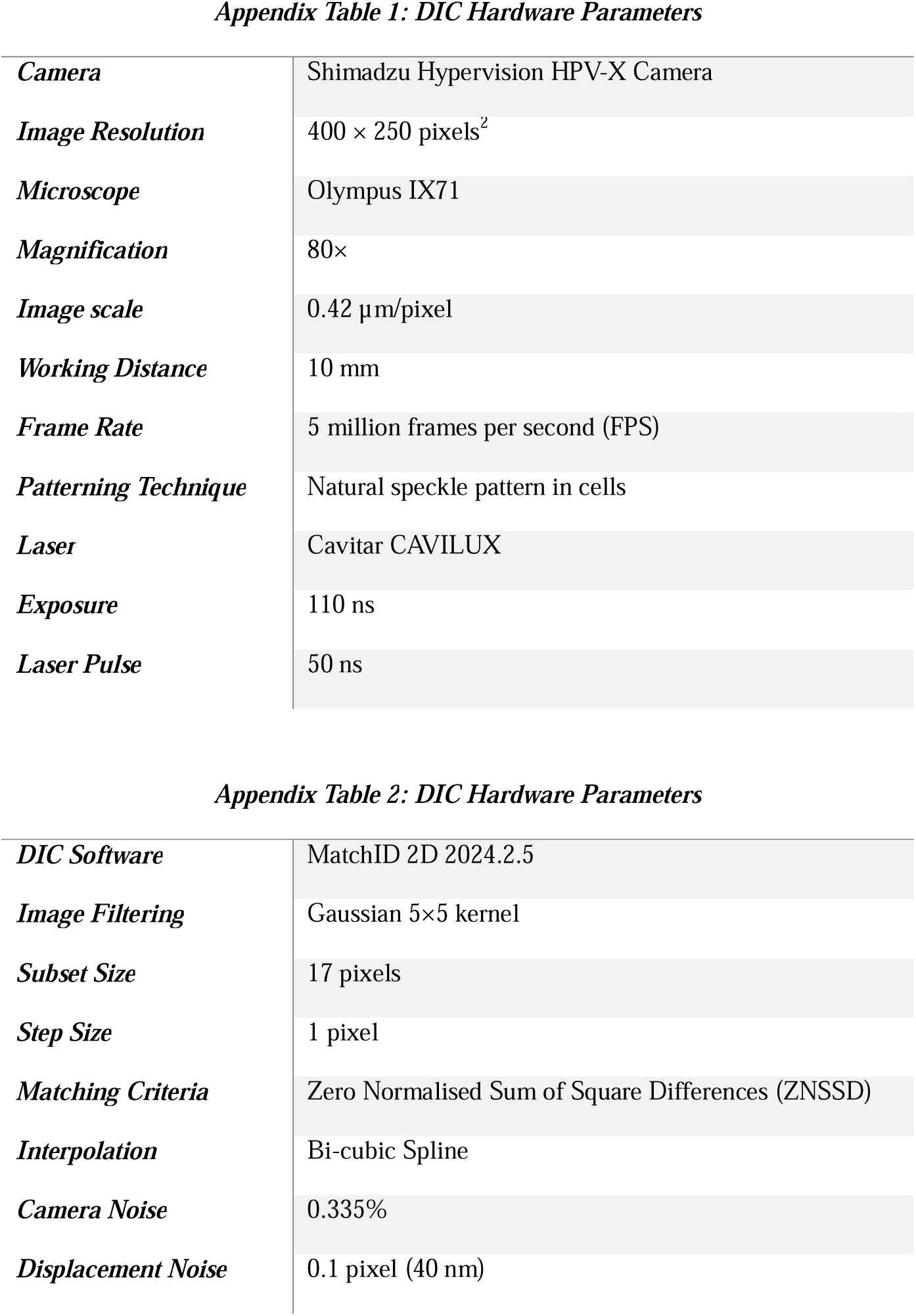
MatchID Hardware and Software Tables.

Supplementary Information 1: All data can be found in zipped and indexed folder

Supplementary Information 2: Wave Speed Derivation

Consider the harmonic oscillation;

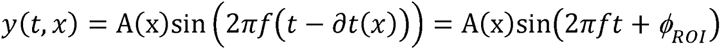

where *y*(*x, t*) is the deformation of a point at a distance *x* from the microbubble and at time *t*, *A*(*x*) is the amplitude of the oscillation at a point at a distance *x* from the microbubble, *f* is the driving frequency in Hz, *∂t*(*x*) is the lag time of a point a distance *x* from the microbubble and *∅_ROI_* (*x*) = -2*πf∂t*(*x*) is the phase change, in radians, of a point a distance *x* from the microbubble. Therefore;

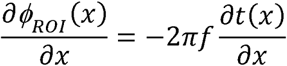

or;

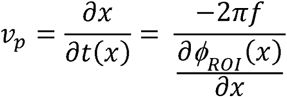

Assuming linearity, *∅_ROI_* (*x*) = *M_∅RO_ x*+ *C*, where *M_∅ROI_* is the fitted gradient of a linear regression and *C* is a constant. You therefore find, 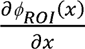=*M_∅ROI_* , and Equation 5:

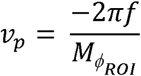

