## Supplementary Information 1 for "Cell material state determines high-frequency cell deformation and microbubble-induced permeabilization": readme.txt

This is the readme file for the folder Supplementary Information 1 accompanying the manuscript "The material properties of the cell determine microbubble-induced cell deformation and permeabilization" by Sloan, Pattinson et al. 

This folder contains (1) videos for all cell-microbubble interactions described in the results, and (2) analysis of cell absolute displacement using DIC for each of the interactions and (3) analysis of cell wave speed using DIC for each of the interactions 

The folders are titled with the name of the treatment first (Blebbistatin, Calyculin A, Cytochalasin D, Latruculin A, Nocodazole, Untreated or Y-27632) and the file type second ('Videos', 'displacement analysis' or 'wave speed'). 

Within each Video folder, the videos are provided as AVI files, with the raw file names of the interactions preserved. 
Within each 'displacement analysis' folder, the files are provided as PNG files, with the file names corresponding directly to the video file in the corresponding folder with the suffix "_Magnitude" for each file.
Within each 'wave speed' folder, the files are provided as PNG files, with the file names corresponding directly to the video file in the corresponding folder with the suffix "_Wave_Speed" for each file.
