## Supplementary figures and images for "Cell material state determines high-frequency cell deformation and microbubble-induced permeabilization"

### Camera_13_44_00_Magnitude.png

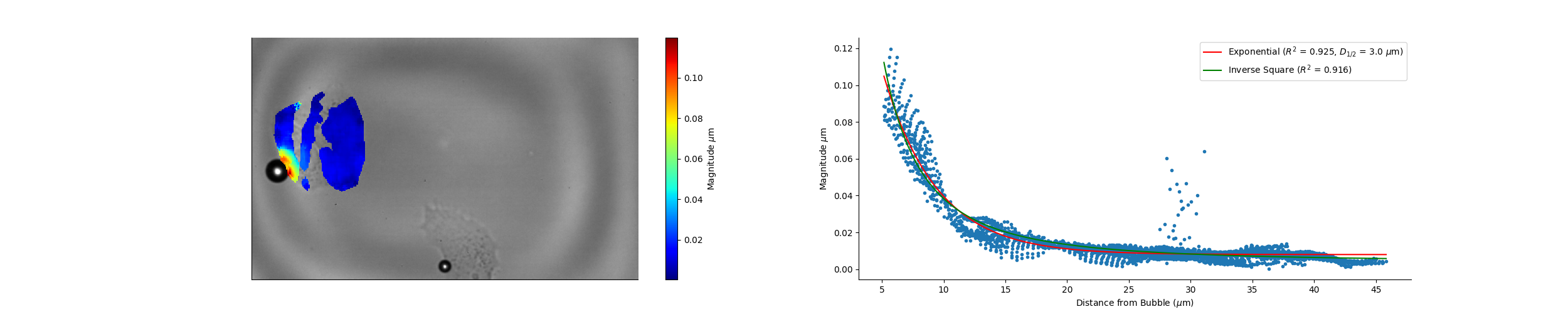

### Camera_13_44_00_Wave_Speed.png

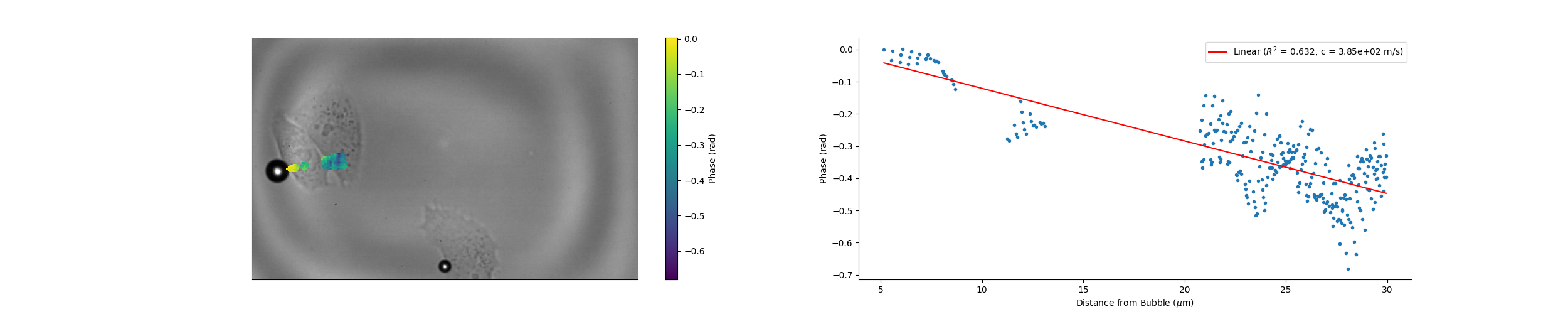

### Camera_13_44_55_Magnitude.png

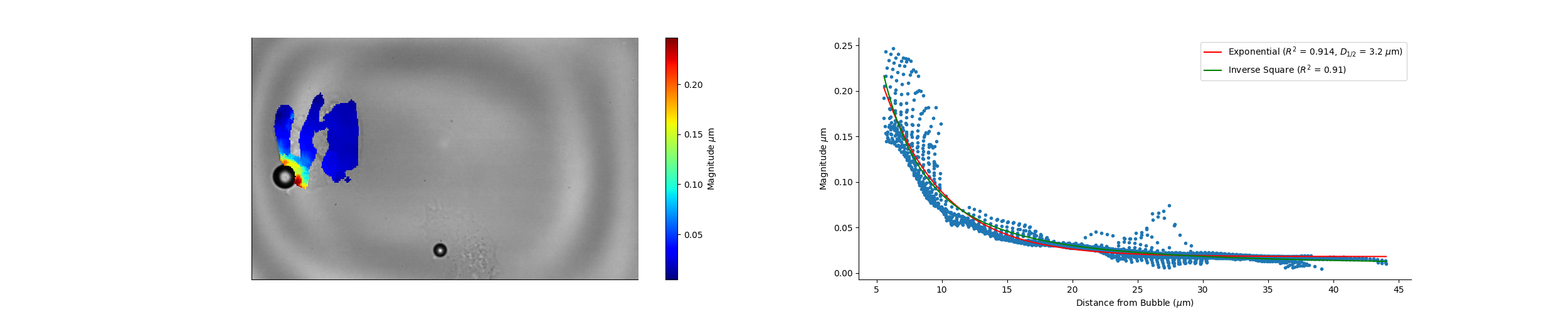

### Camera_13_44_55_Wave_Speed.png

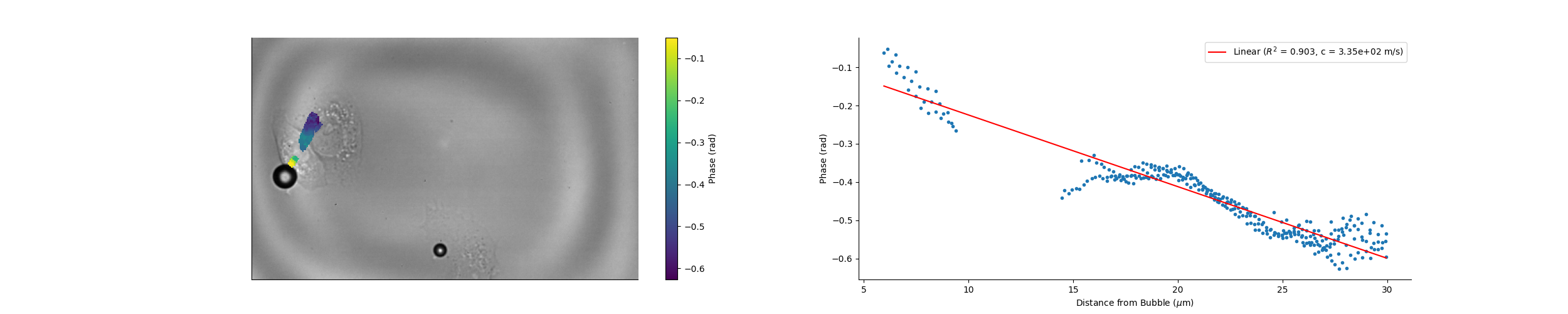

### Camera_13_48_23_Magnitude.png

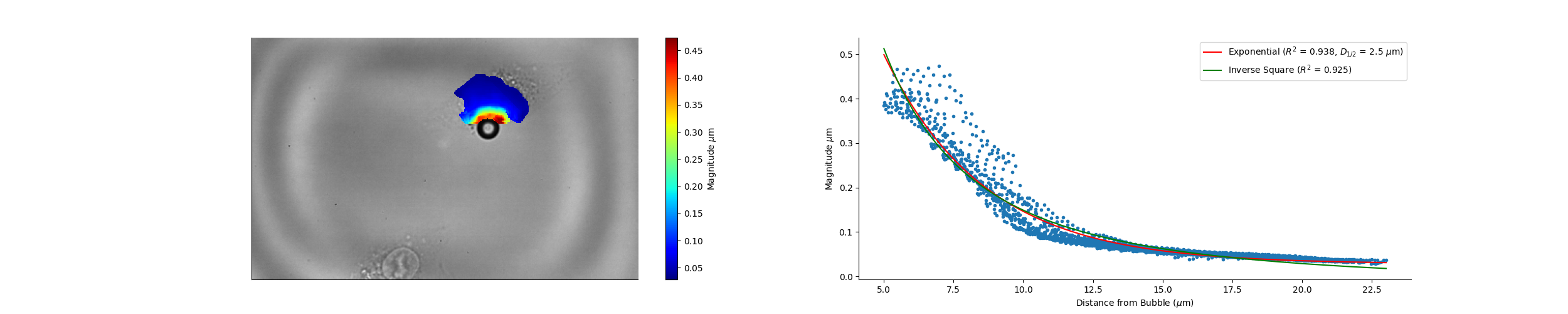

### Camera_14_07_28_Magnitude.png

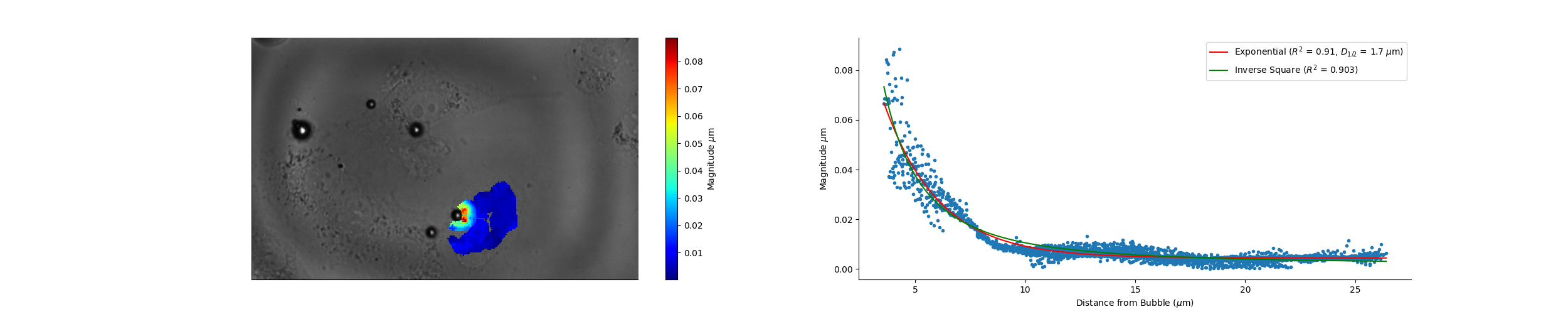

### Camera_14_07_28_Wave_Speed.png

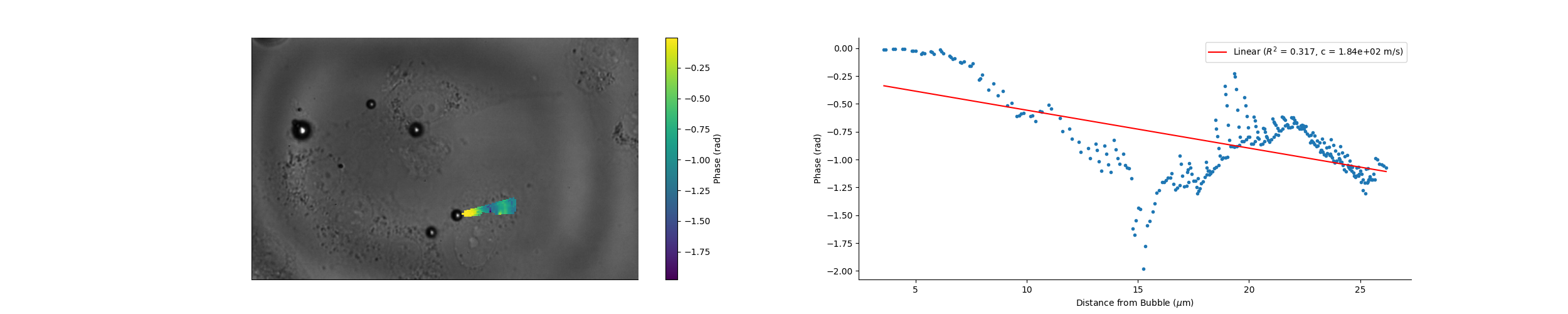

### Camera_14_16_07_Magnitude.png

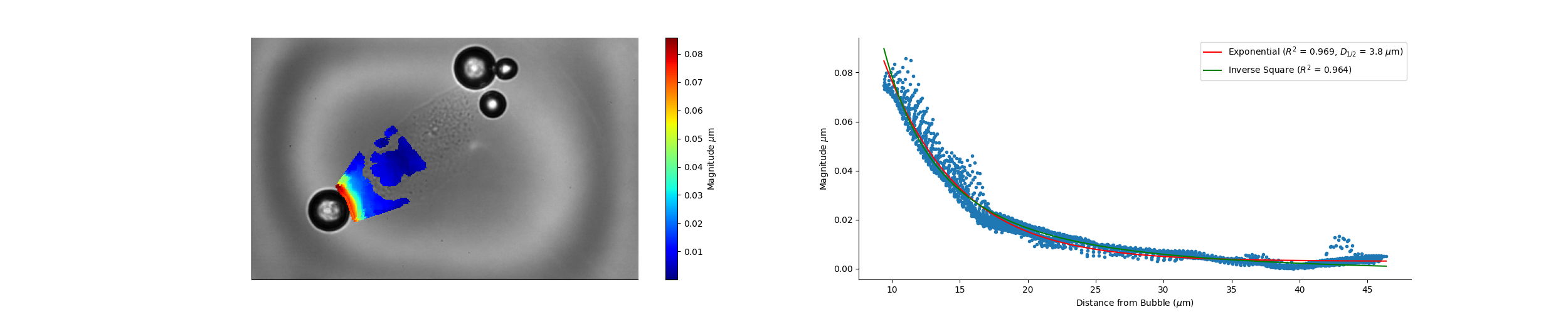

### Camera_14_16_07_Wave_Speed.png

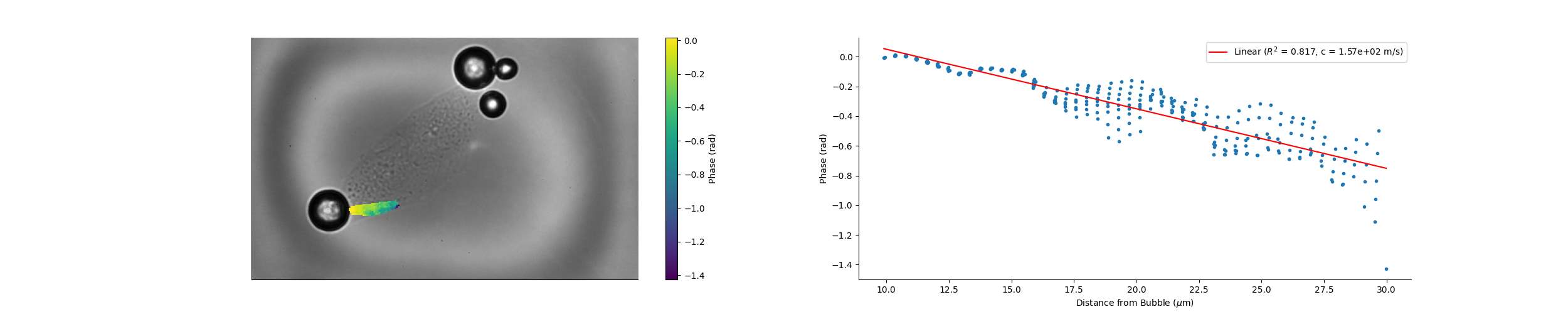

### Camera_15_00_36_Magnitude.png

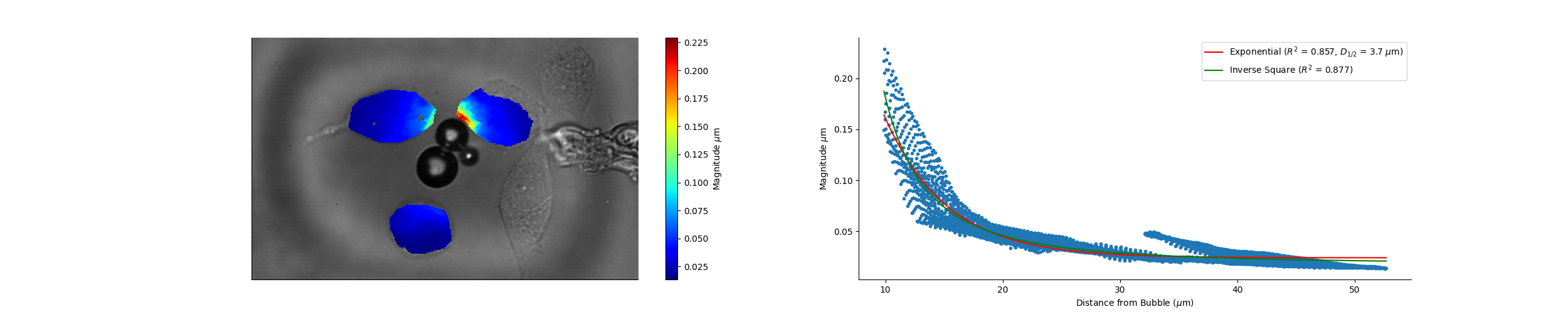

### Camera_15_00_36_Wave_Speed.png

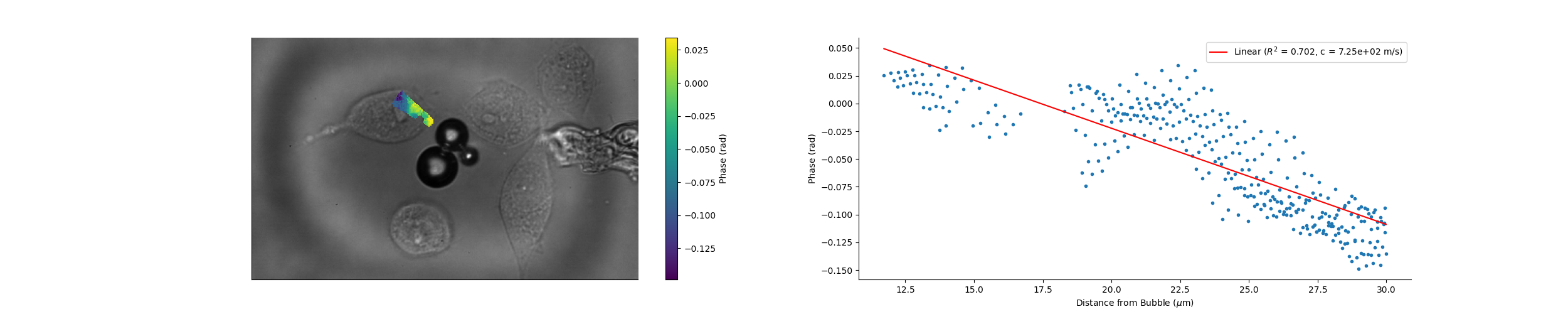

### Camera_15_04_05_Magnitude.png

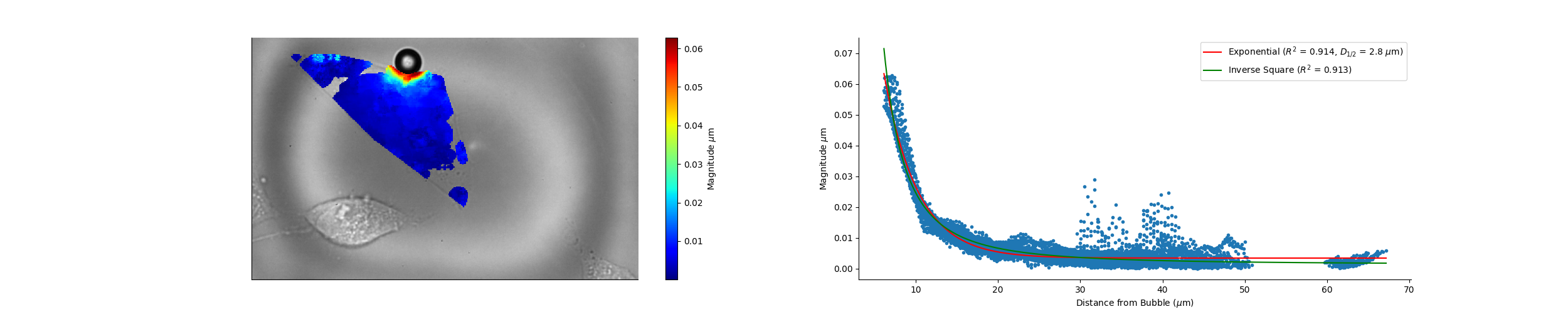

### Camera_15_04_05_Wave_Speed.png

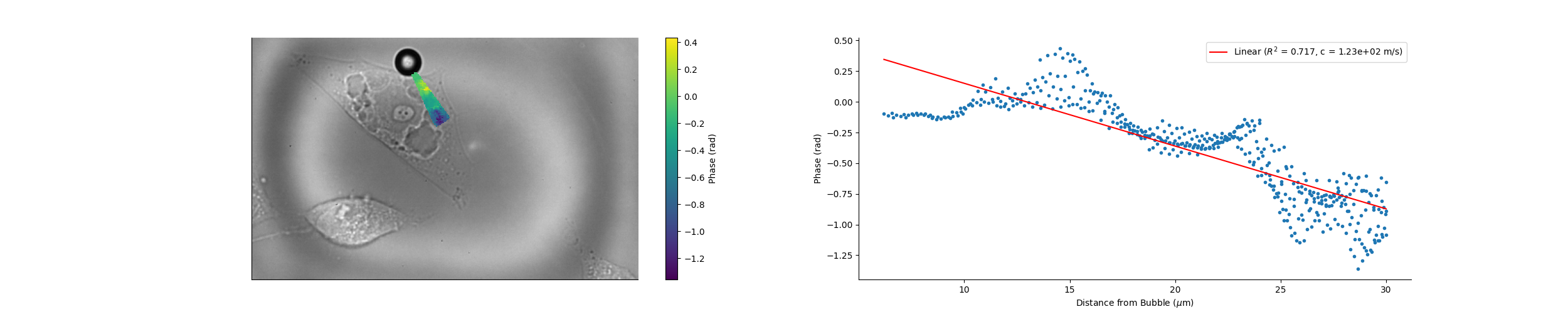

### Camera_15_24_33_Magnitude.png

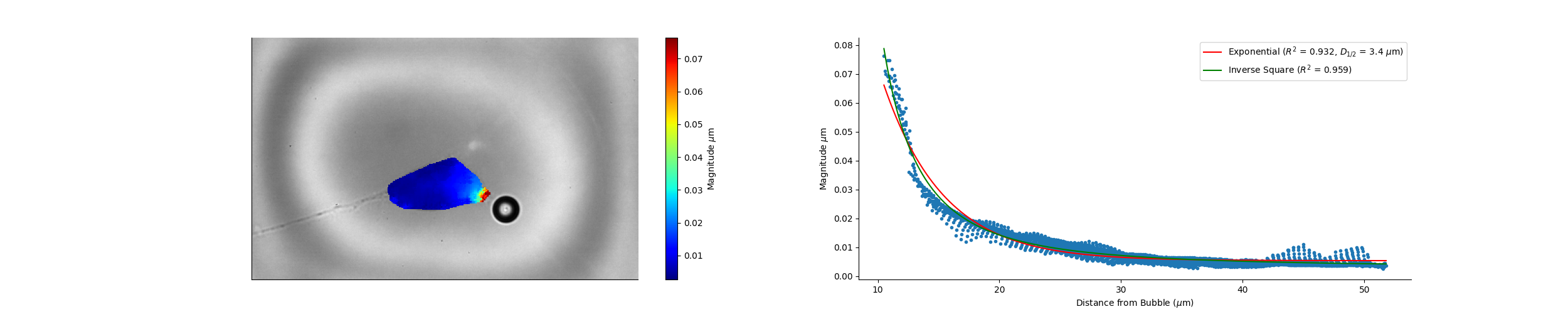

### Camera_15_24_33_Wave_Speed.png

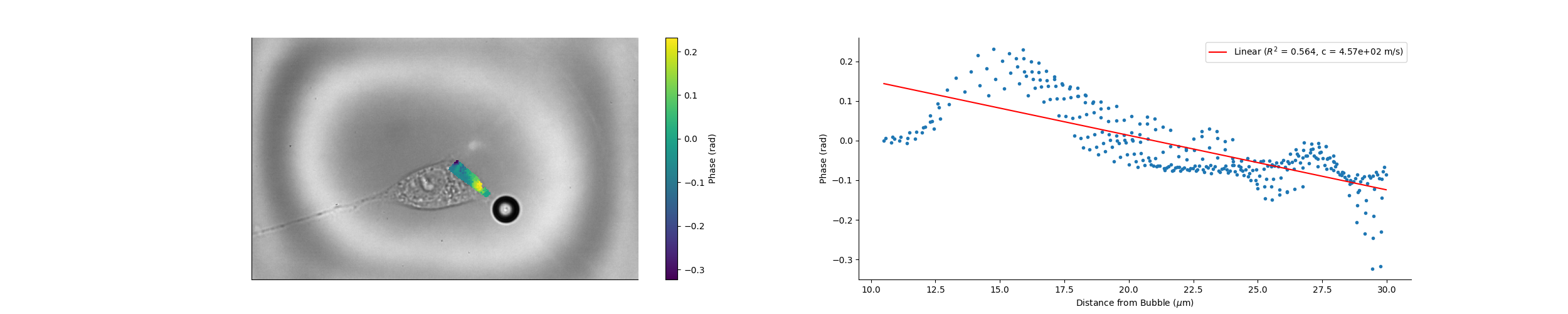

### Camera_15_27_40_Magnitude.png

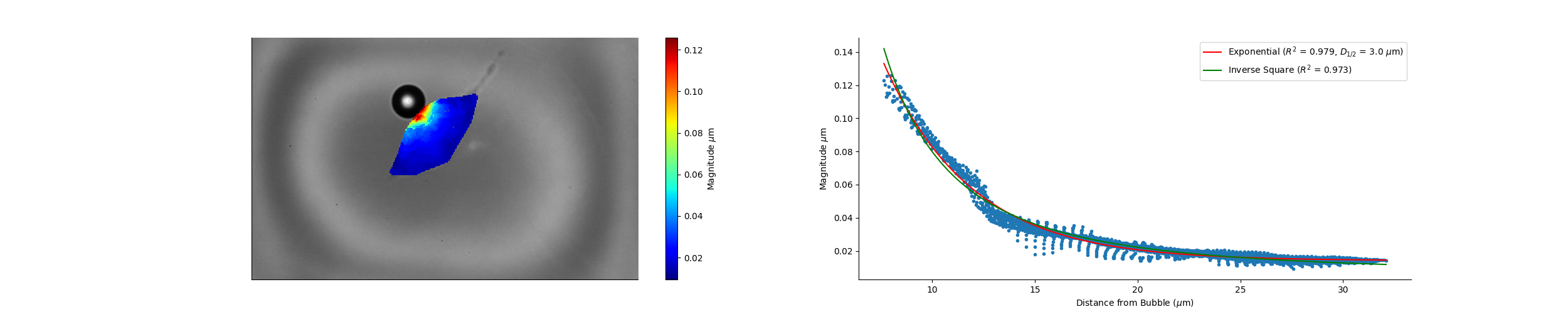

### Camera_15_27_40_Wave_Speed.png

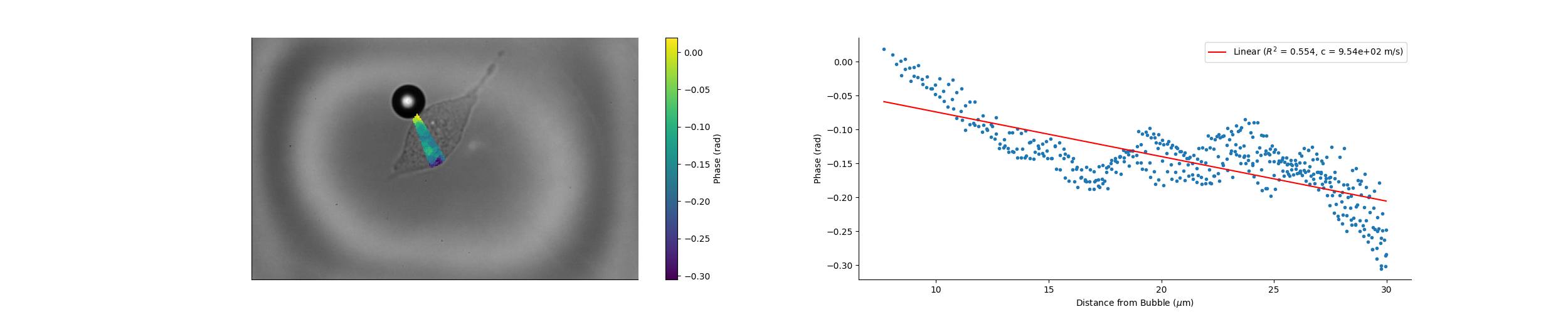

### Camera_15_43_02_Magnitude.png

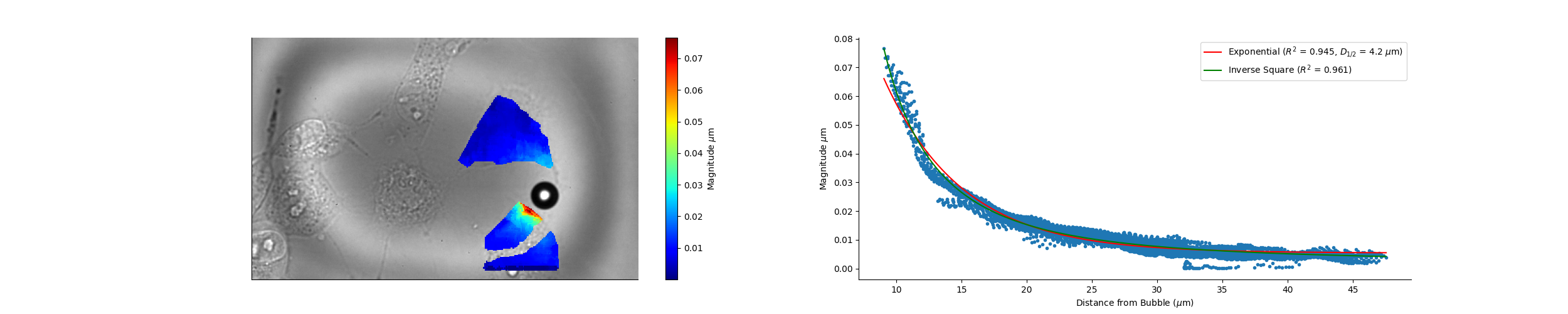

### Camera_15_43_02_Wave_Speed.png

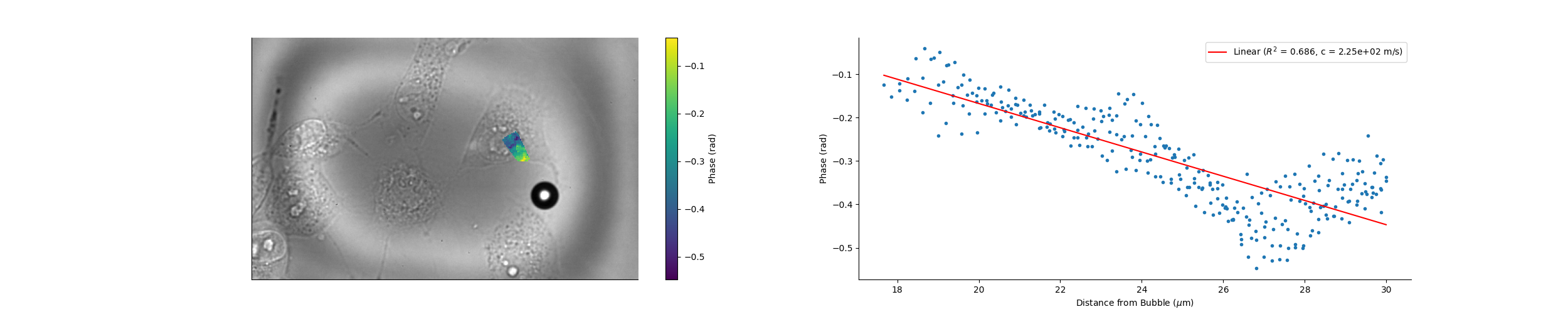

### Camera_15_50_28_Magnitude.png

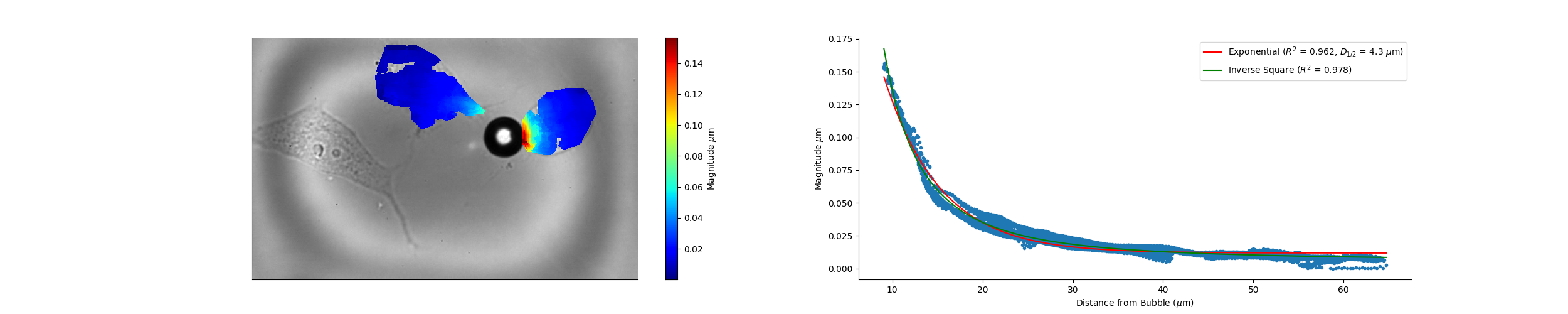

### Camera_15_50_28_Wave_Speed.png

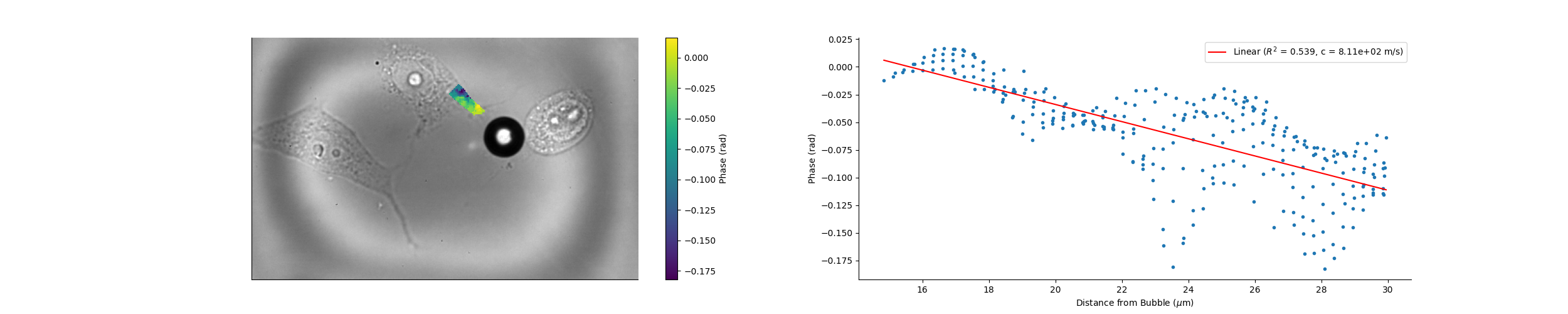

### Camera_15_51_43_Magnitude.png

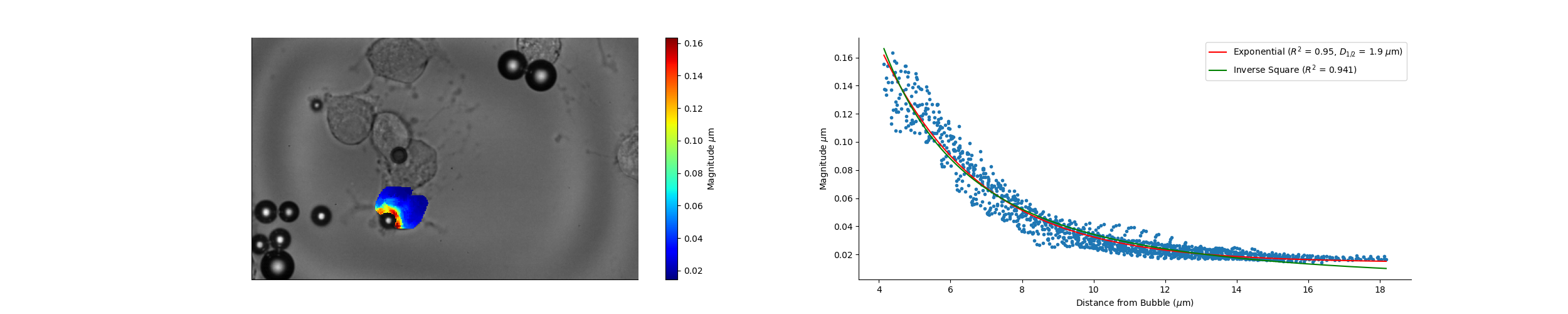

### Camera_15_56_35_Magnitude.png

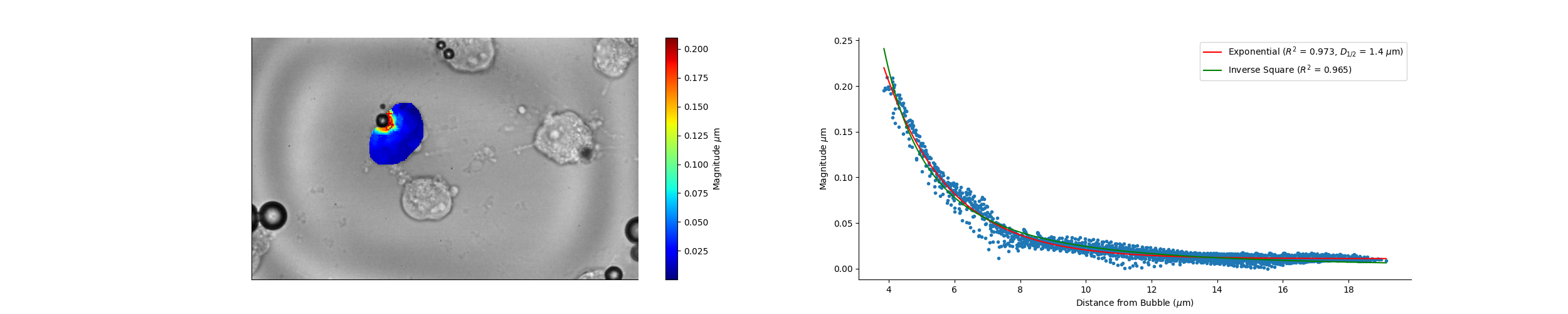

### Camera_15_58_22_Magnitude.png

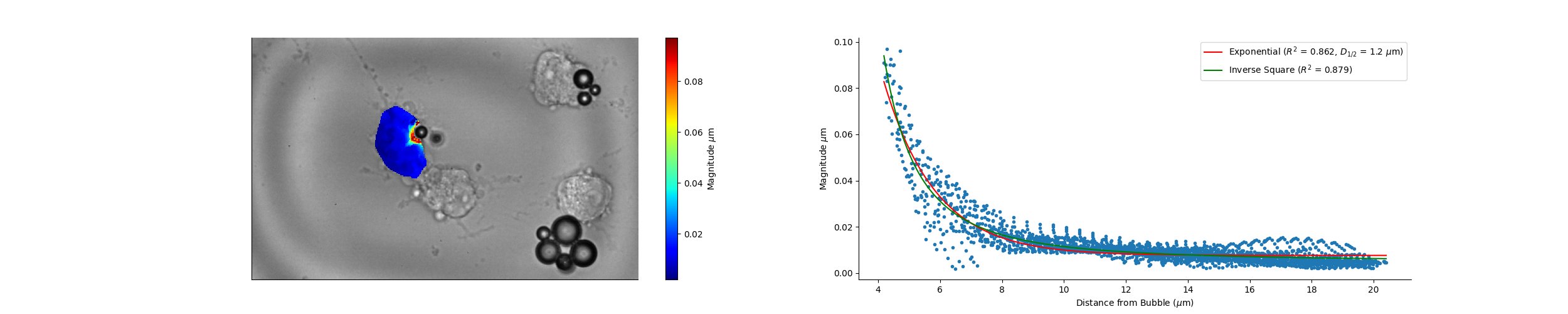

### Camera_16_02_12_Wave_Speed.png

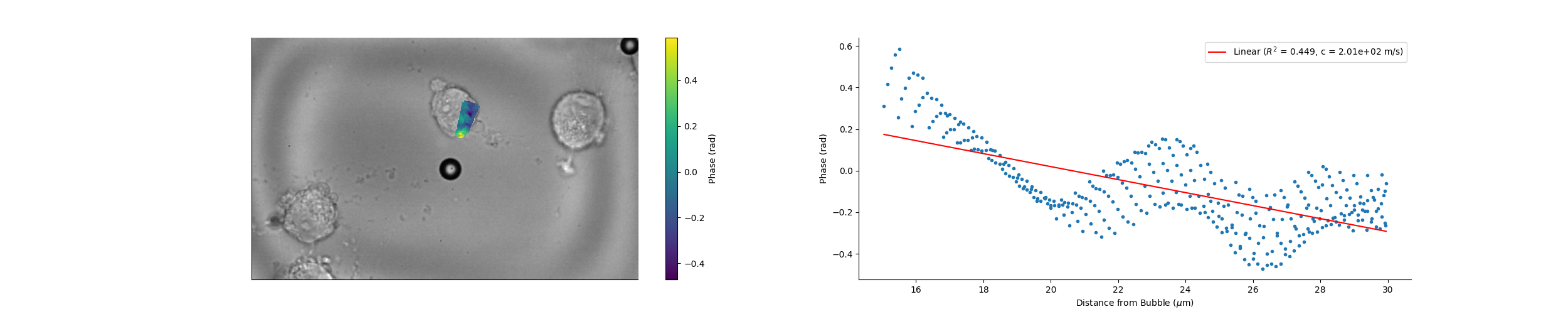

### Camera_16_08_07_Magnitude.png

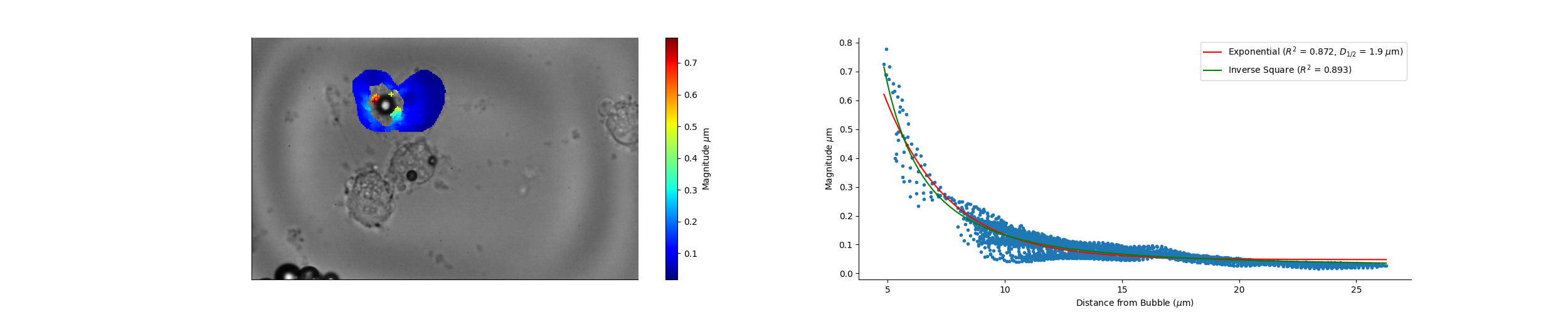

### Camera_16_08_07_Wave_Speed.png

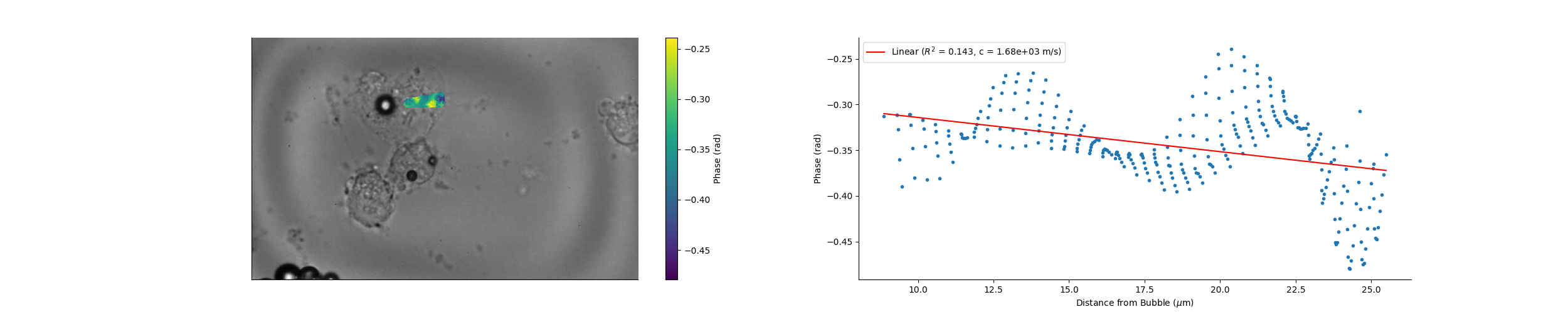

### Camera_16_13_44_Magnitude.png

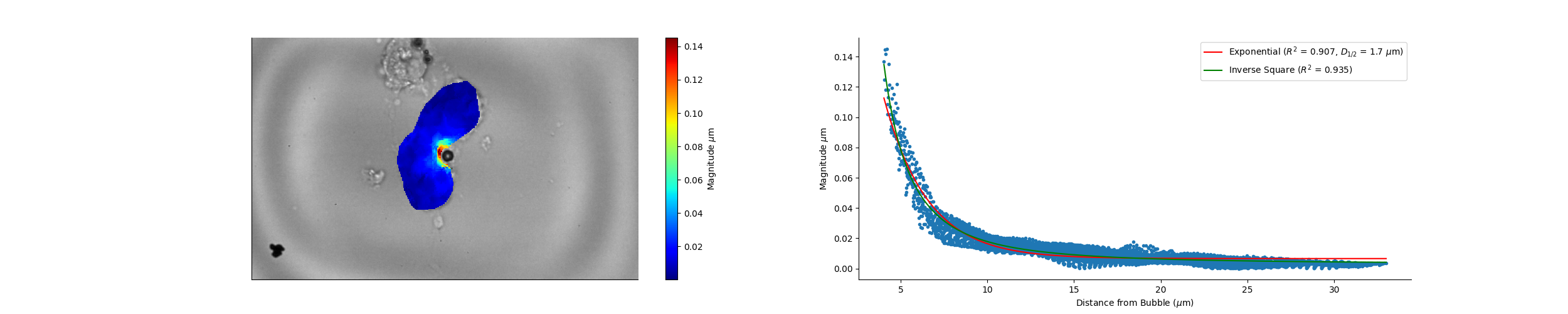

### Camera_16_13_44_Wave_Speed.png

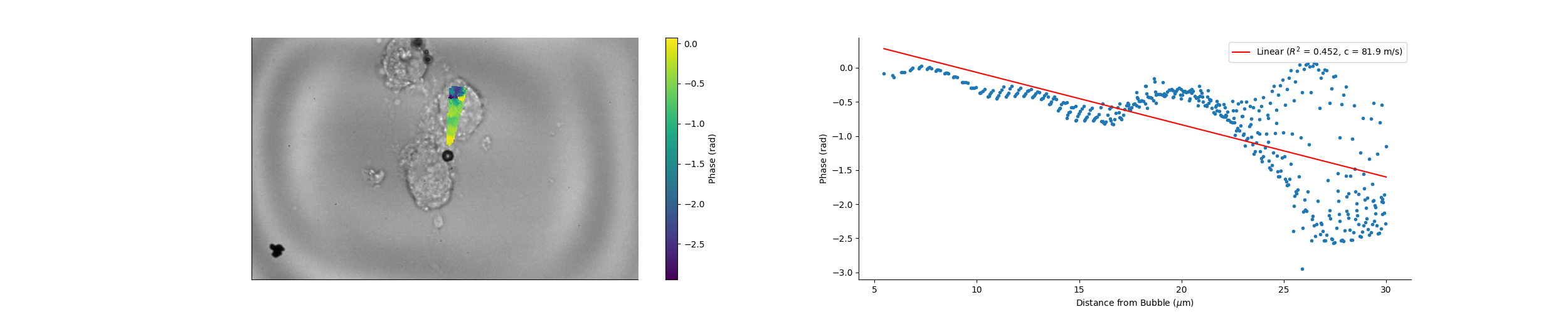

### Camera_16_15_19_Magnitude.png

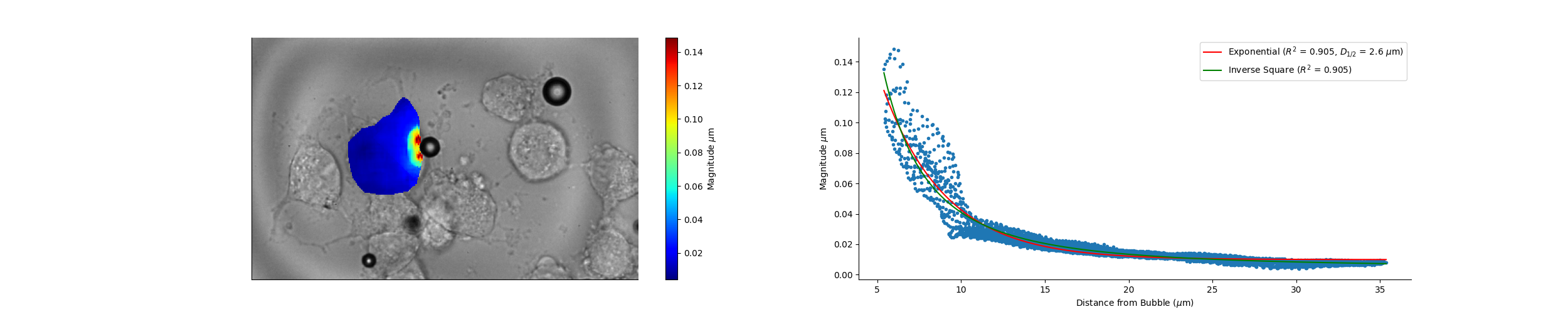

### Camera_16_15_19_Wave_Speed.png

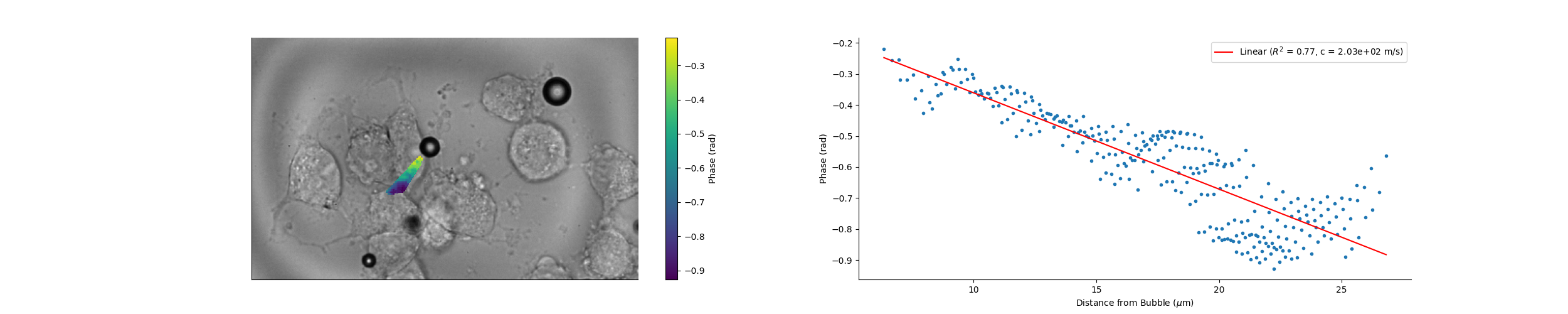

### Camera_16_18_02_Magnitude.png

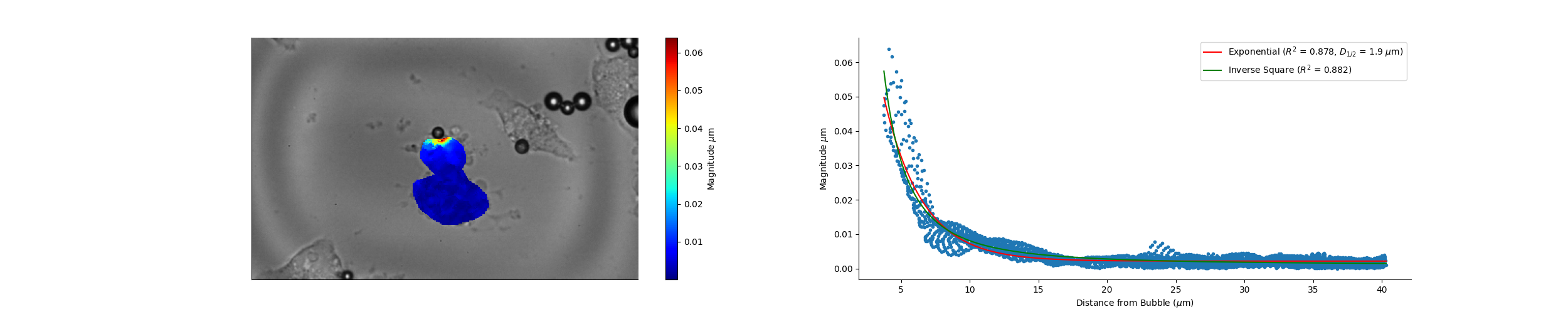

### Camera_16_18_02_Wave_Speed.png

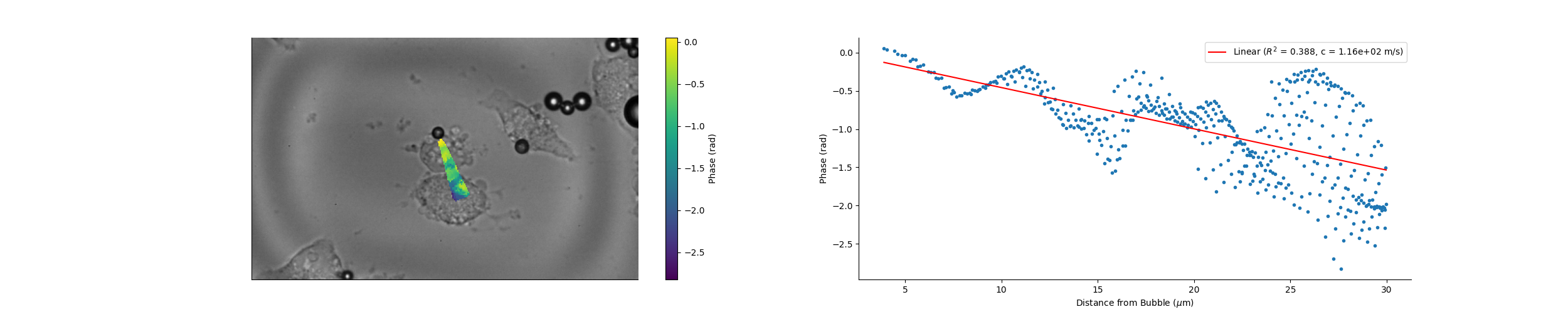

### Camera_16_20_04_Magnitude.png

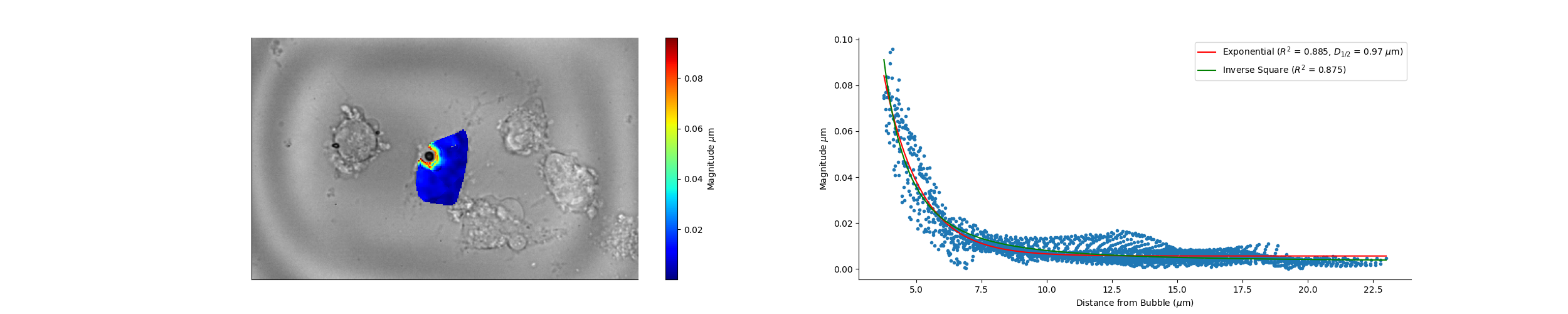

### Camera_16_27_53_Magnitude.png

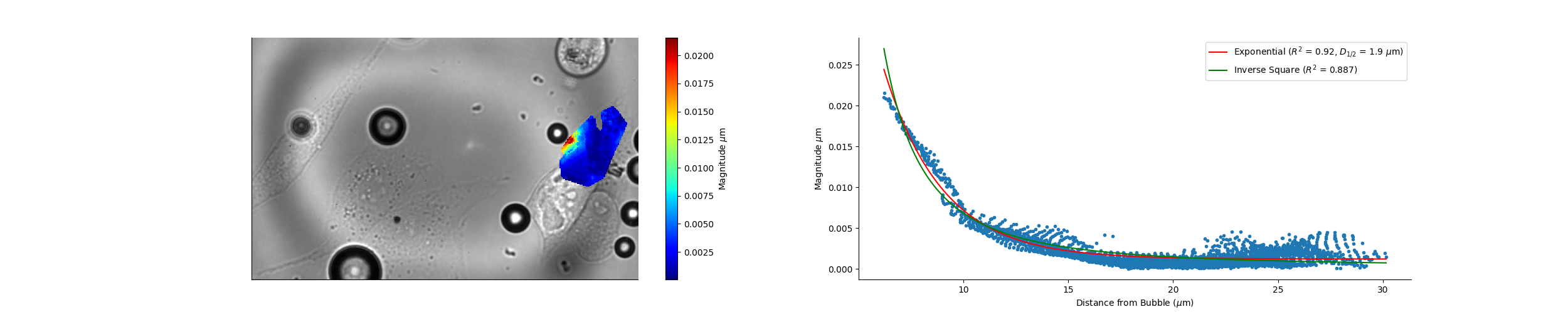

### Camera_16_27_53_Wave_Speed.png

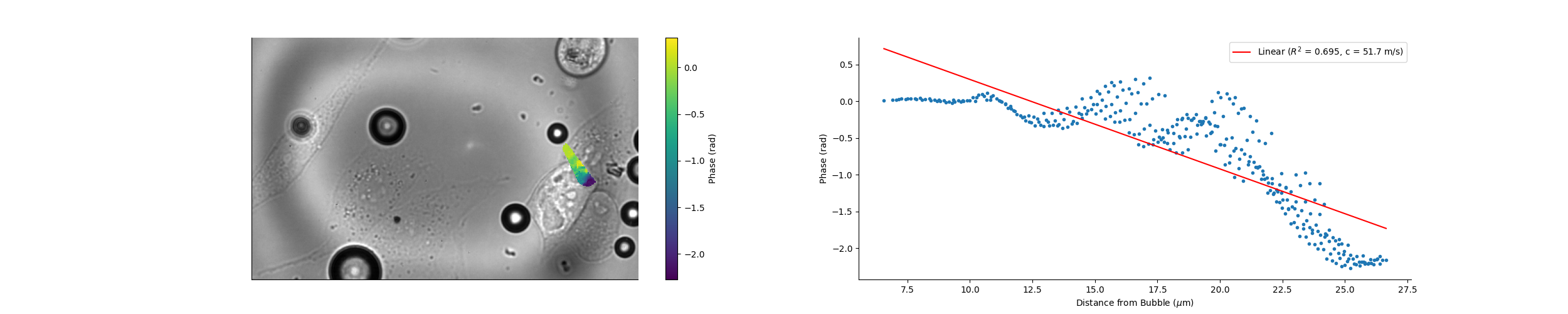

### Camera_16_35_20_Magnitude.png

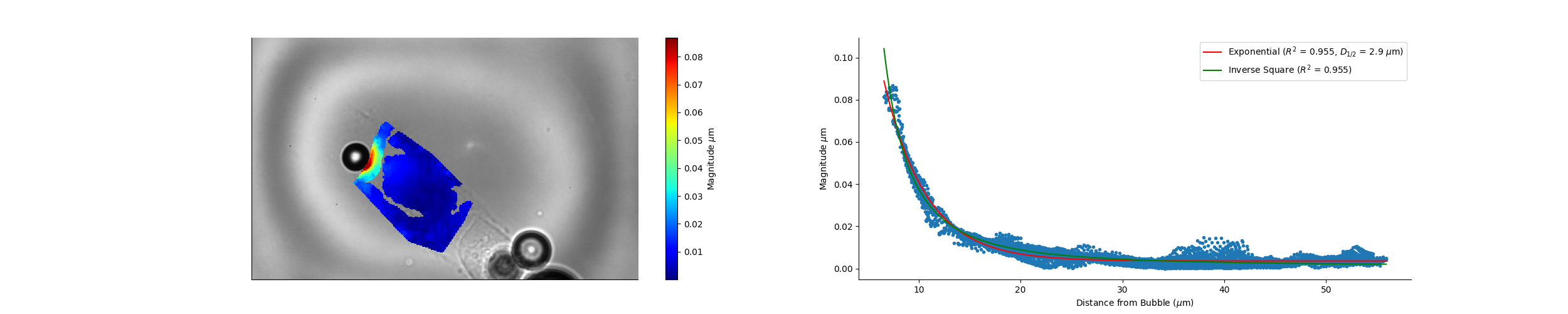

### Camera_16_35_20_Wave_Speed.png

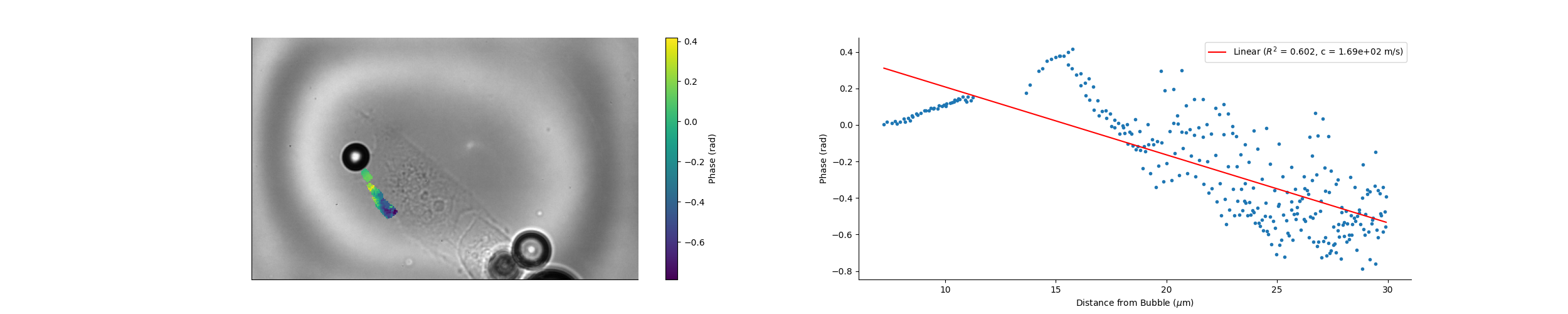

### Camera_16_36_40_Wave_Speed.png

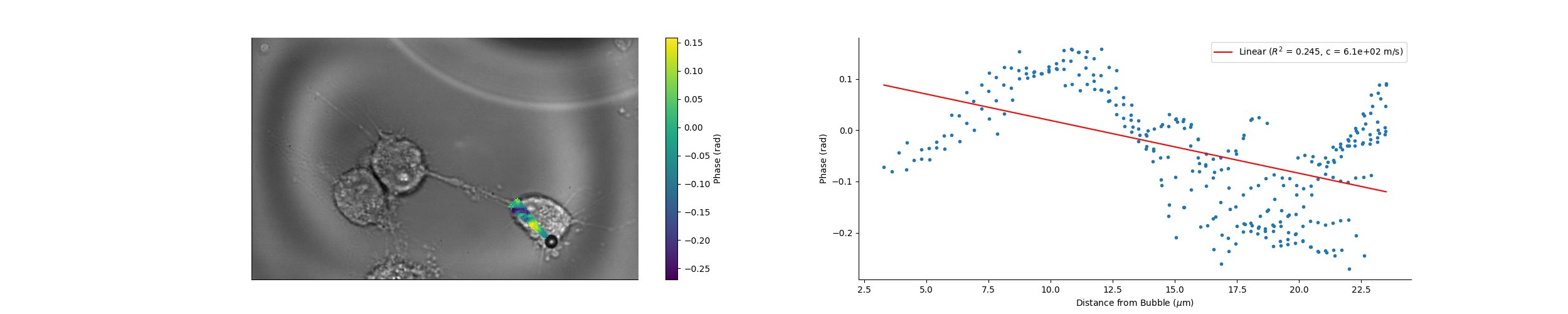

### Camera_16_38_18_Wave_Speed.png

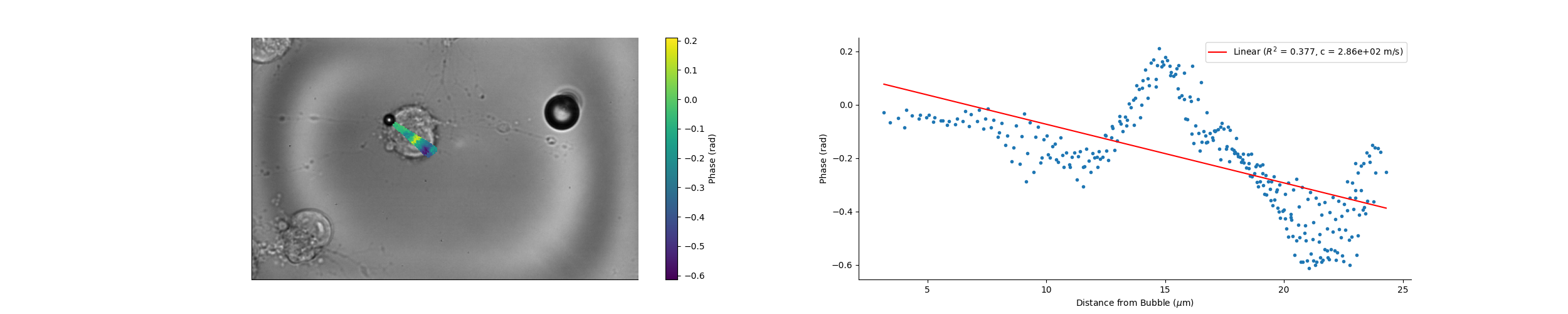

### Camera_16_38_20_Wave_Speed.png

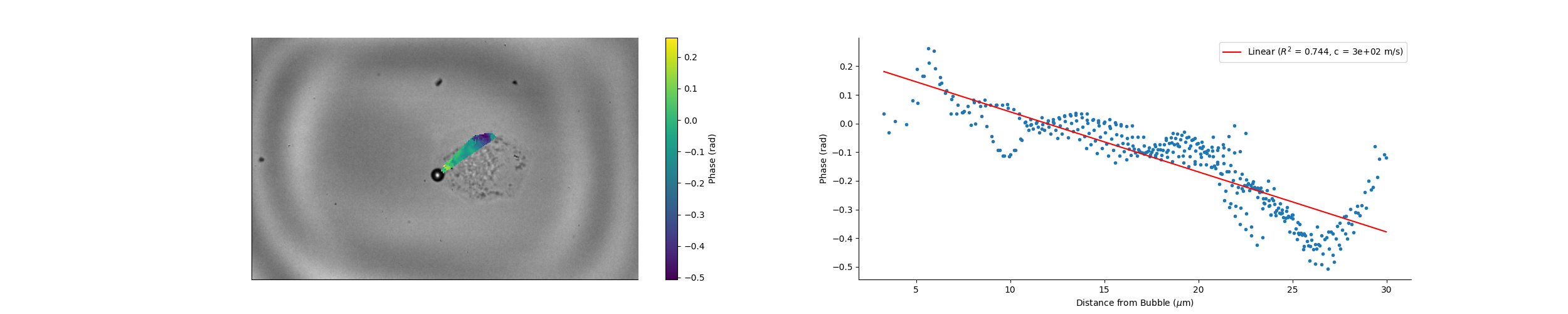

### Camera_16_41_20_Magnitude.png

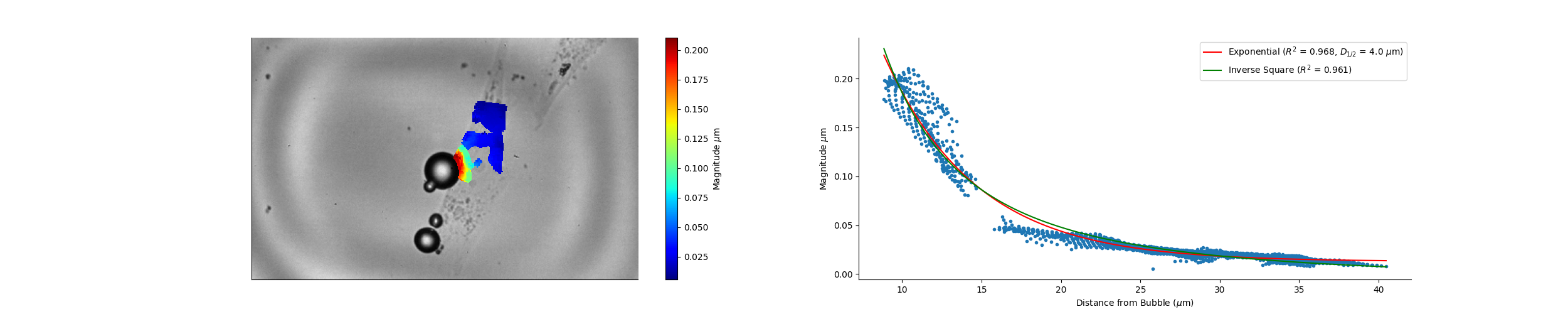

### Camera_16_43_05_Wave_Speed.png

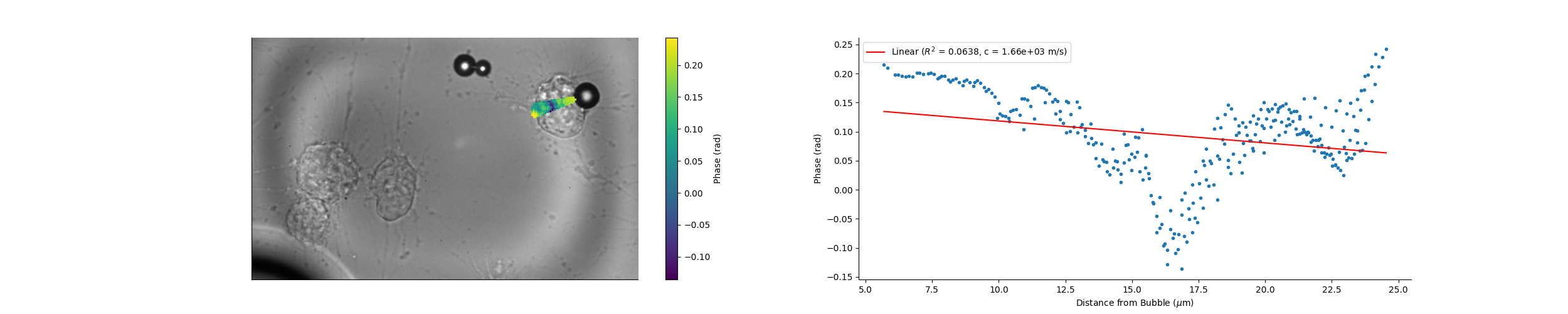

### Camera_16_43_55_Magnitude.png

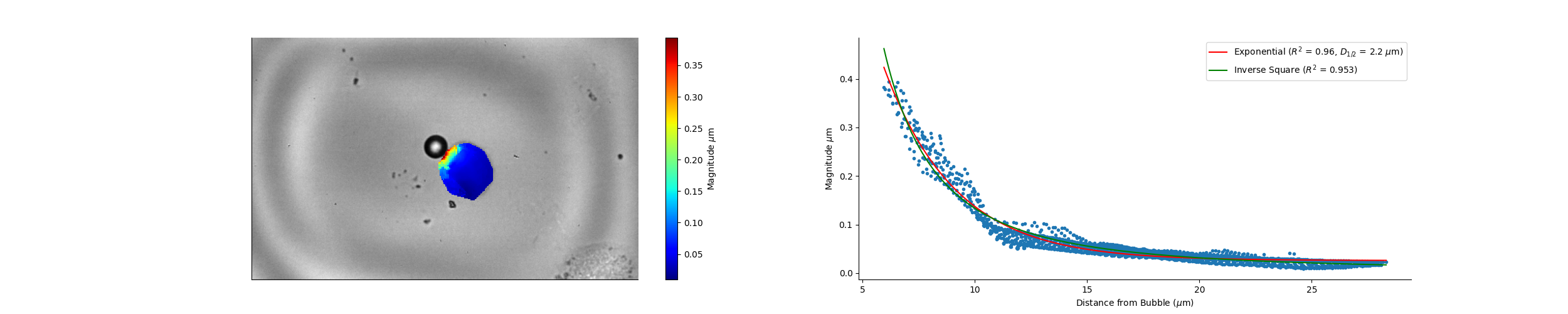

### Camera_16_43_55_Wave_Speed.png

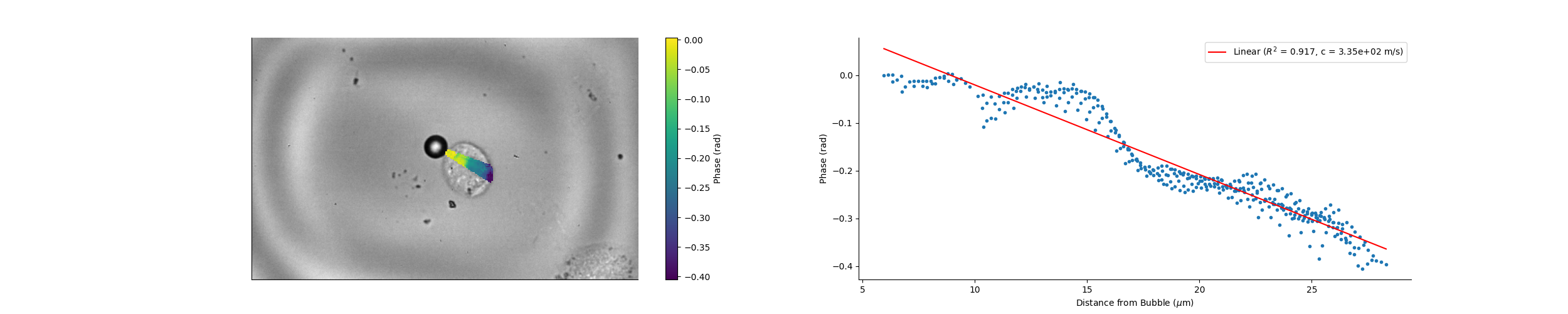

### Camera_16_47_28_Magnitude.png

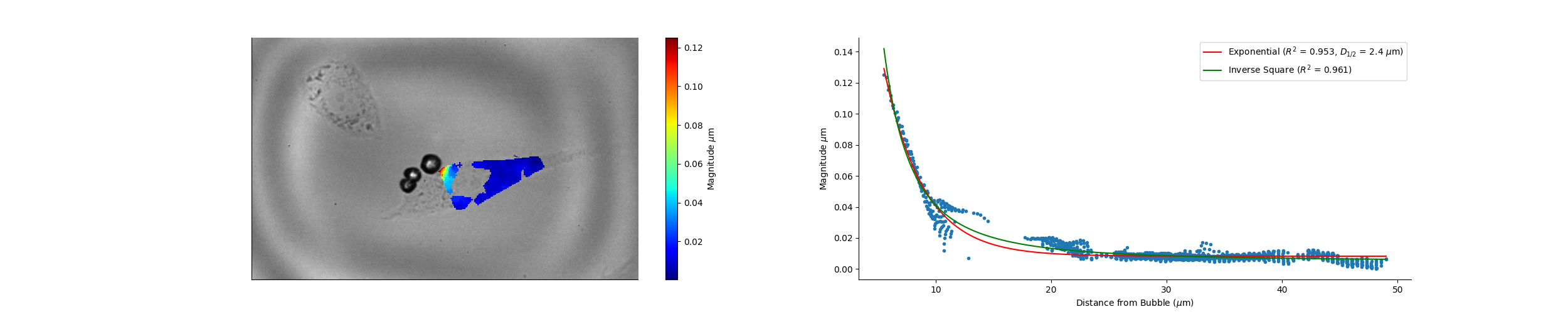

### Camera_16_51_46_Wave_Speed.png

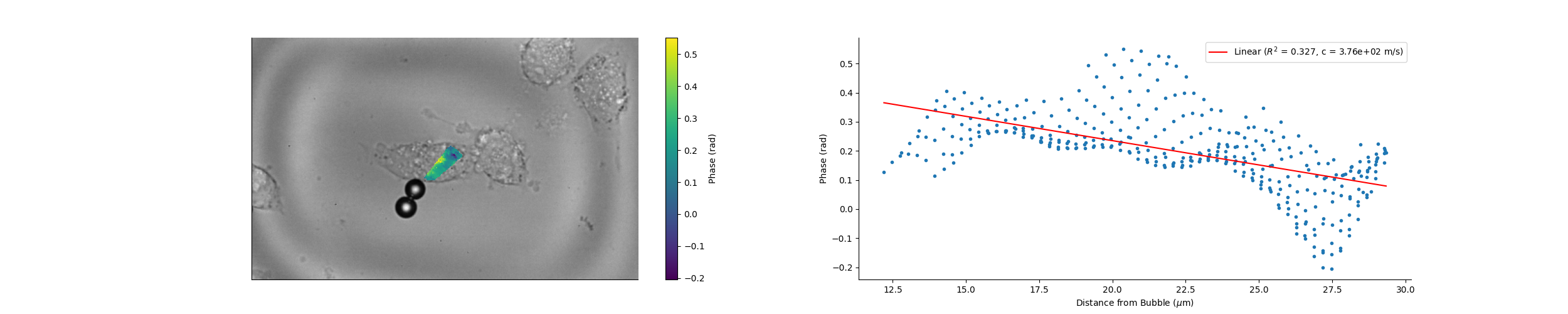

### Camera_16_53_25_Magnitude.png

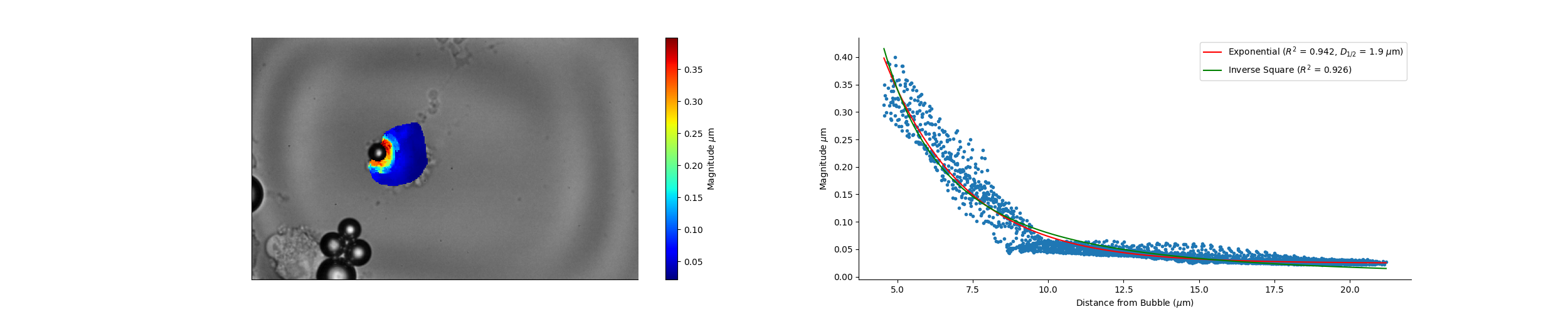

### Camera_16_56_27_Wave_Speed.png

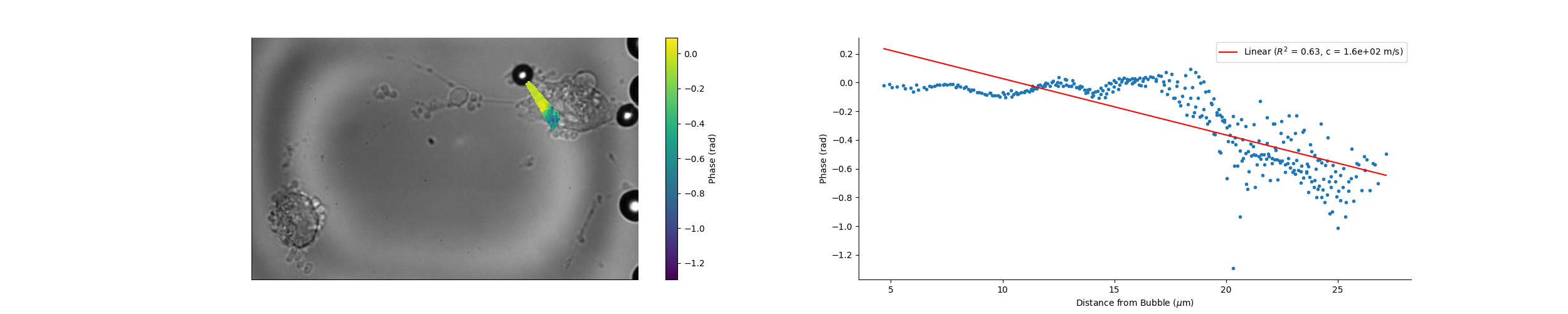

### Camera_16_57_16_Wave_Speed.png

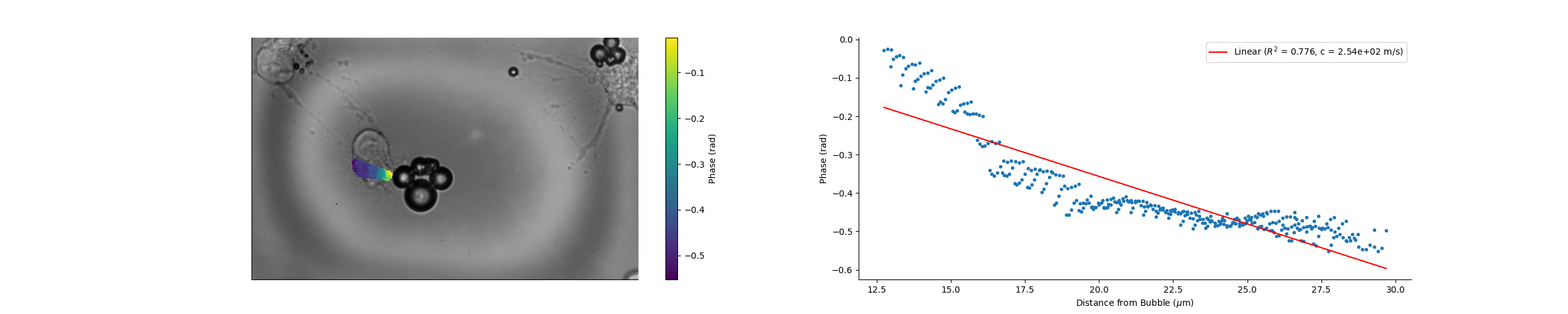

### Camera_16_57_40_Wave_Speed.png

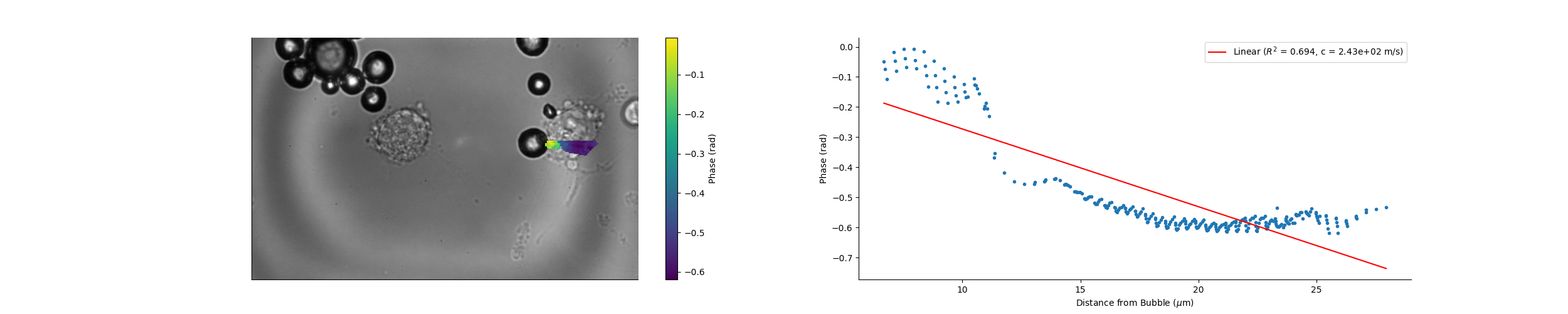

### Camera_17_02_49_Wave_Speed.png

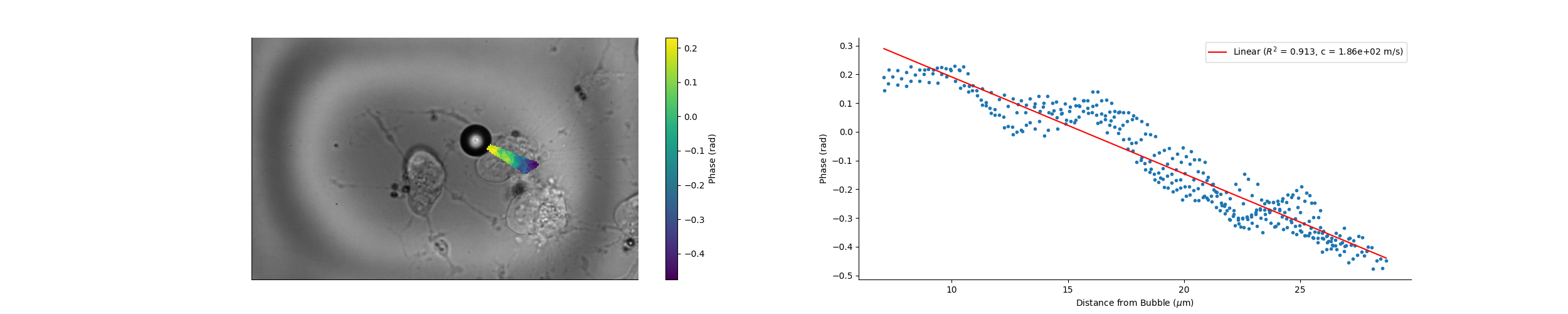

### Camera_17_07_37_Wave_Speed.png

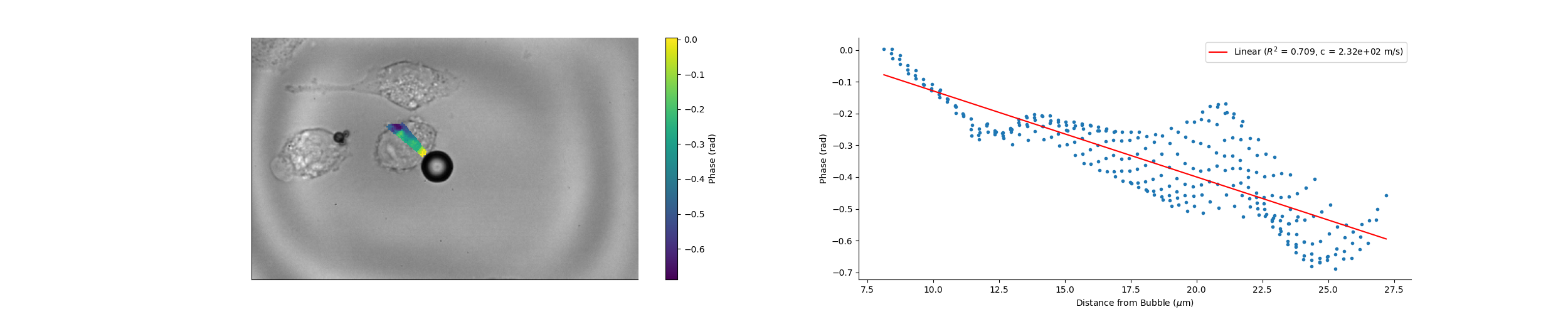

### Camera_17_11_05_Magnitude.png

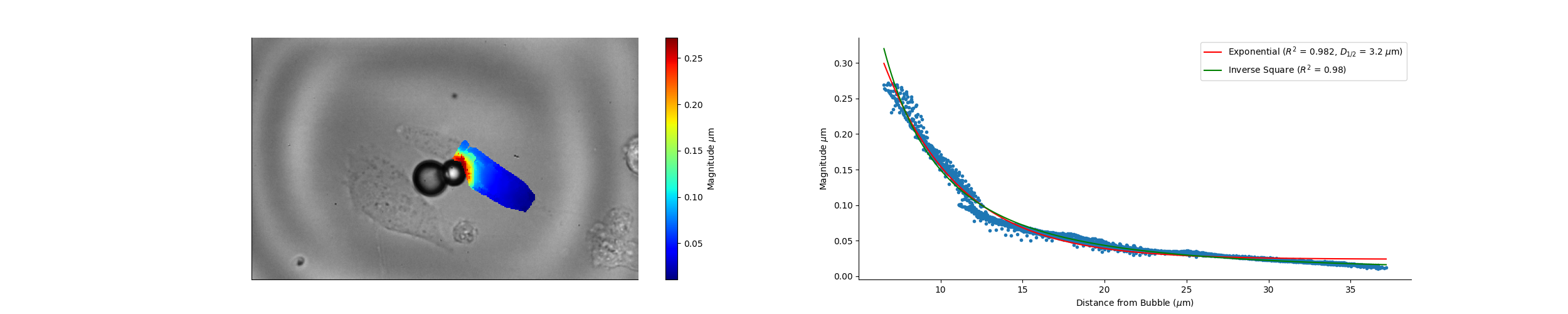

### Camera_17_11_05_Wave_Speed.png

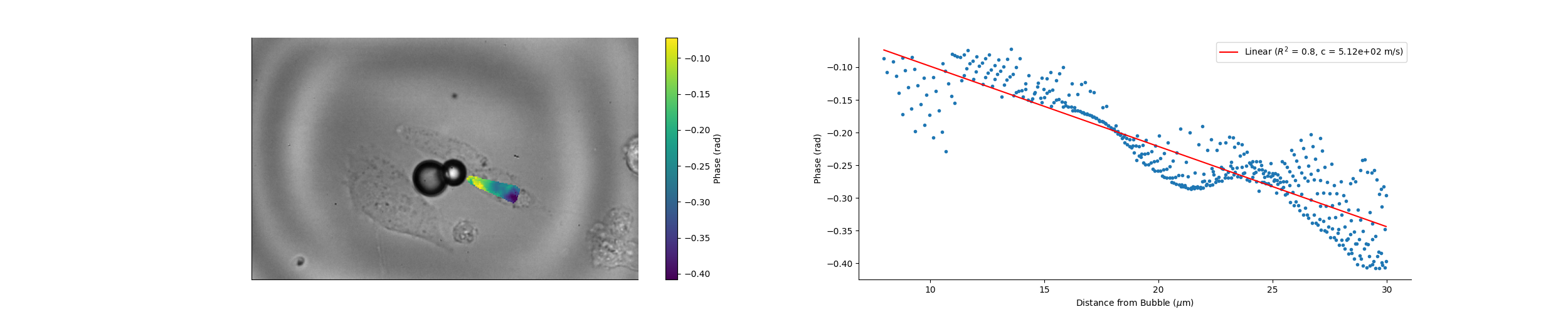

### Camera_17_11_58_Magnitude.png

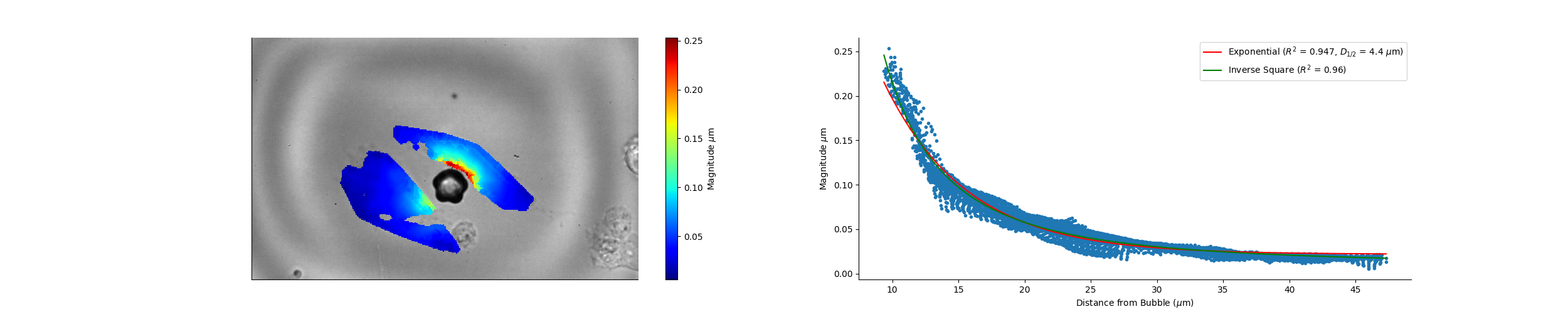

### Camera_17_11_58_Wave_Speed.png

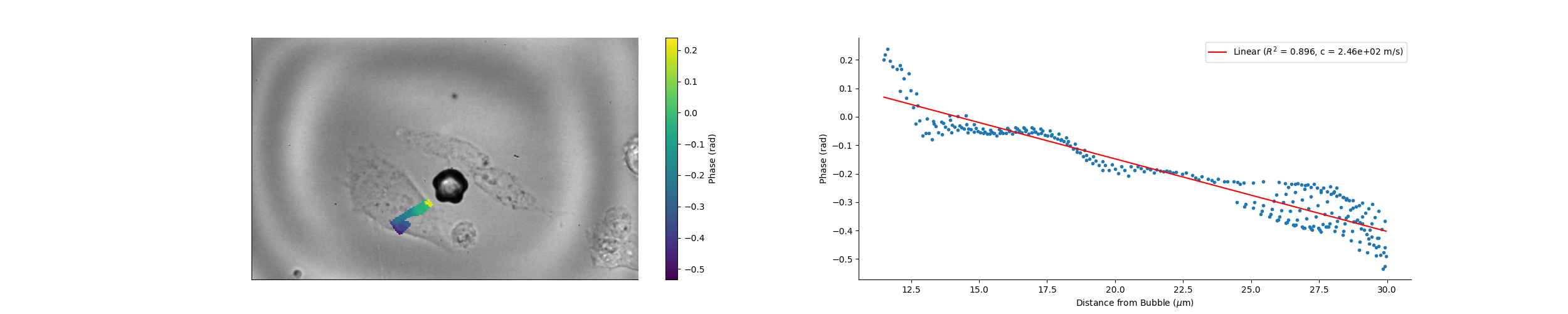

### Camera_17_14_56_Magnitude.png

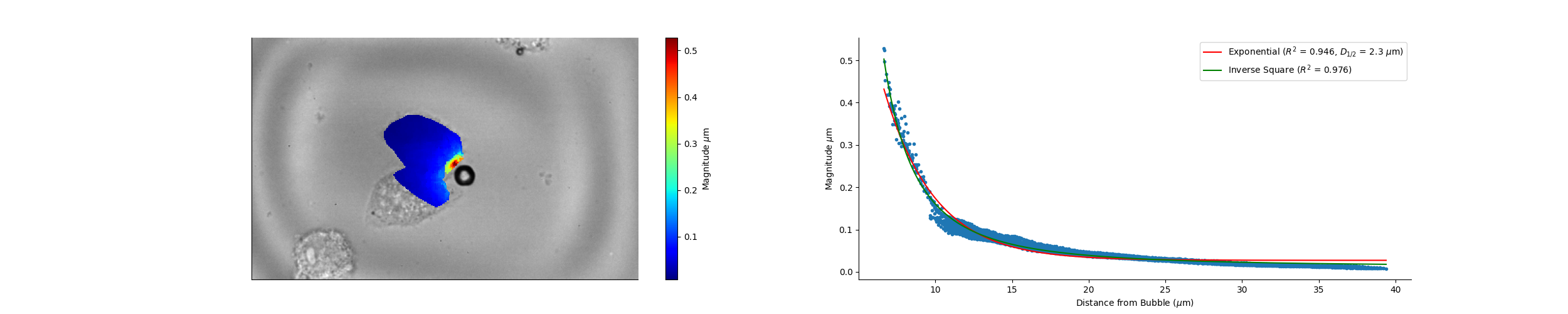

### Camera_17_14_56_Wave_Speed.png

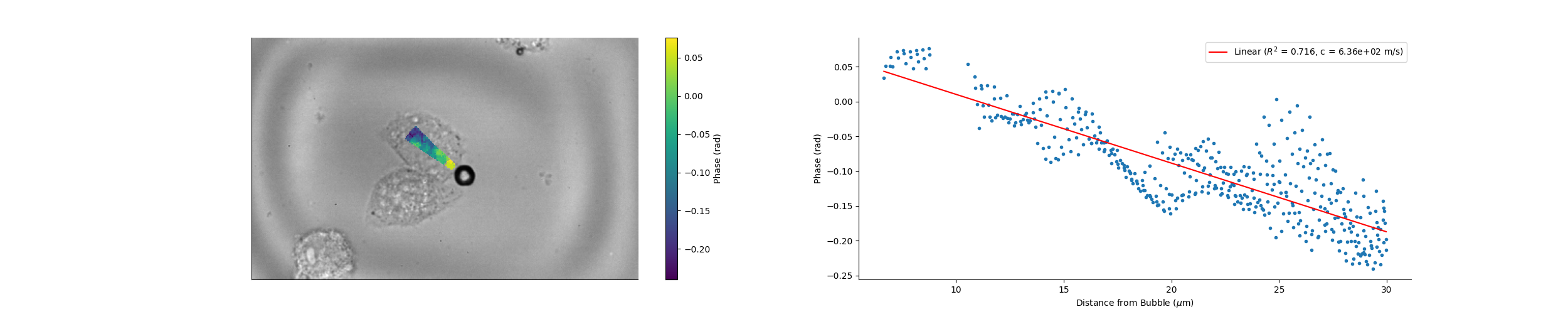

### Camera_17_15_54_Wave_Speed.png

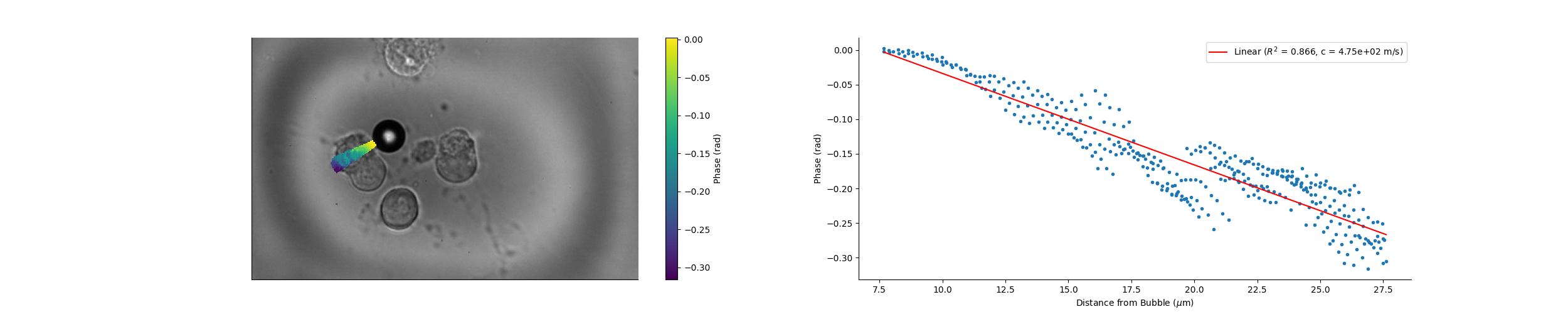

### Camera_17_16_20_Magnitude.png

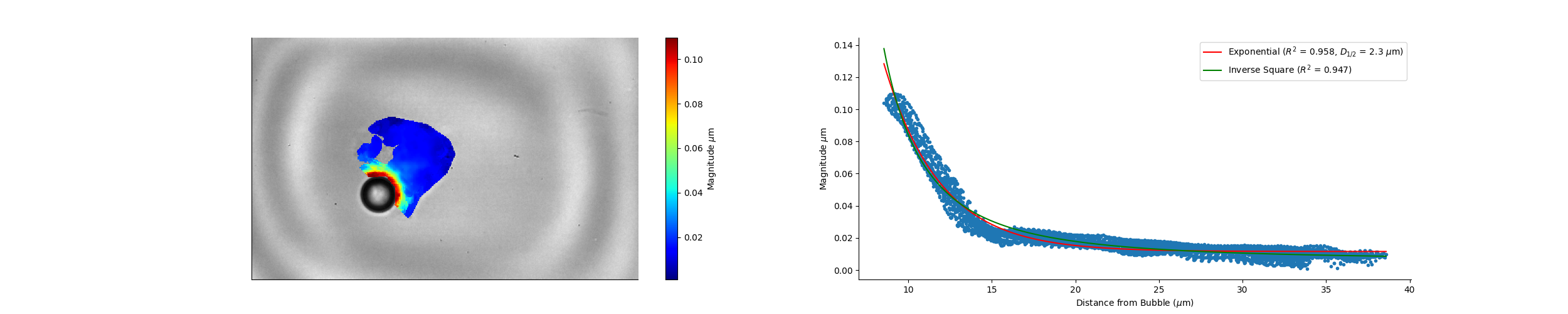

### Camera_17_17_18_Wave_Speed.png

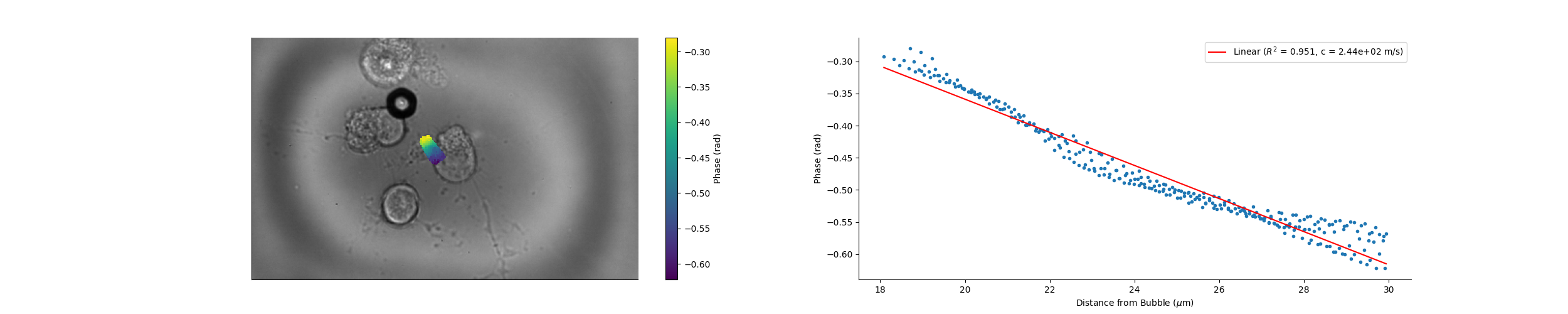

### Camera_17_17_31_Magnitude.png

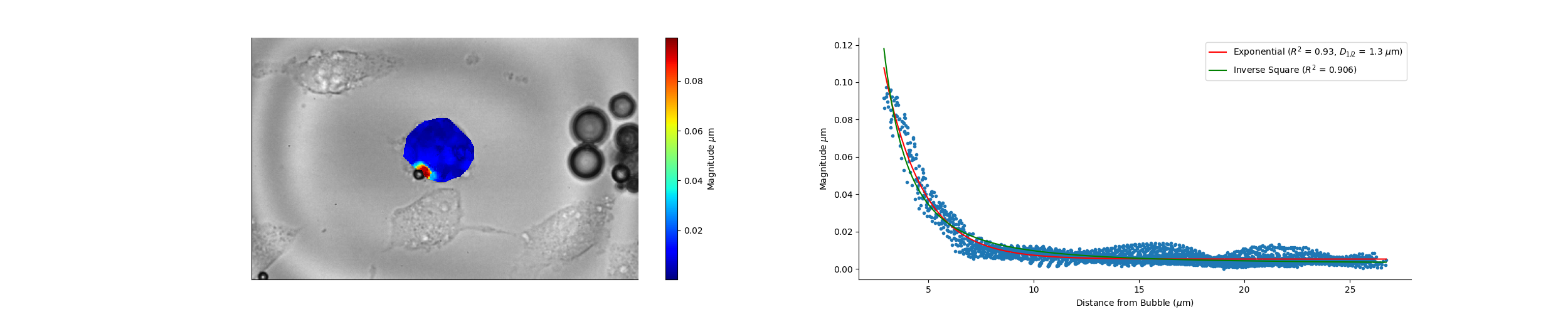

### Camera_17_20_02_Magnitude.png

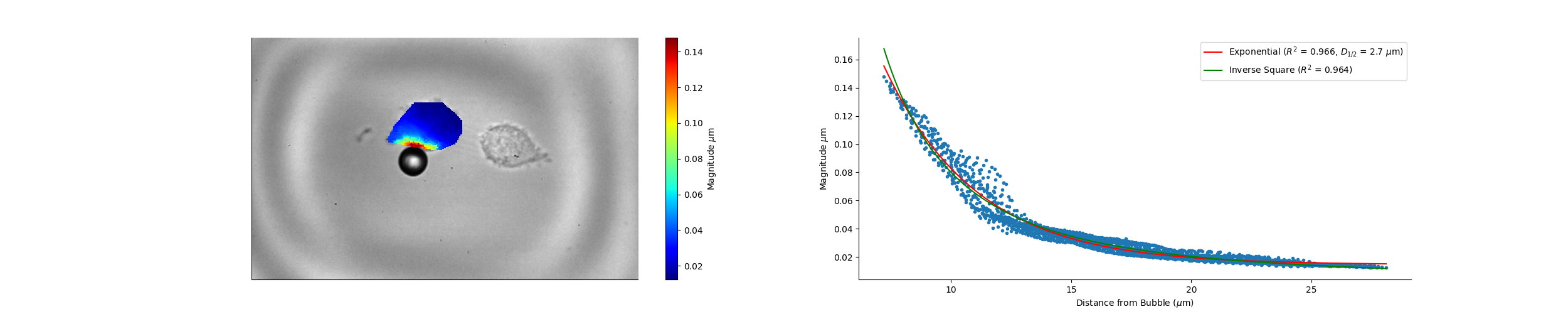

### Camera_17_20_02_Wave_Speed.png

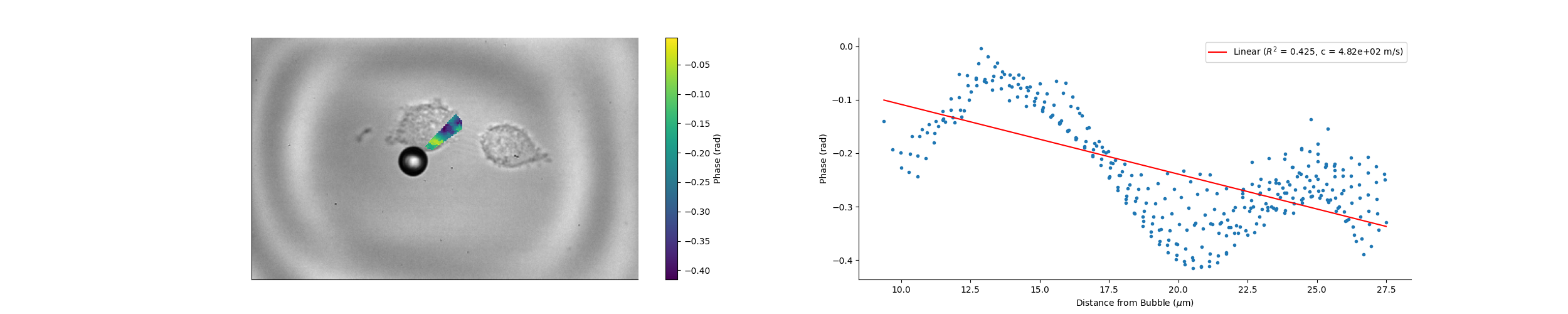

### Camera_17_21_43_Wave_Speed.png

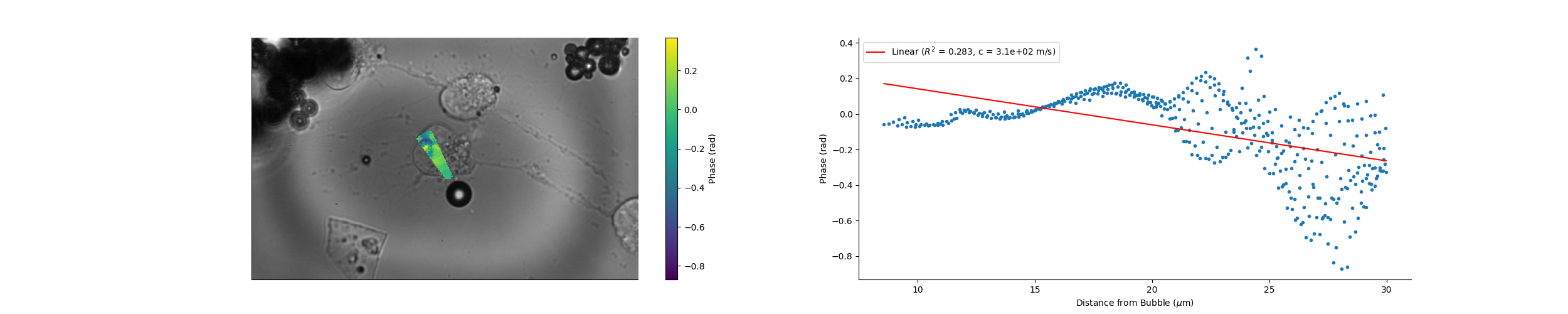

### Camera_17_22_49_Magnitude.png

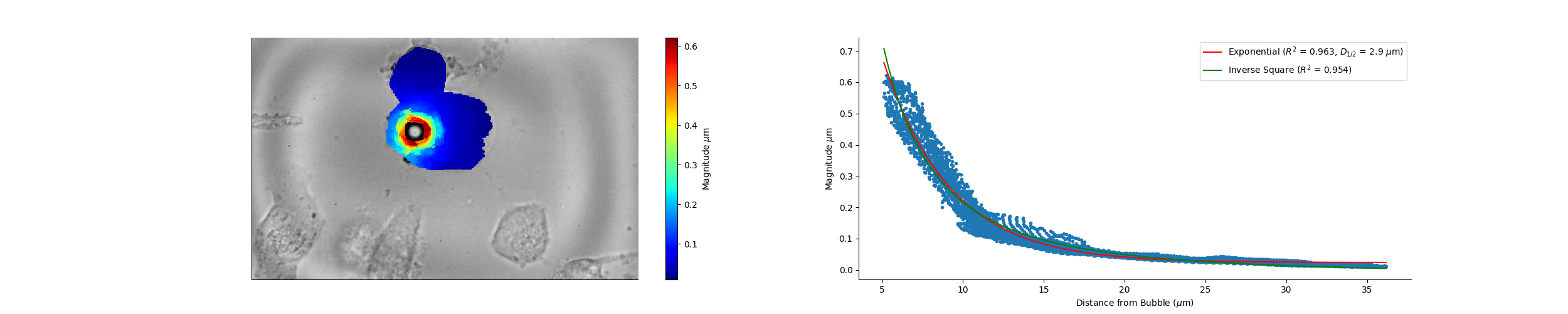

### Camera_17_22_49_Wave_Speed.png

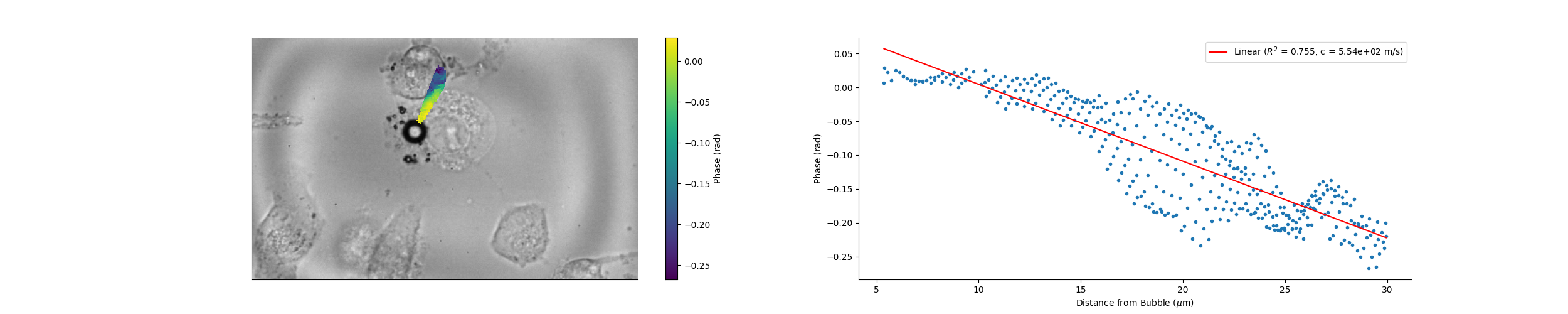

### Camera_17_29_58_Magnitude.png

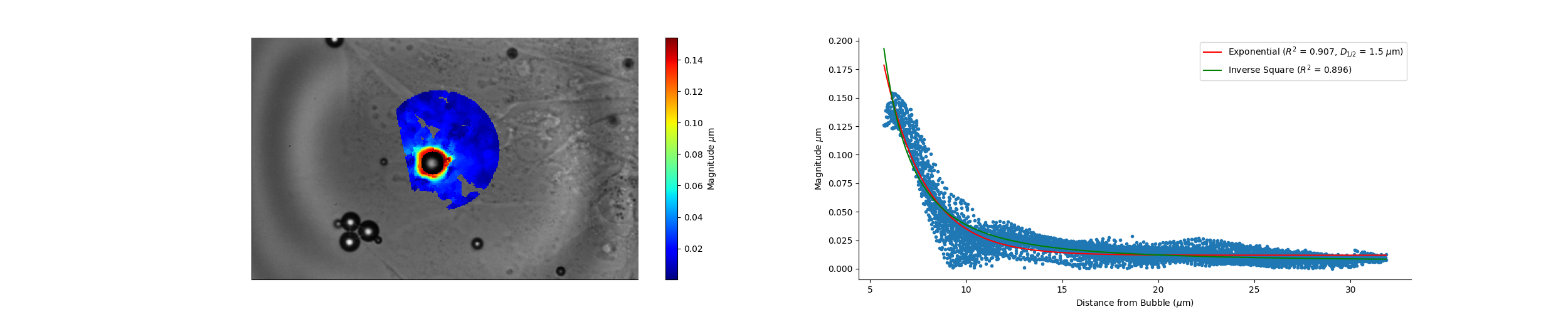

### Camera_17_29_58_Wave_Speed.png

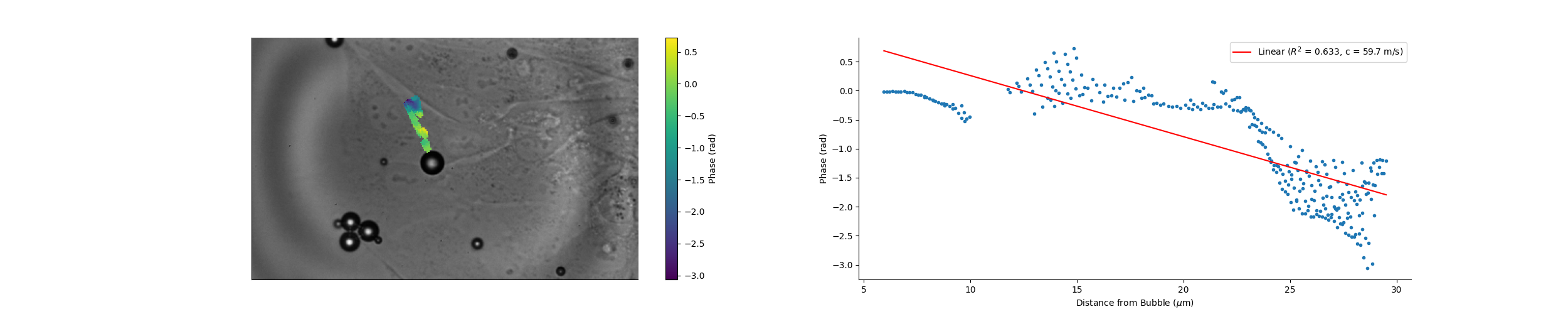

### Camera_17_30_42_Magnitude.png

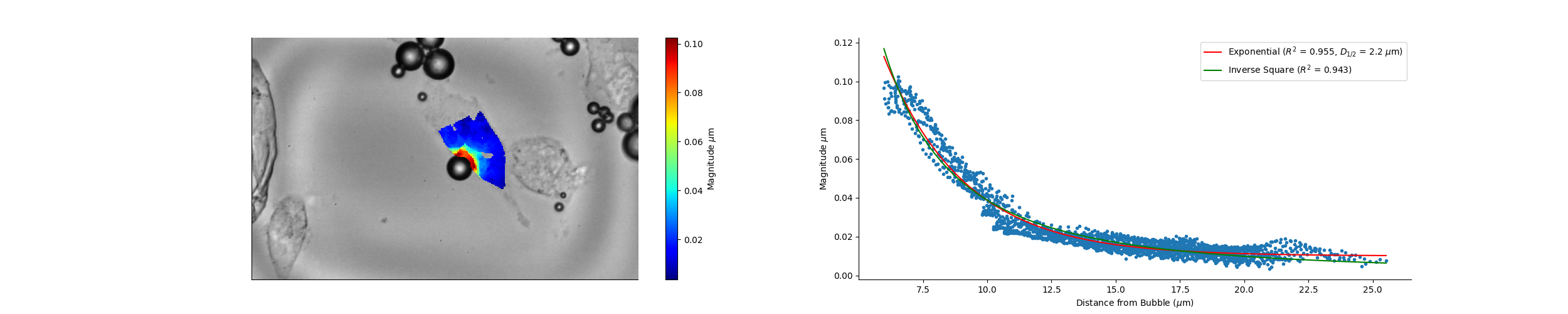

### Camera_17_30_42_Wave_Speed.png

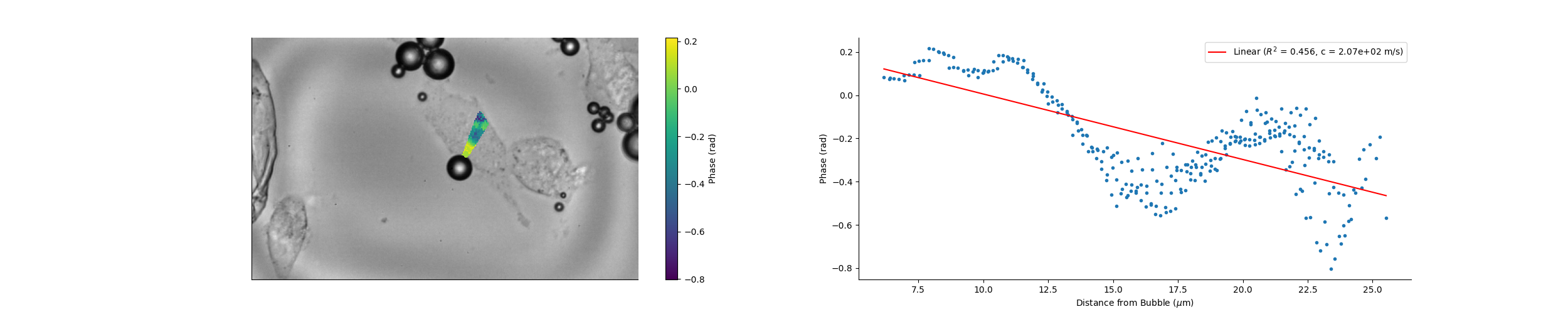

### Camera_17_40_29_Magnitude.png

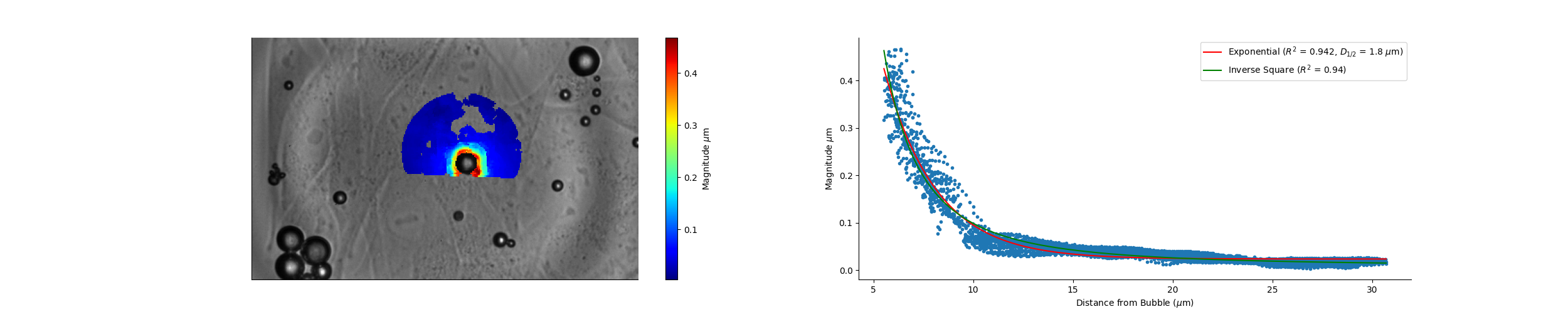

### Camera_17_40_29_Wave_Speed.png

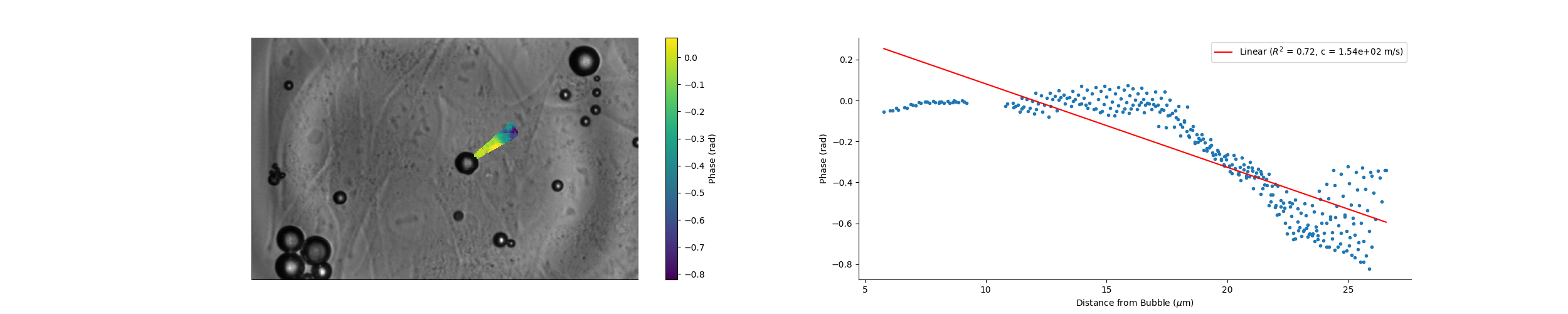

### Camera_17_41_58_Magnitude.png

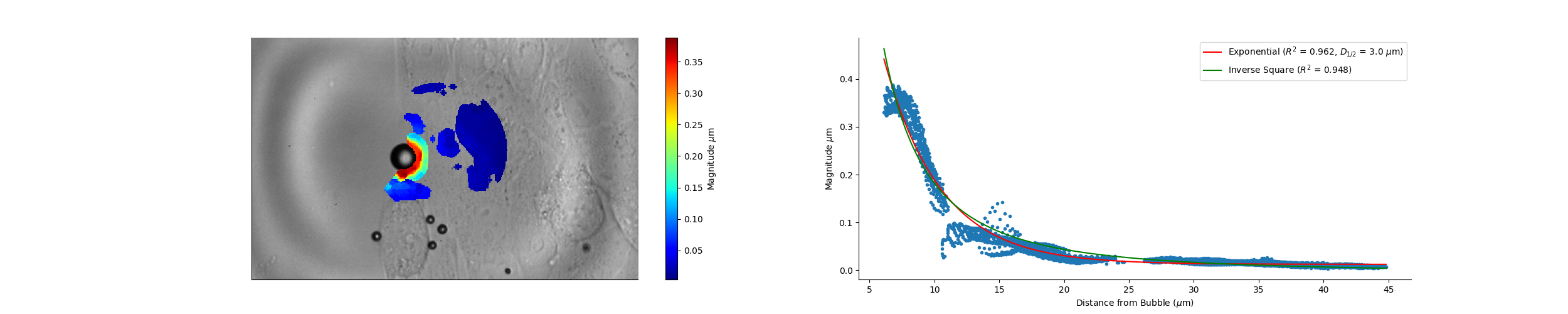

### Camera_17_42_59_Magnitude.png

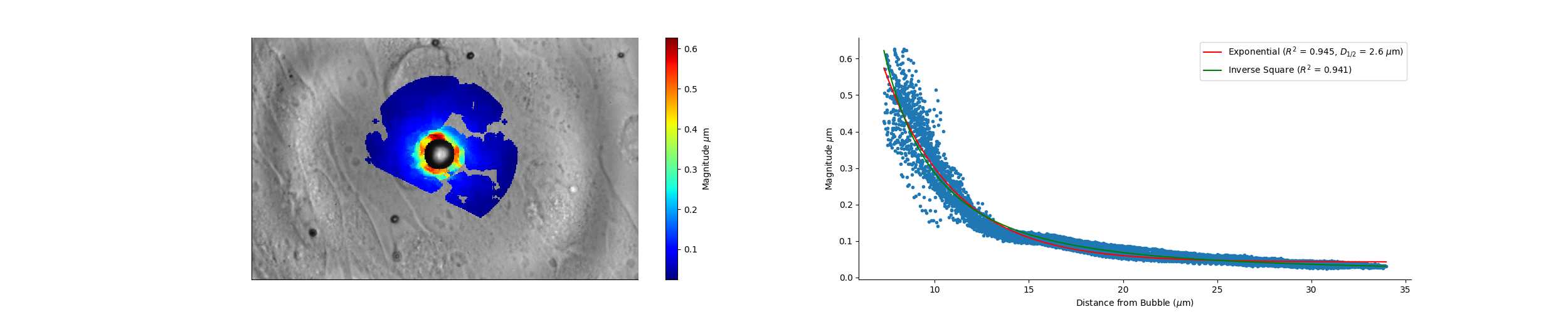

### Camera_17_42_59_Wave_Speed.png

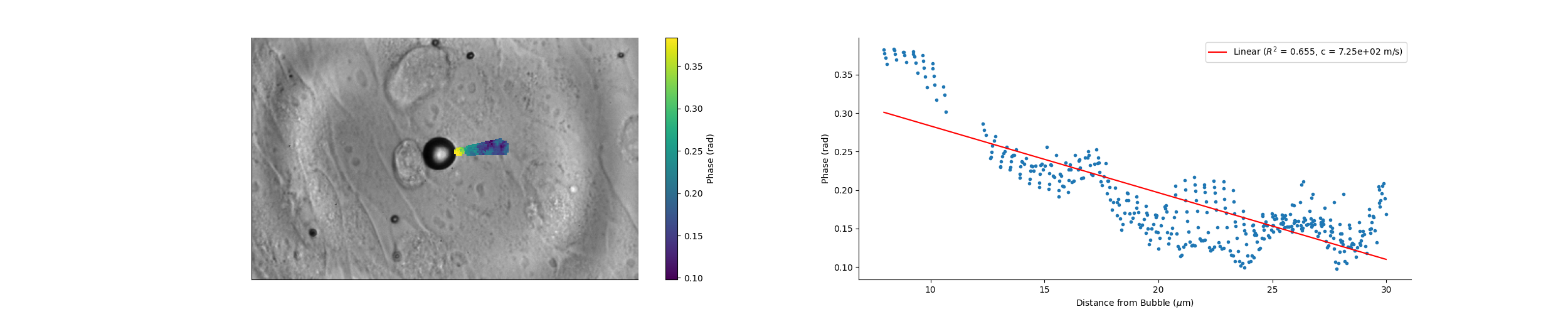

### Camera_17_43_57_Magnitude.png

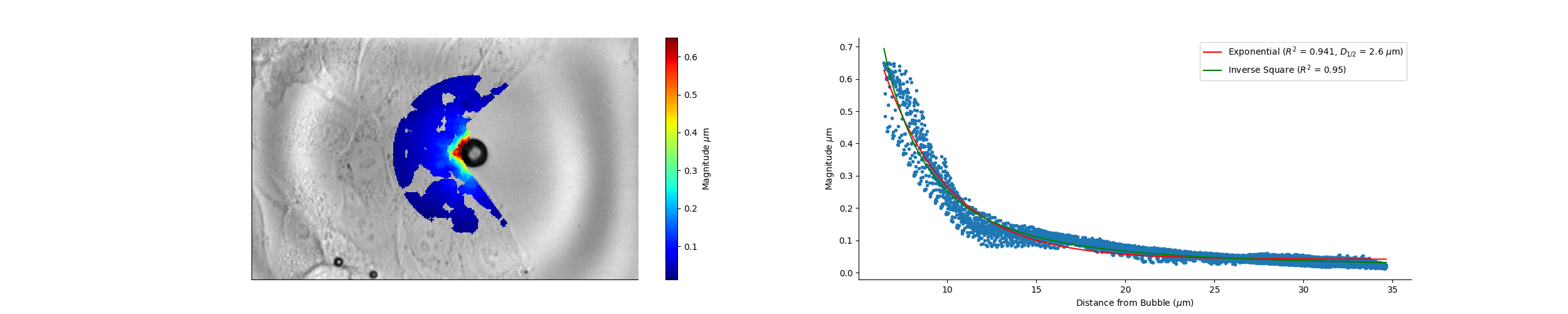

### Camera_17_43_57_Wave_Speed.png

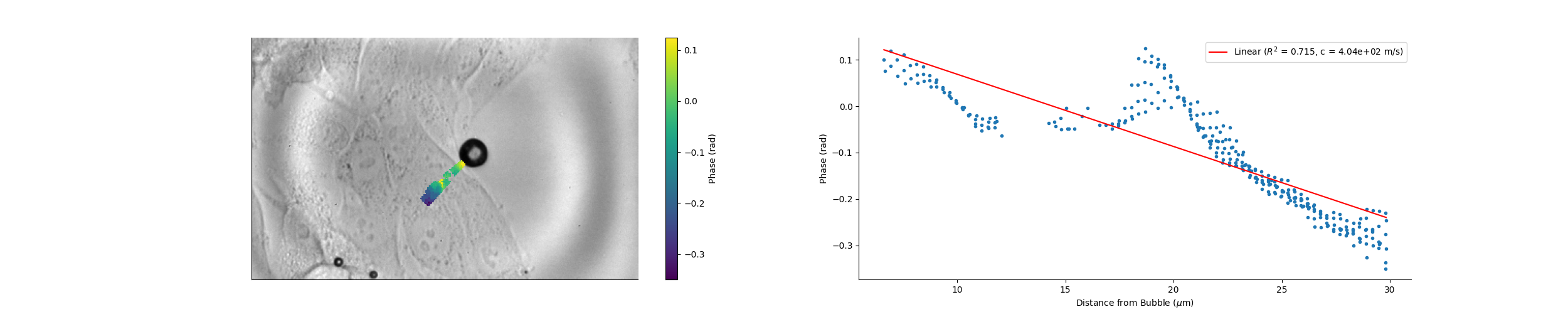

### Camera_17_44_57_Magnitude.png

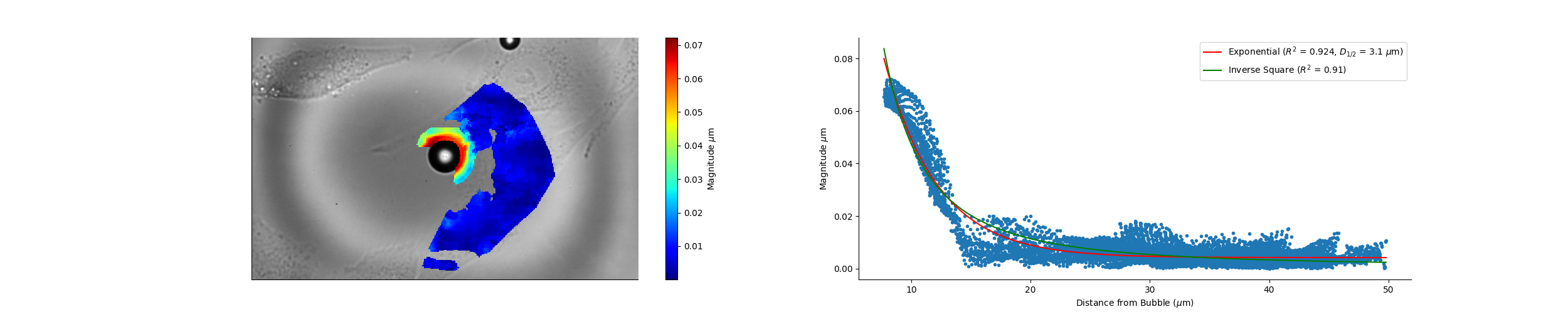

### Camera_17_44_57_Wave_Speed.png

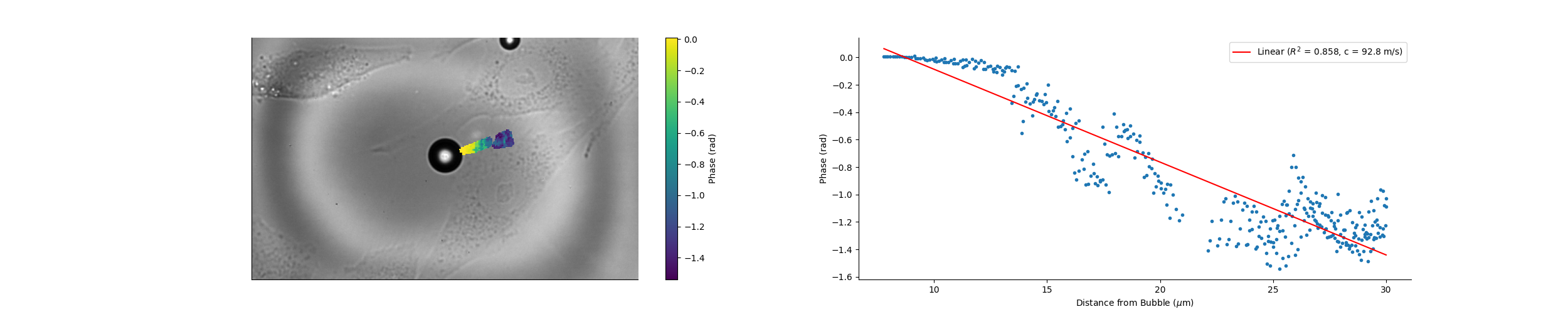

### Camera_17_48_17_Magnitude.png

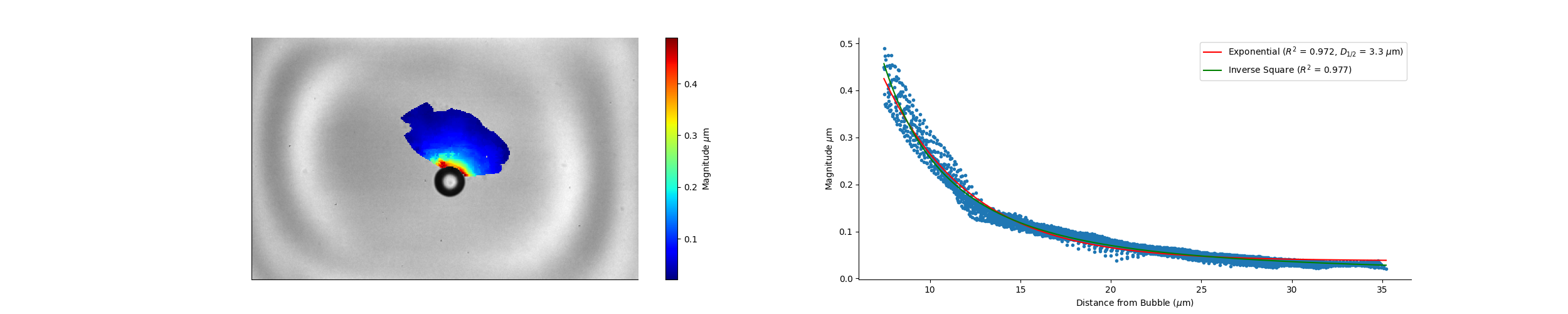

### Camera_17_48_17_Wave_Speed.png

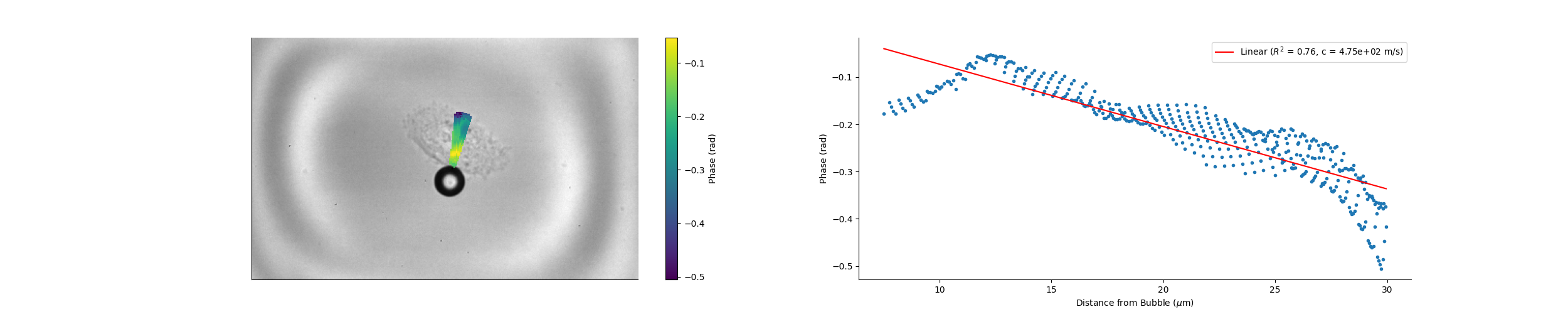

### Camera_17_49_47_Magnitude.png

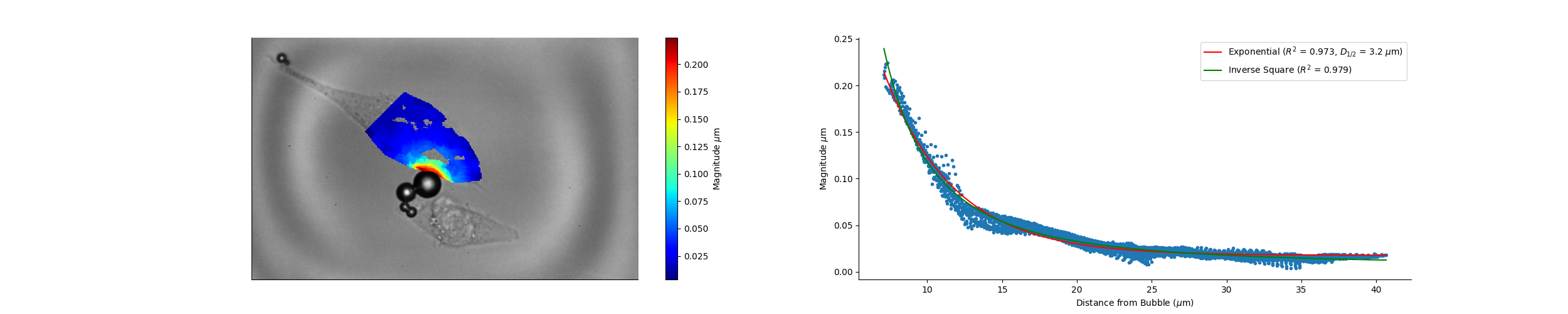

### Camera_17_49_53_Magnitude.png

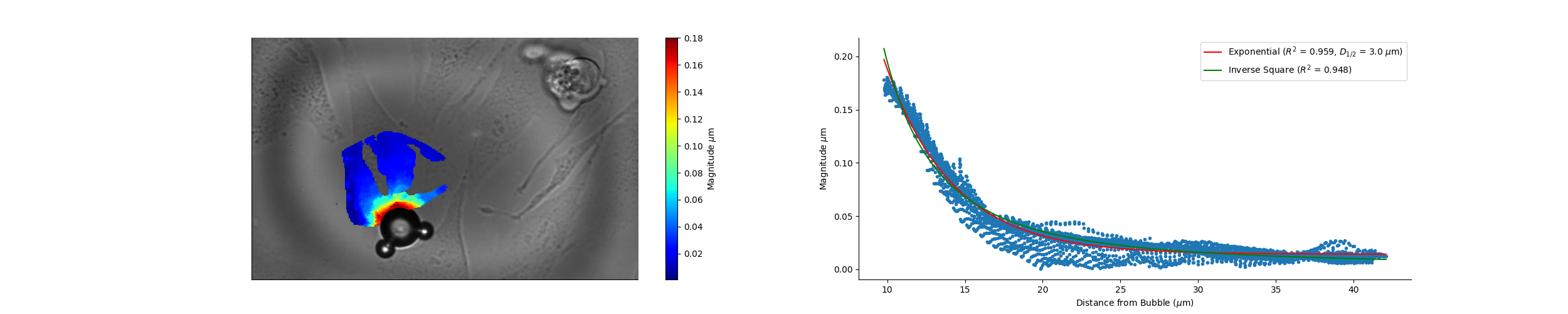

### Camera_17_50_46_Magnitude.png

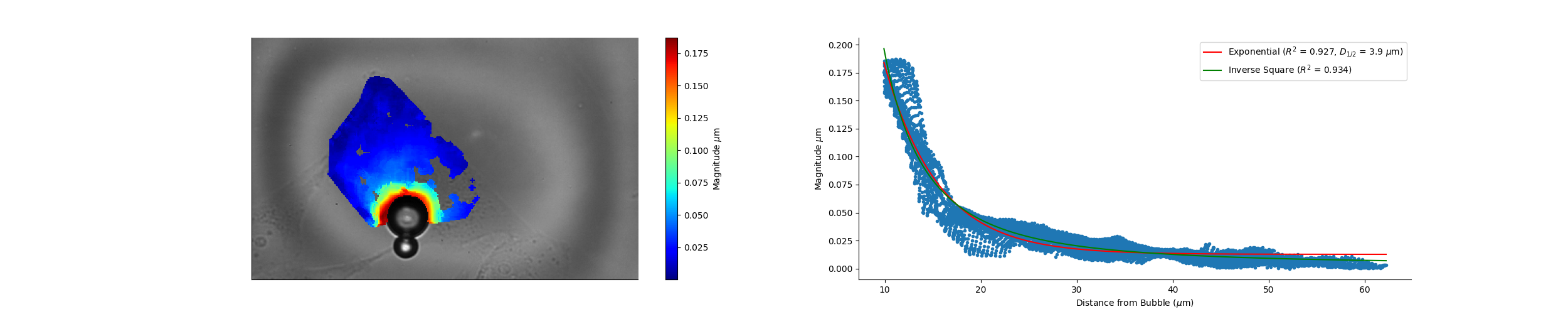

### Camera_17_51_54_Magnitude.png

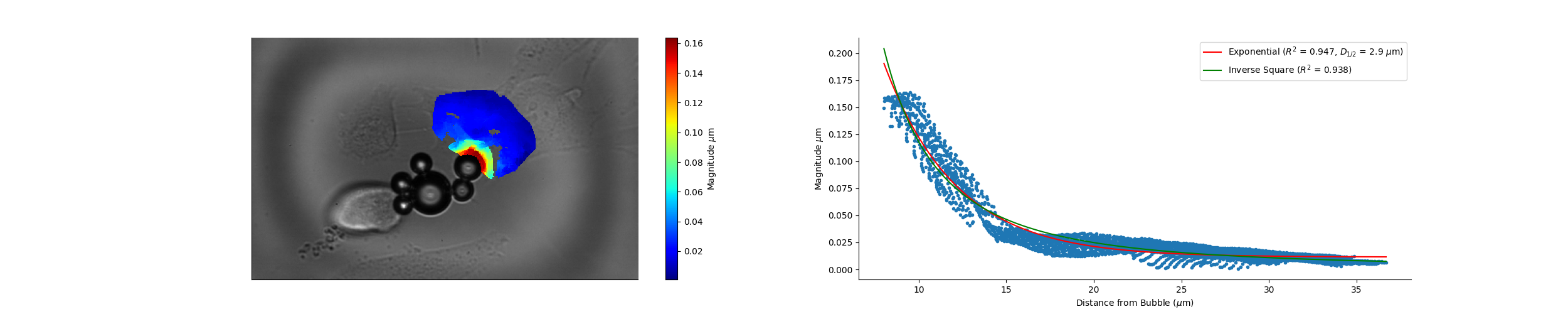

### Camera_17_53_30_Magnitude.png

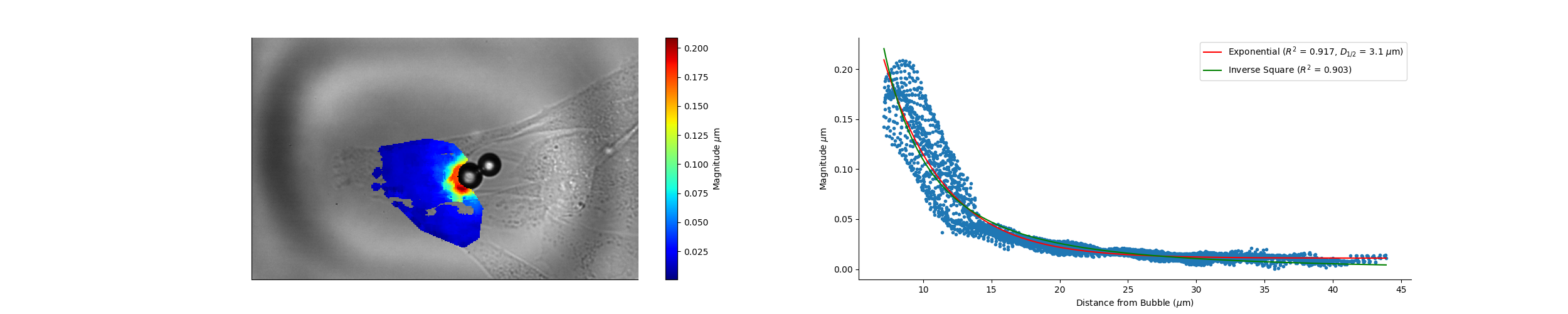

### Camera_17_54_25_Magnitude.png

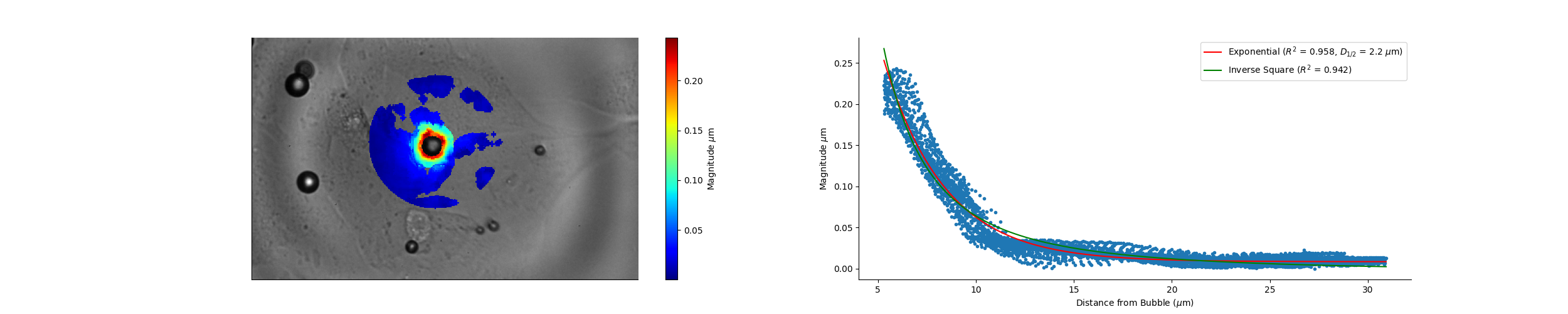

### Camera_17_55_30_Magnitude.png

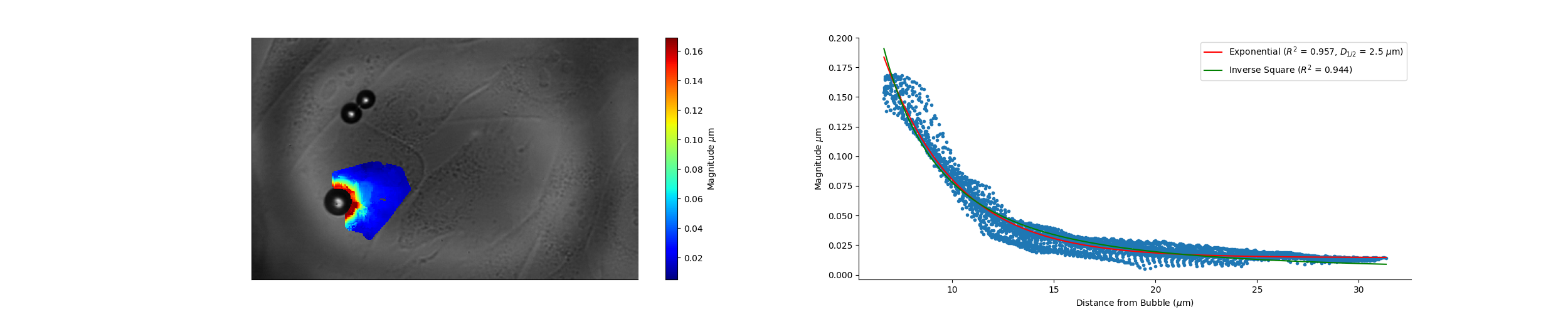

### Camera_17_55_33_Magnitude.png

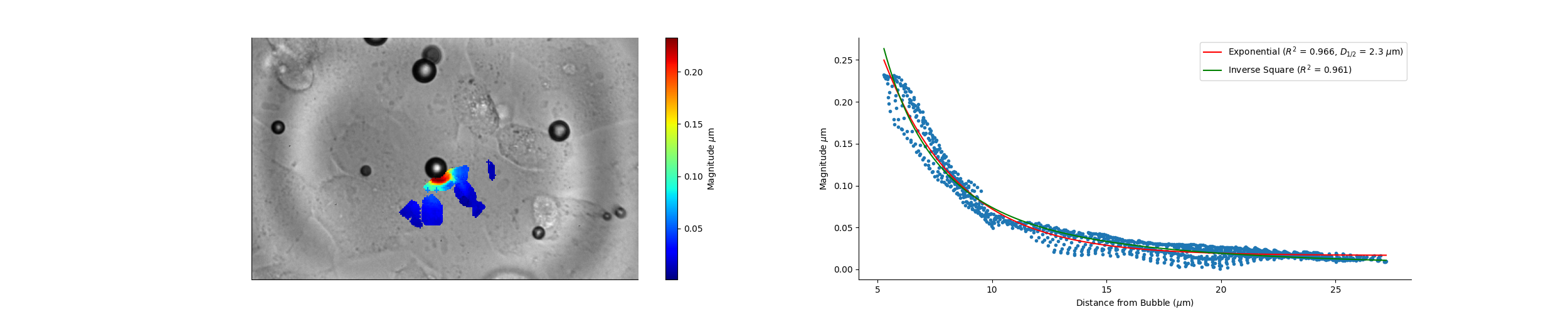

### Camera_17_57_34_Magnitude.png

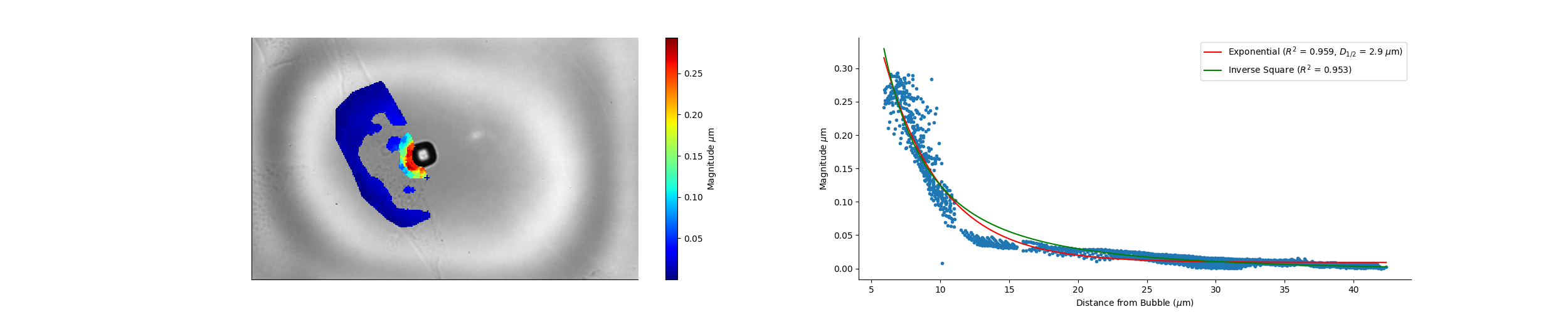

### Camera_17_58_32_Magnitude.png

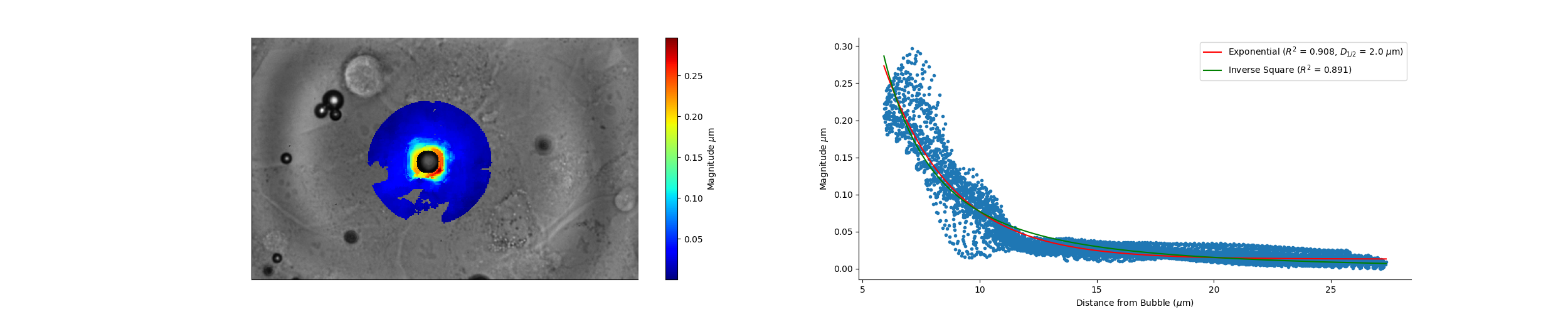

### Camera_17_59_39_Magnitude.png

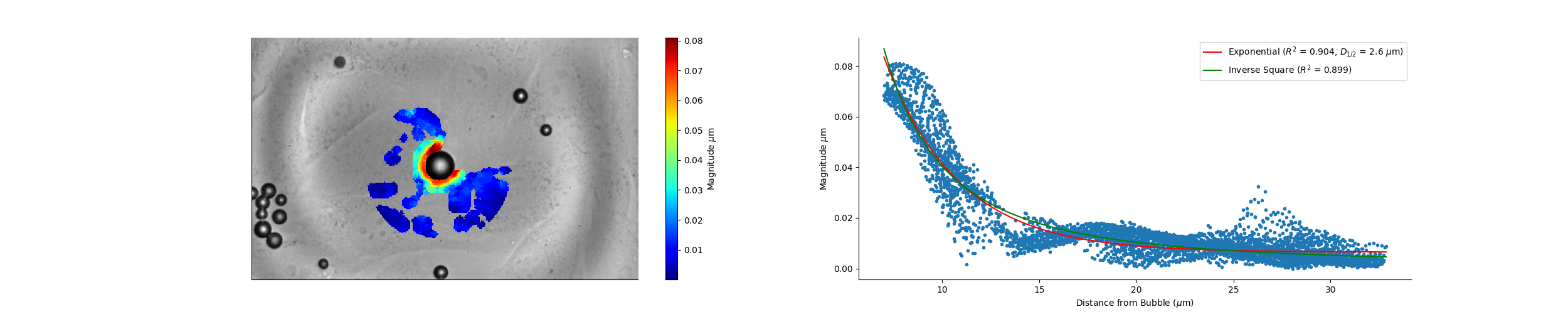

### Camera_18_03_52_Magnitude.png

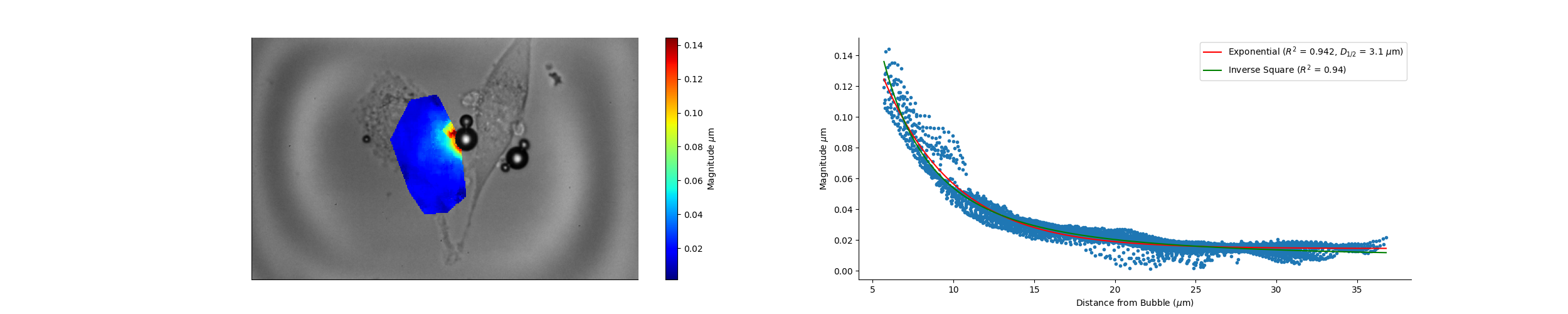

### Camera_18_06_31_Magnitude.png

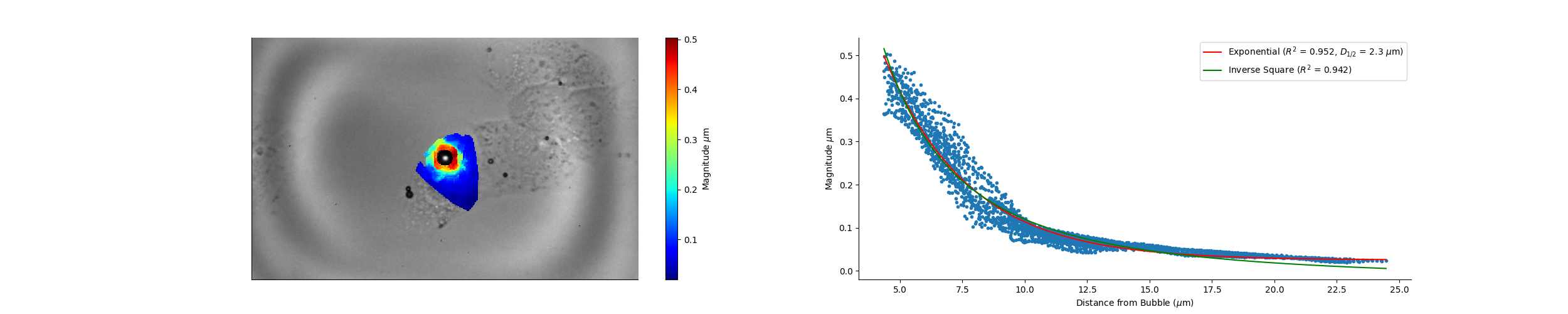

### Camera_18_17_59_Magnitude.png

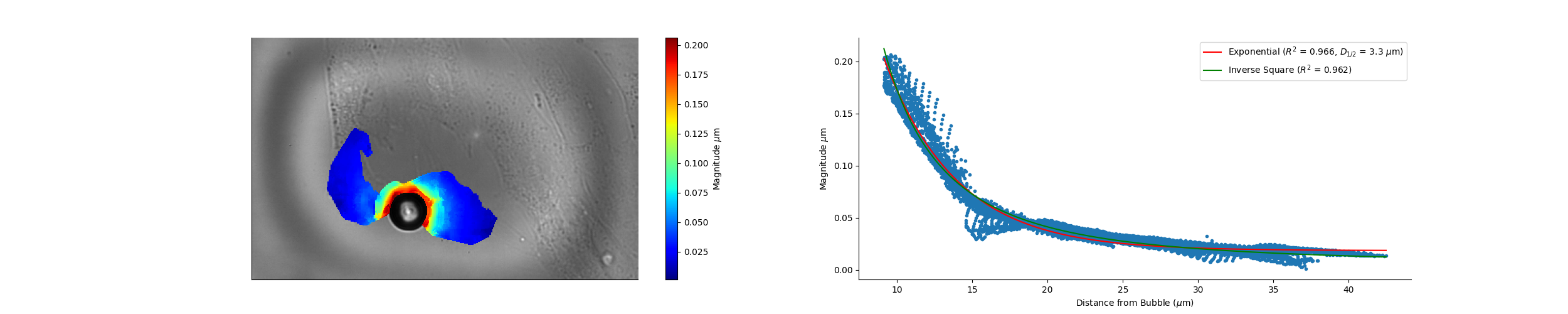

### Camera_18_22_27_Magnitude.png

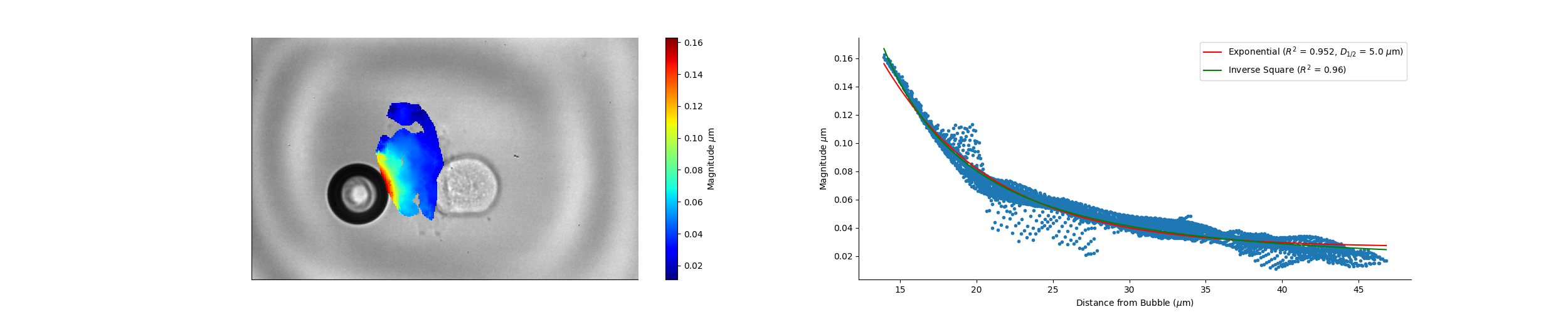

### Camera_18_22_27_Wave_Speed.png

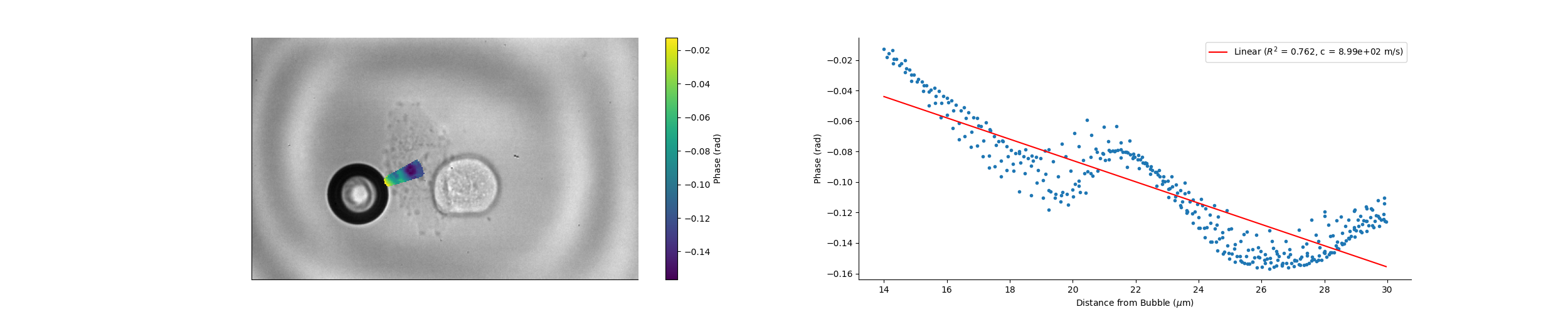
