## Supplementary figures and images for "Cell material state determines high-frequency cell deformation and microbubble-induced permeabilization"

### Camera_14_23_25_Wave_Speed.png

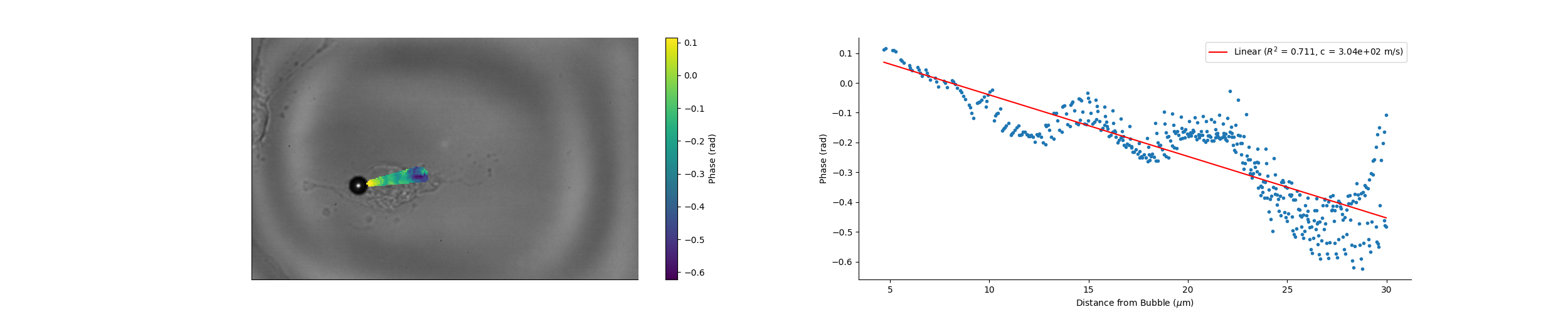

### Camera_14_25_31_Magnitude.png

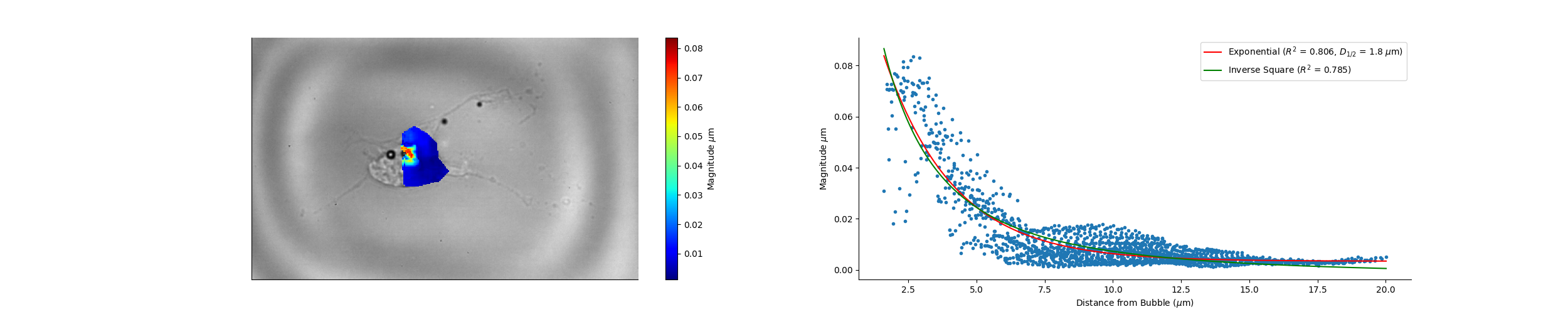

### Camera_14_36_14_Magnitude.png

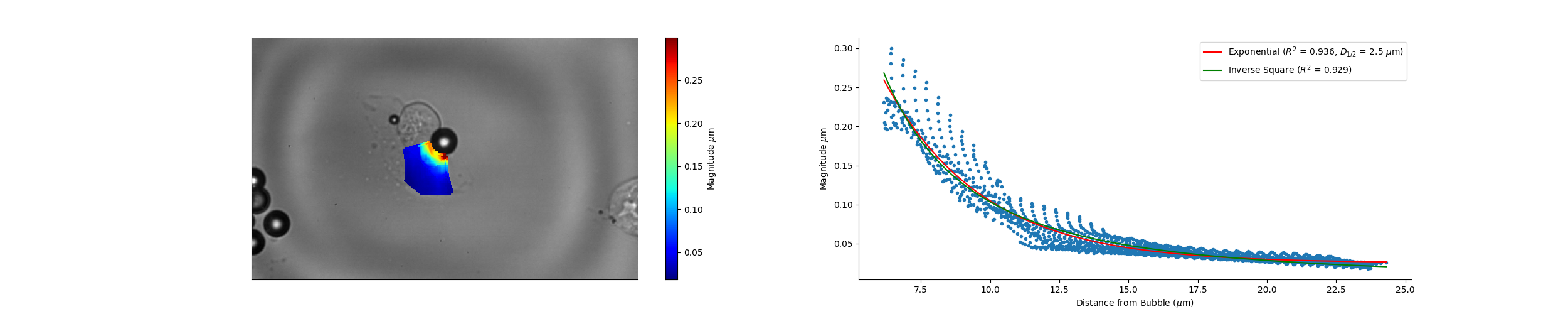

### Camera_14_43_17_Magnitude.png

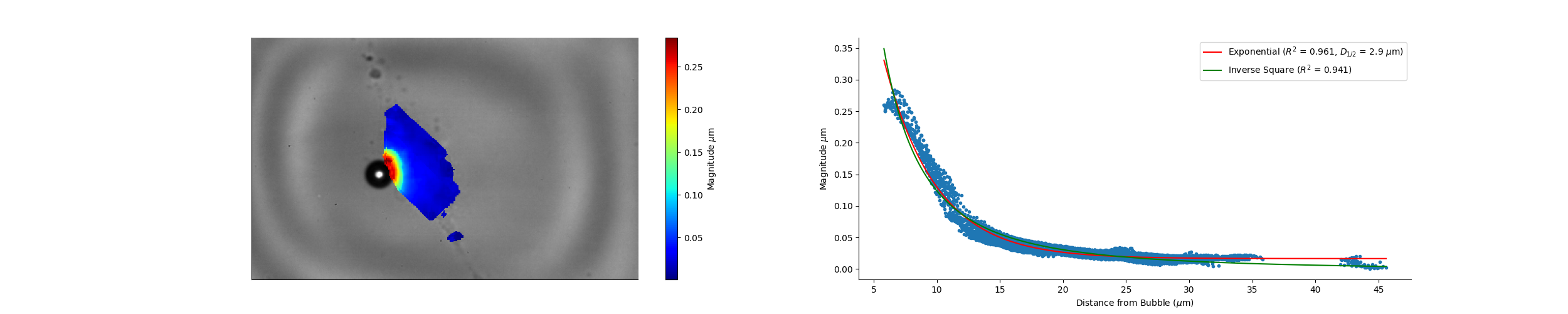

### Camera_14_43_17_Wave_Speed.png

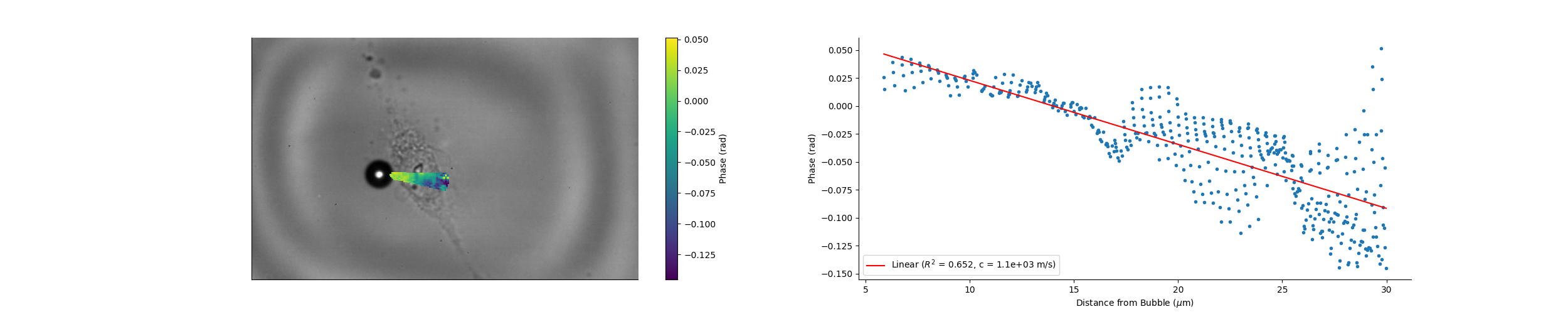

### Camera_14_44_12_Magnitude.png

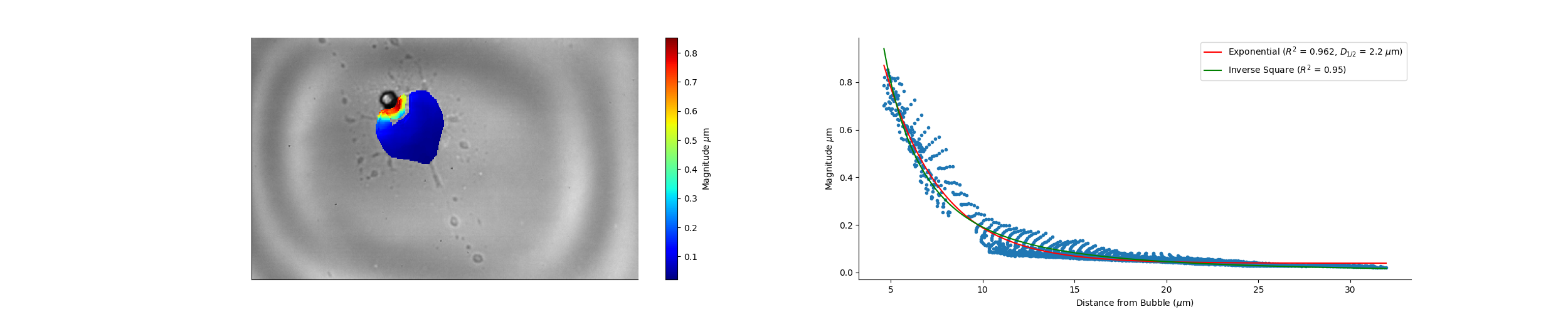

### Camera_14_44_12_Wave_Speed.png

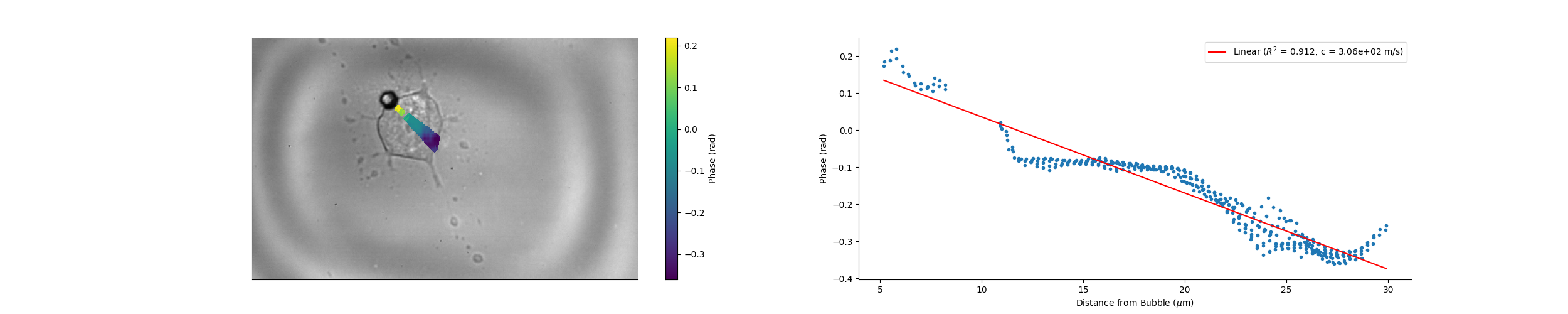

### Camera_14_45_29_Magnitude.png

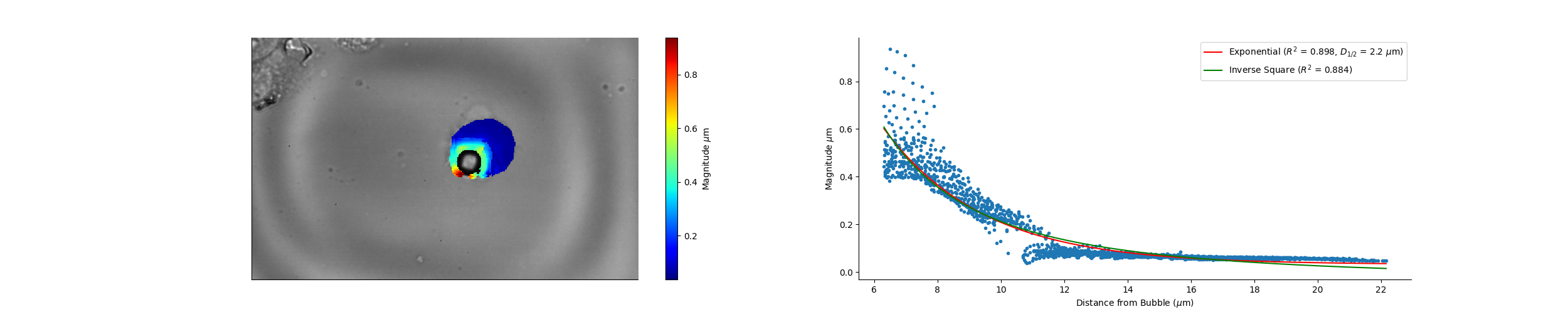

### Camera_14_47_04_Magnitude.png

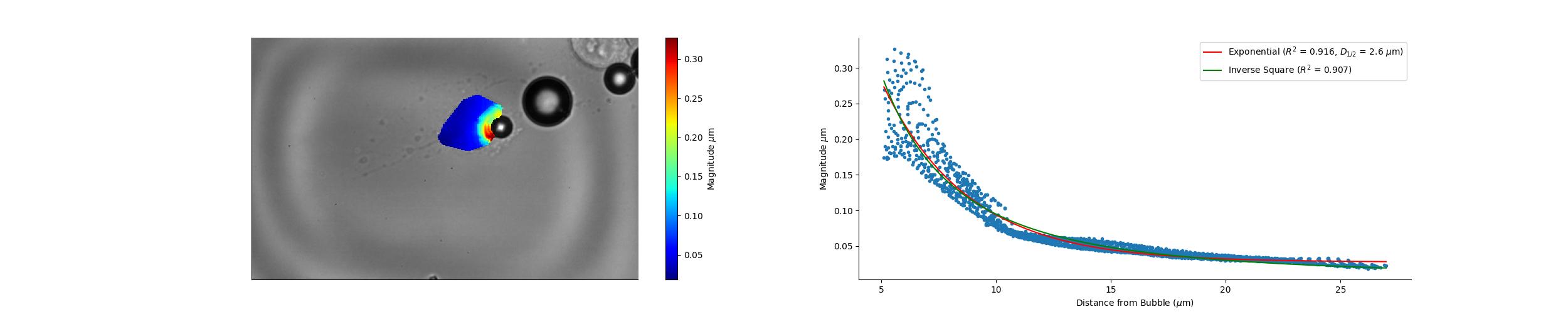

### Camera_14_47_04_Wave_Speed.png

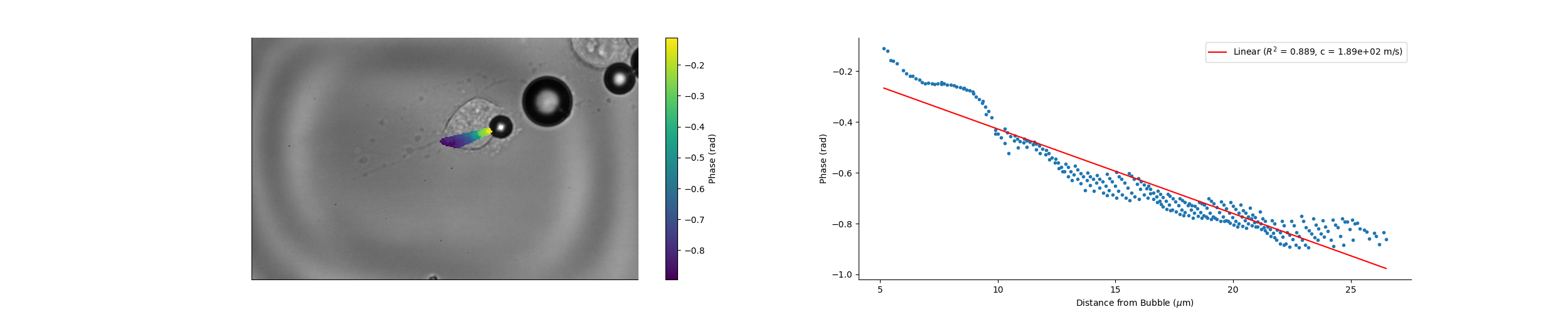

### Camera_14_48_16_Magnitude.png

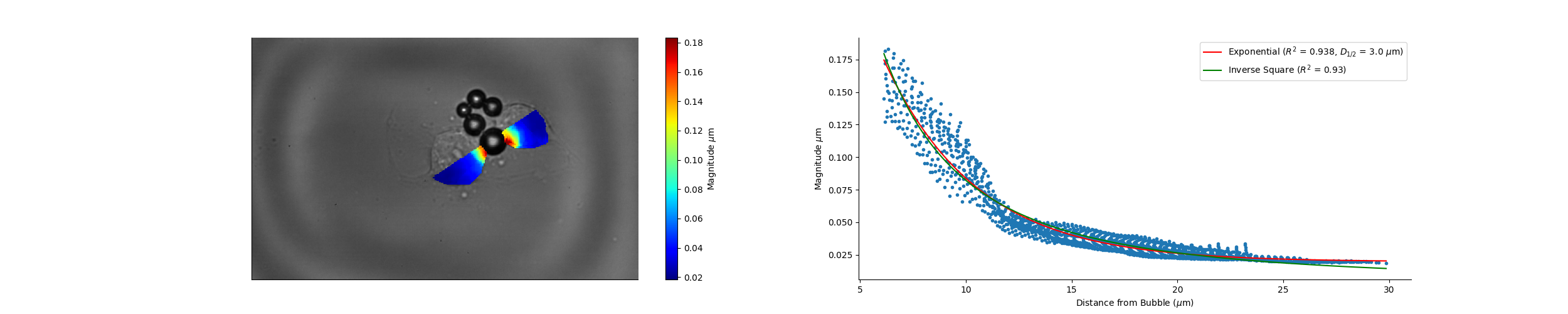

### Camera_14_48_16_Wave_Speed.png

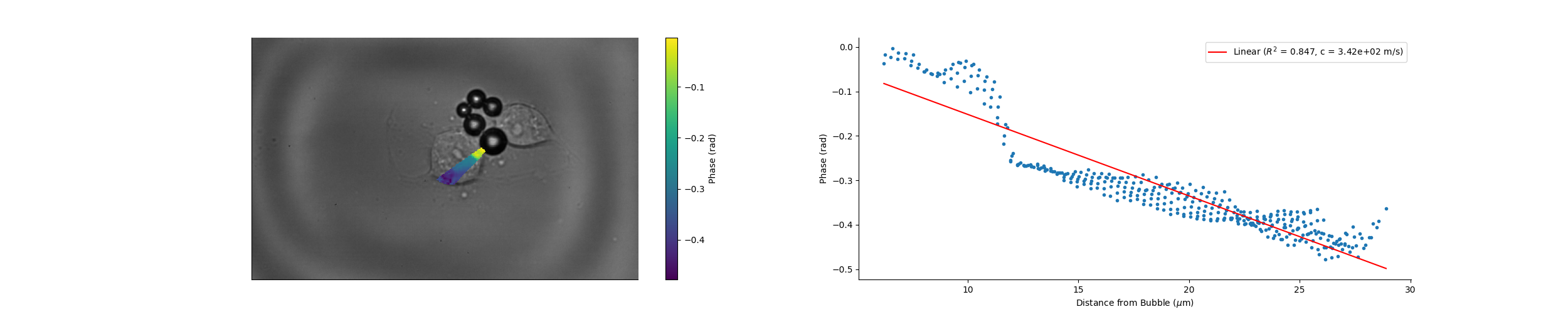

### Camera_14_49_16_Magnitude.png

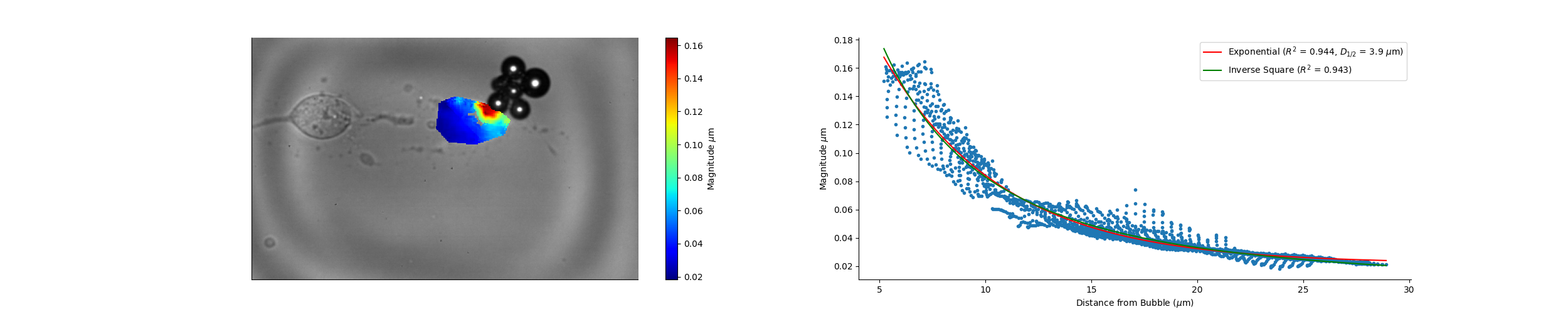

### Camera_14_49_16_Wave_Speed.png

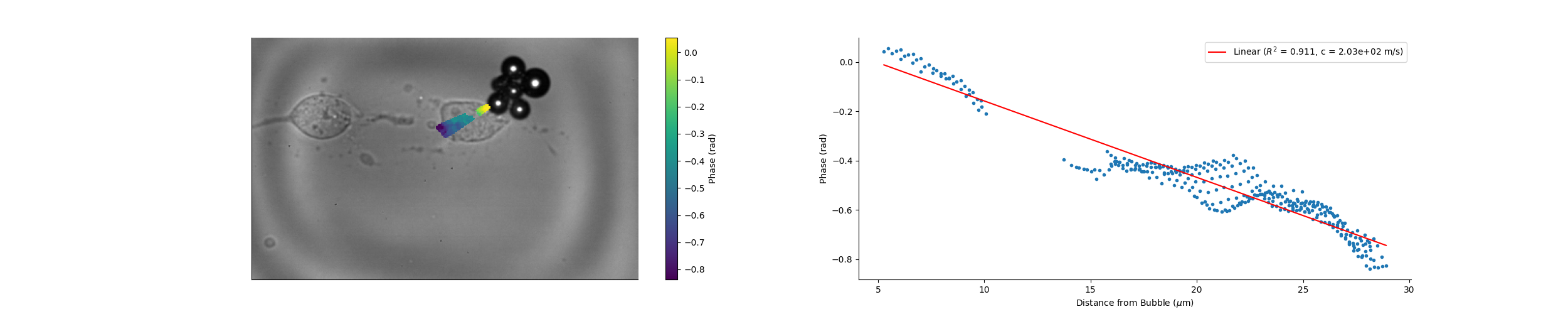

### Camera_14_52_16_Magnitude.png

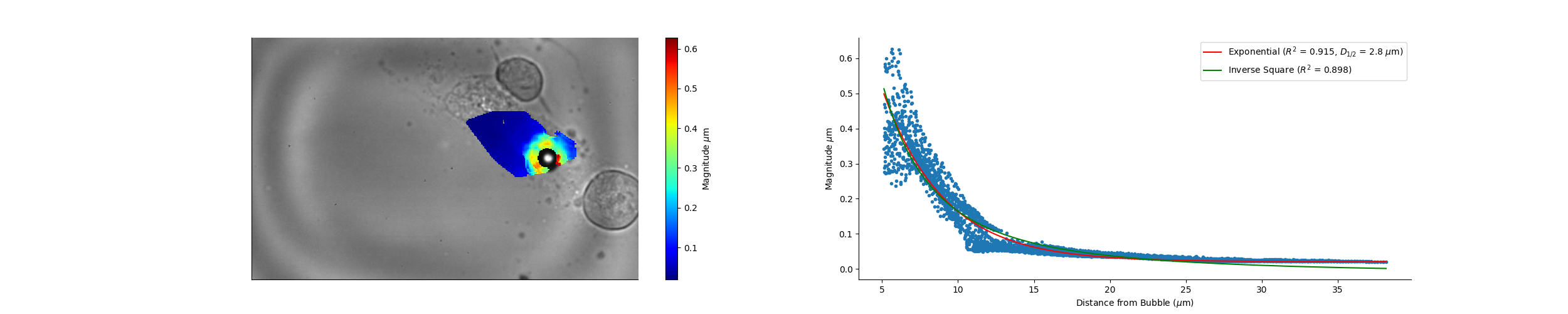

### Camera_14_52_16_Wave_Speed.png

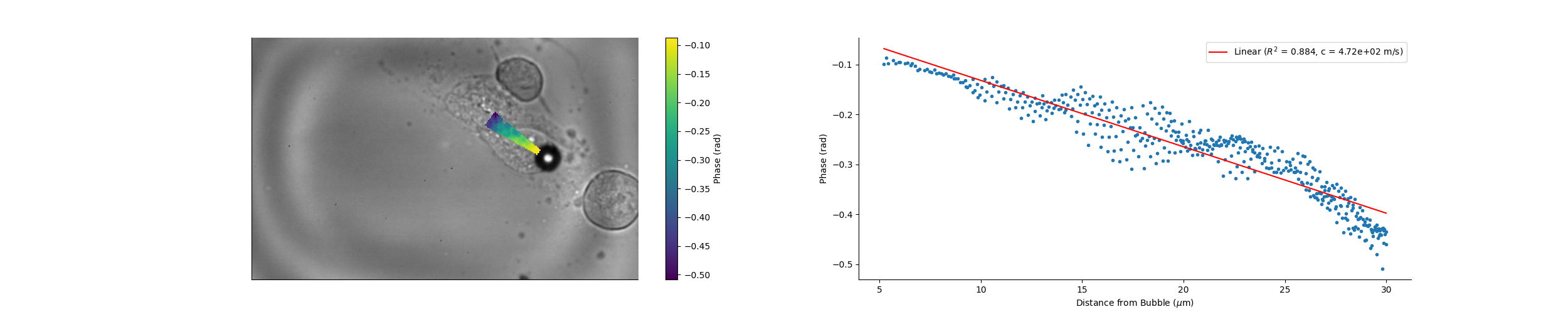

### Camera_14_56_53_Magnitude.png

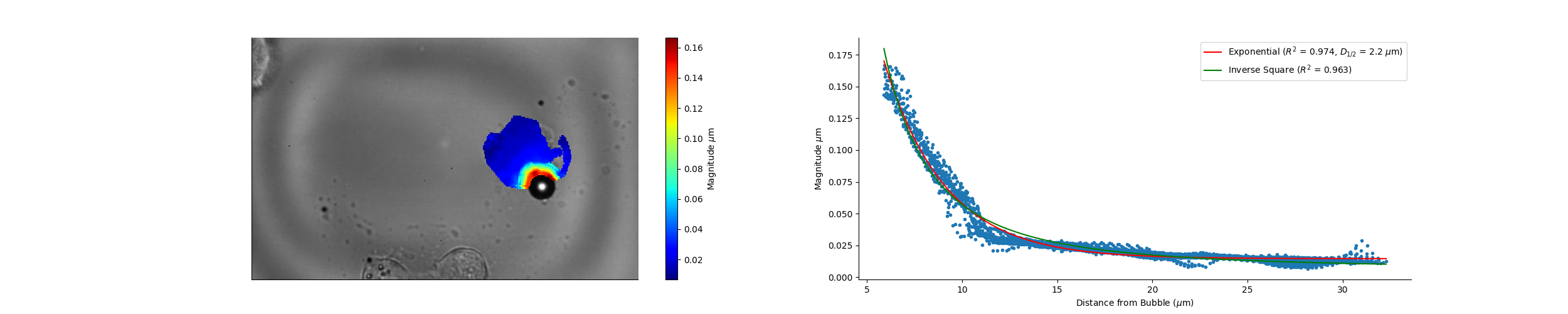

### Camera_14_56_53_Wave_Speed.png

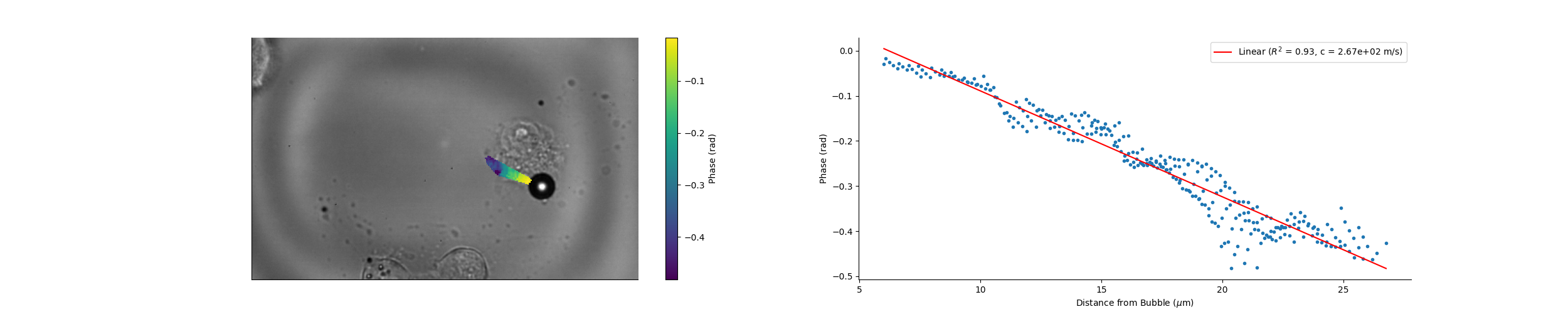

### Camera_14_58_15_Magnitude.png

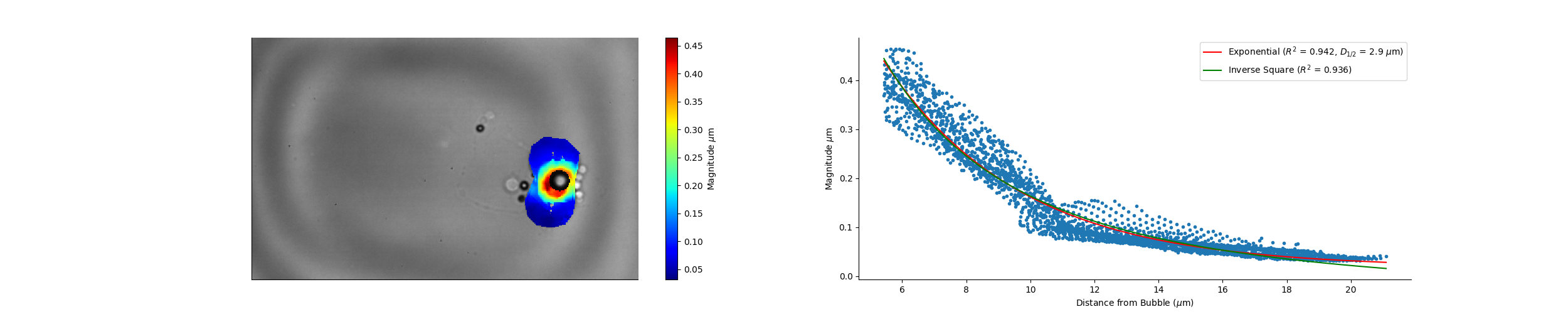

### Camera_14_59_04_Magnitude.png

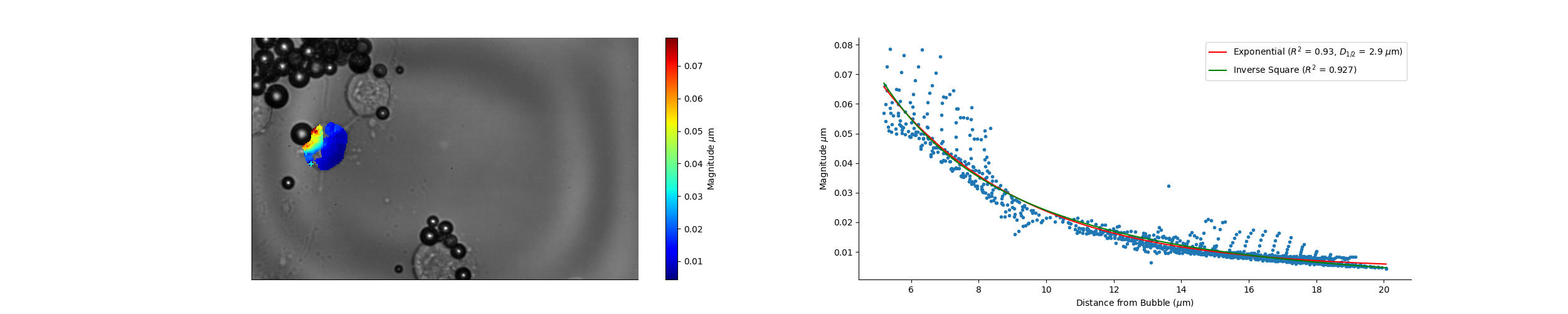

### Camera_15_00_28_Magnitude.png

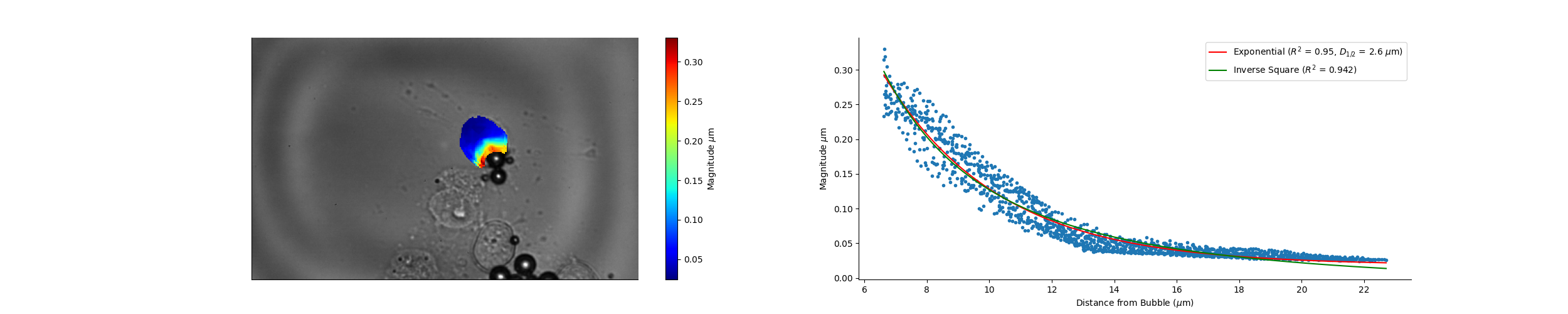

### Camera_15_08_03_Magnitude.png

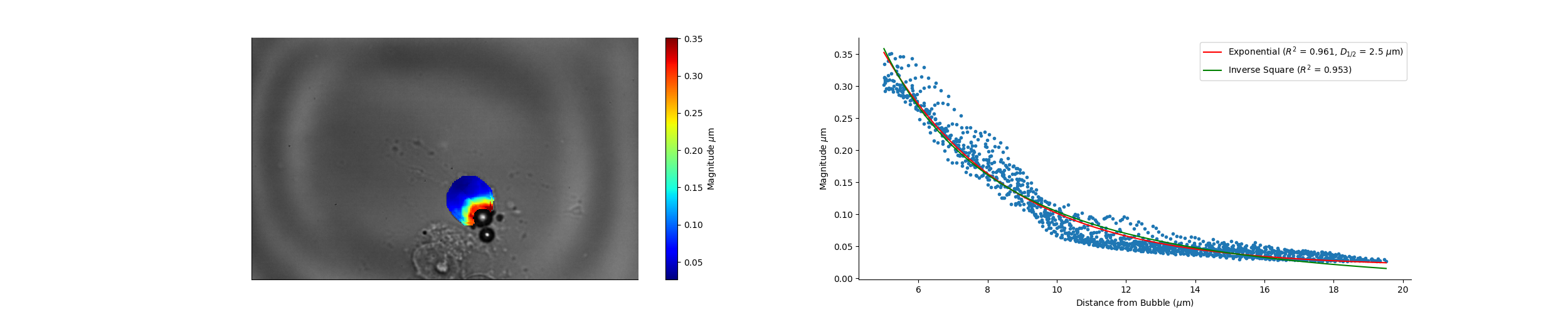

### Camera_15_09_57_Magnitude.png

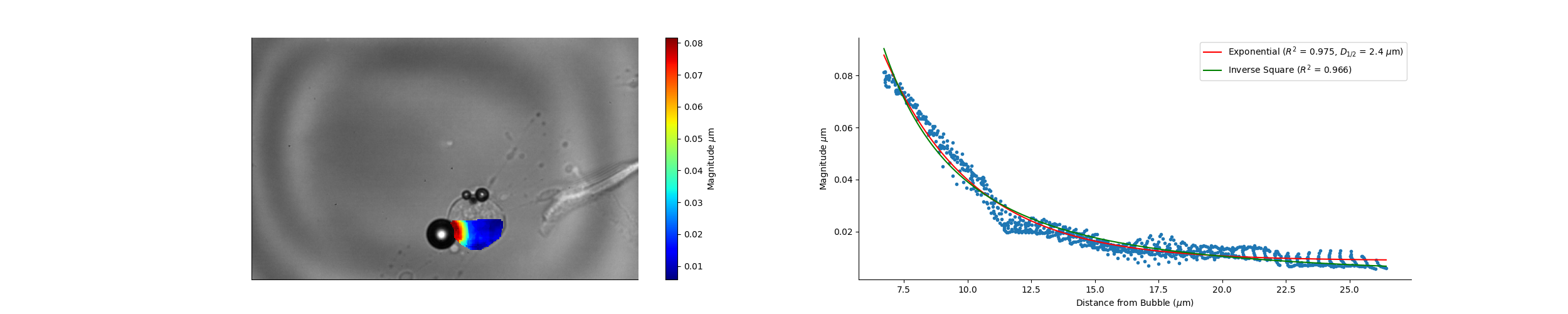

### Camera_15_09_57_Wave_Speed.png

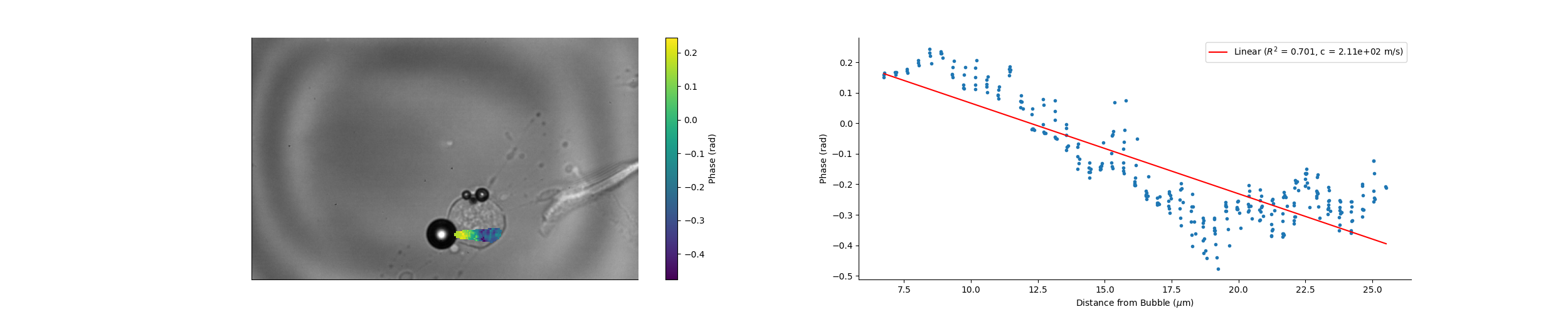

### Camera_16_18_20_Magnitude.png

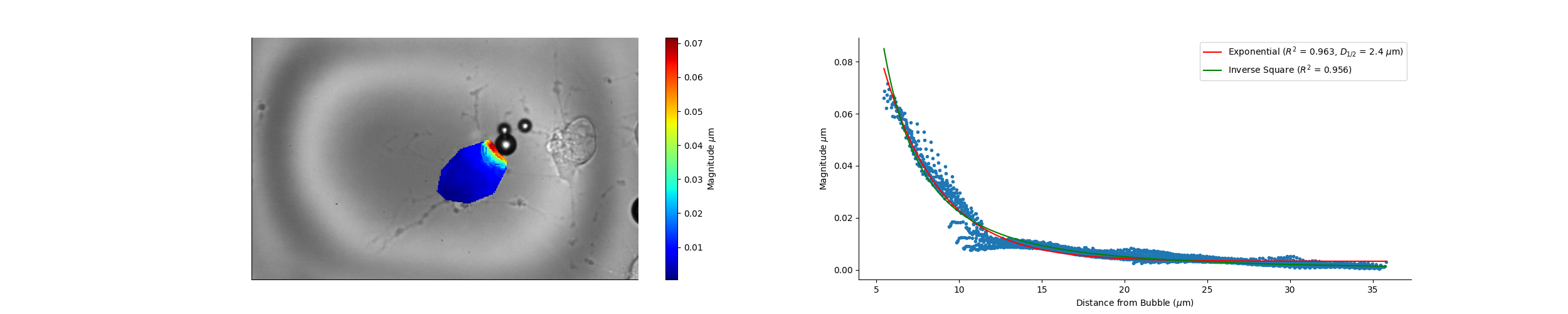

### Camera_16_18_20_Wave_Speed.png

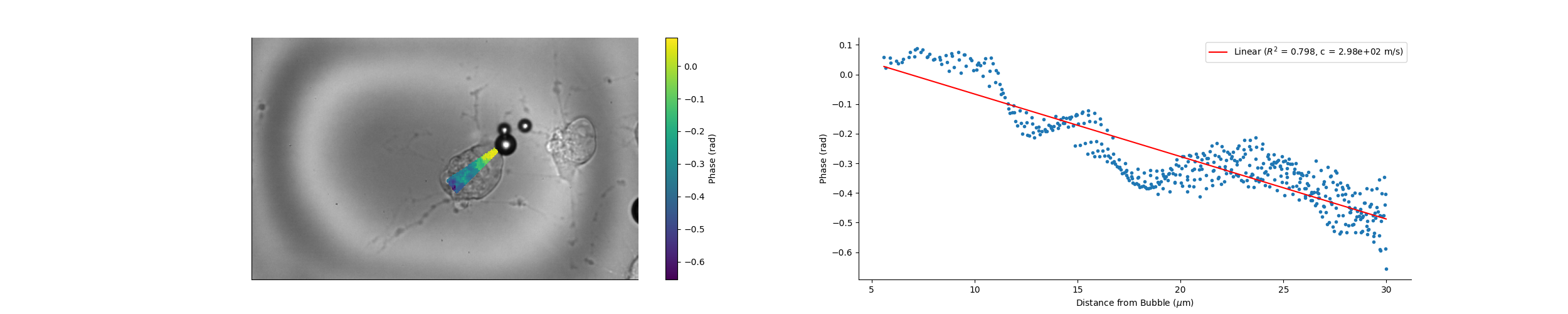

### Camera_16_29_56_Magnitude.png

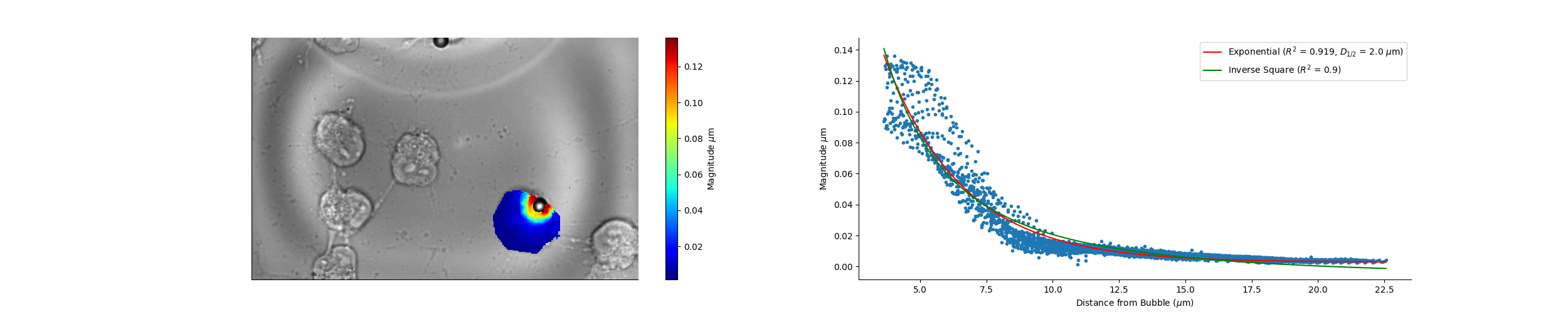

### Camera_16_30_41_Magnitude.png

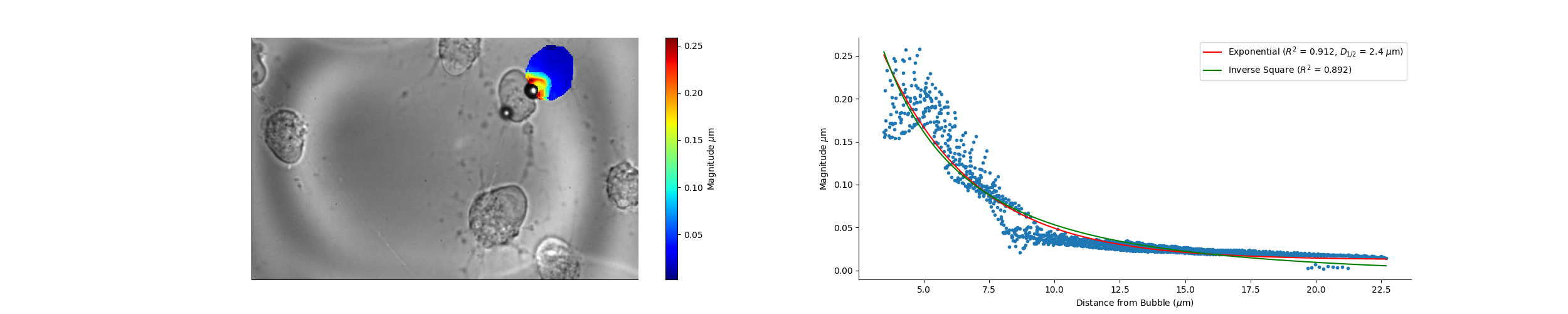

### Camera_16_33_22_Magnitude.png

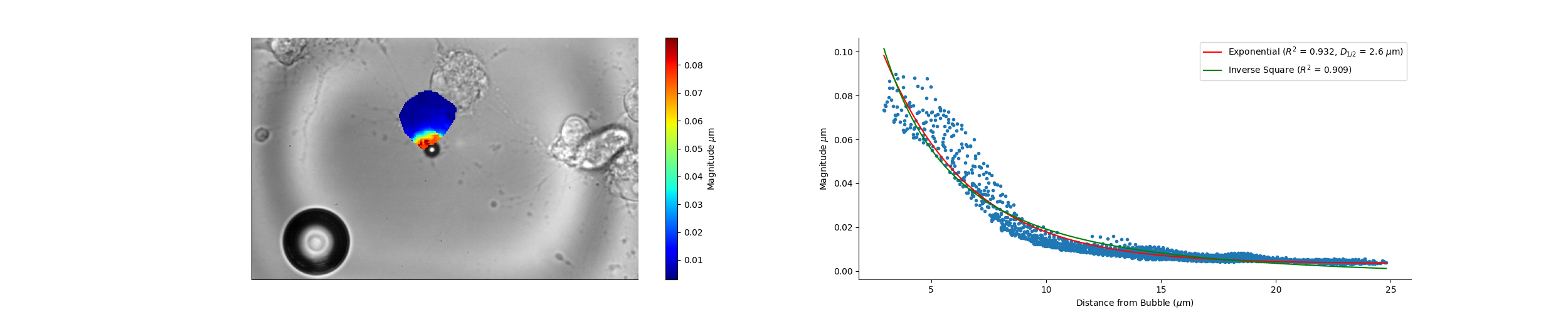

### Camera_16_33_22_Wave_Speed.png

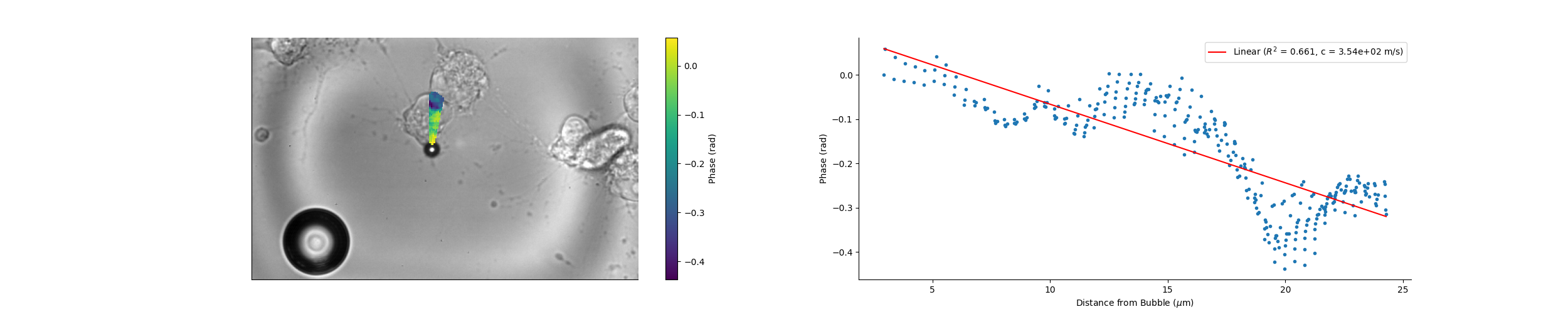

### Camera_16_33_32_Magnitude.png

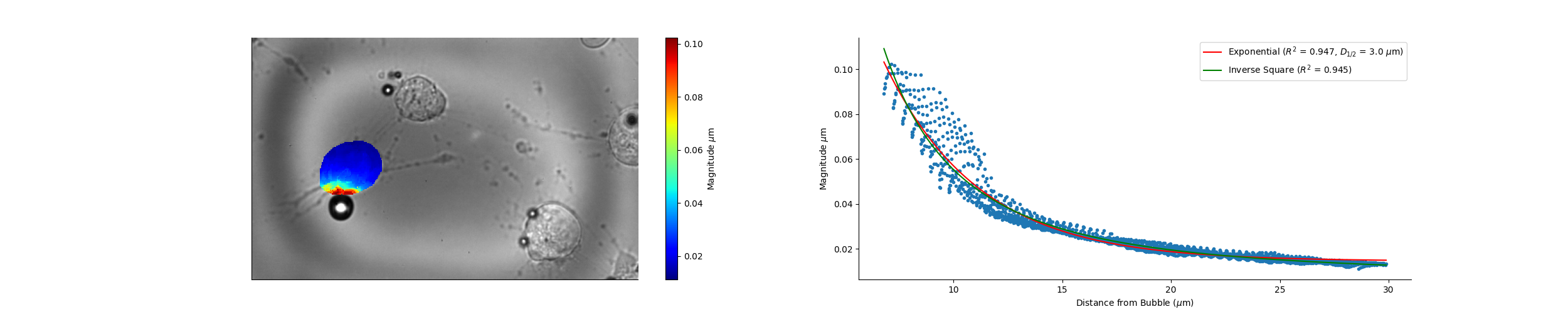

### Camera_16_33_32_Wave_Speed.png

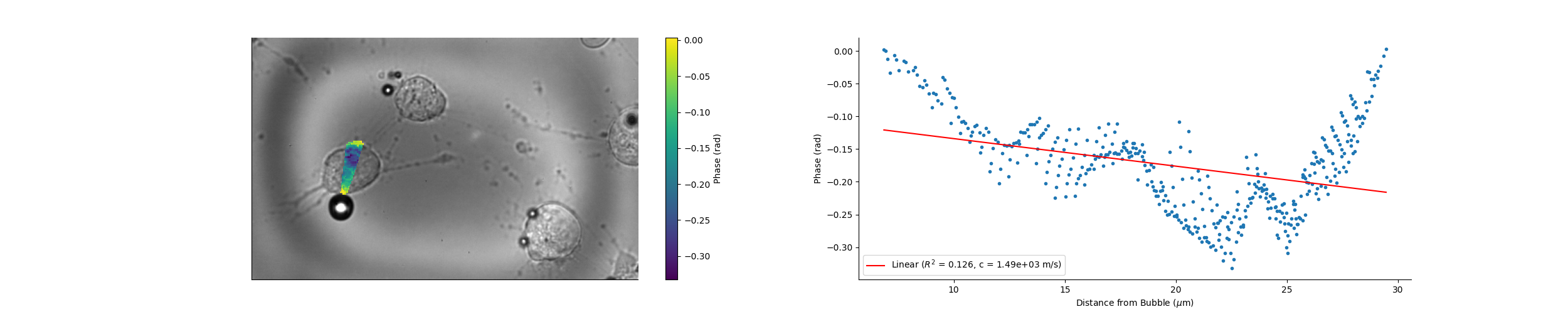

### Camera_16_36_40_Magnitude.png

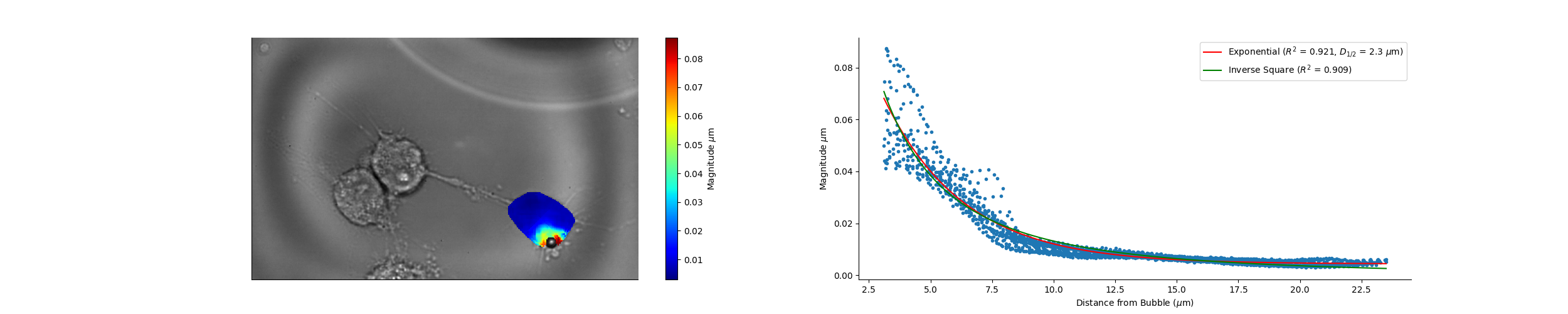

### Camera_16_38_18_Magnitude.png

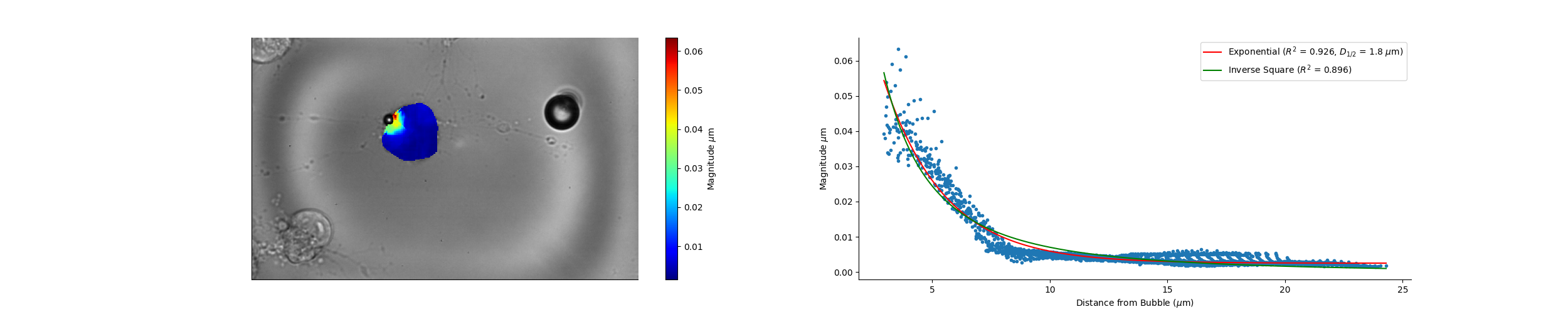

### Camera_16_43_05_Magnitude.png

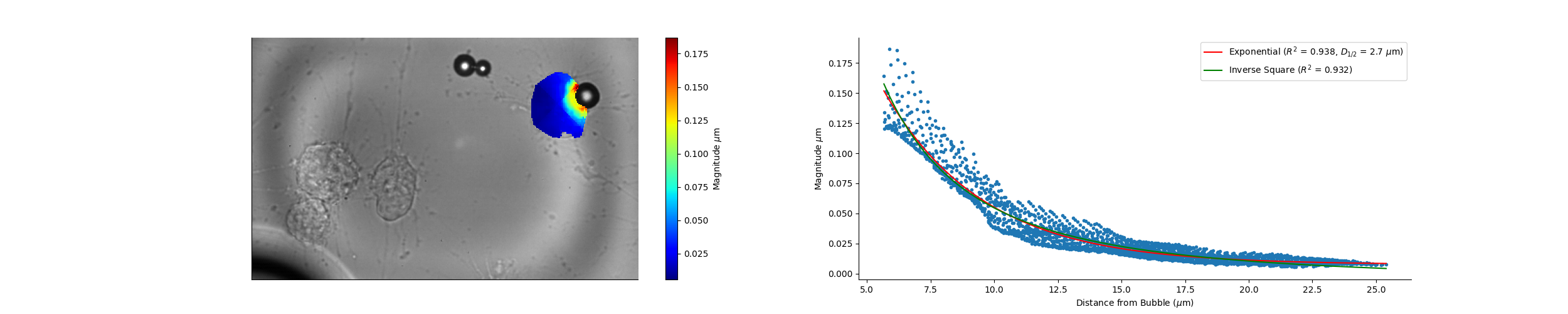

### Camera_16_45_03_Magnitude.png

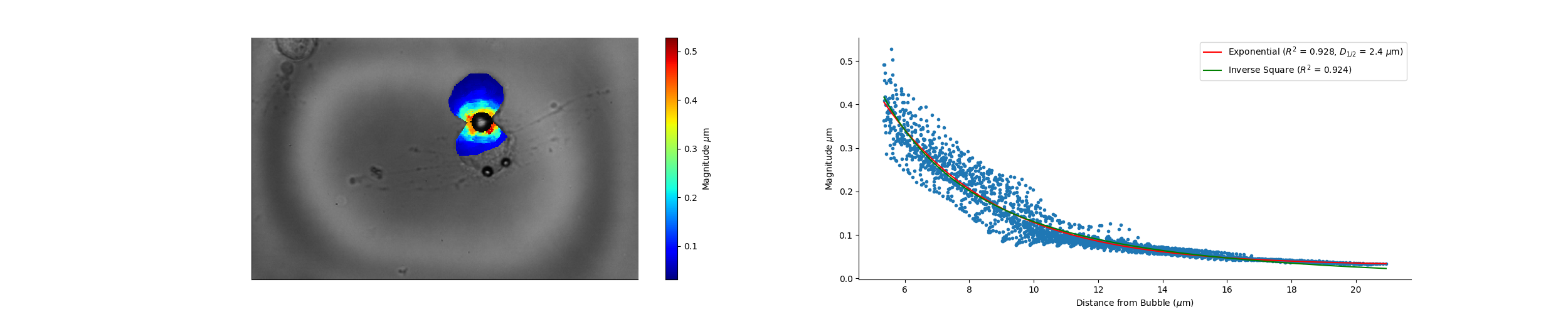

### Camera_16_56_27_Magnitude.png

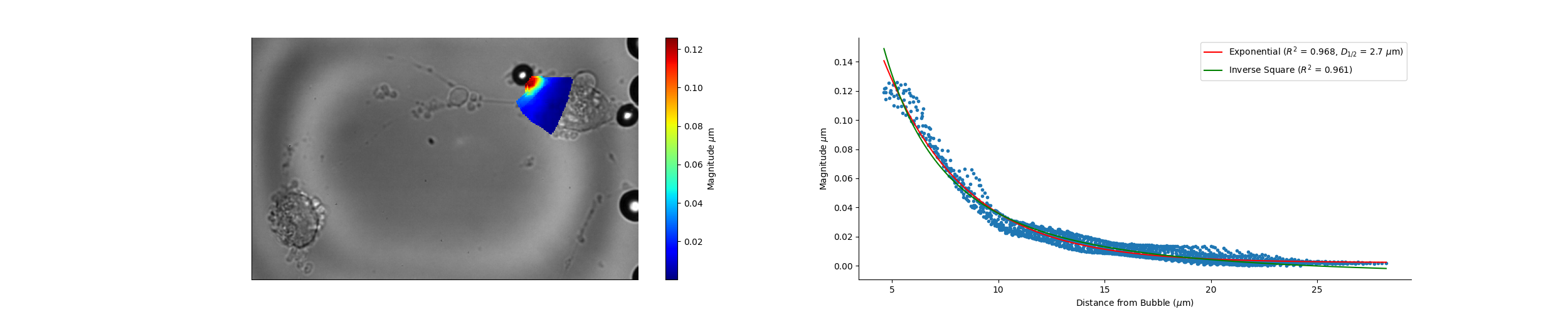

### Camera_16_57_16_Magnitude.png

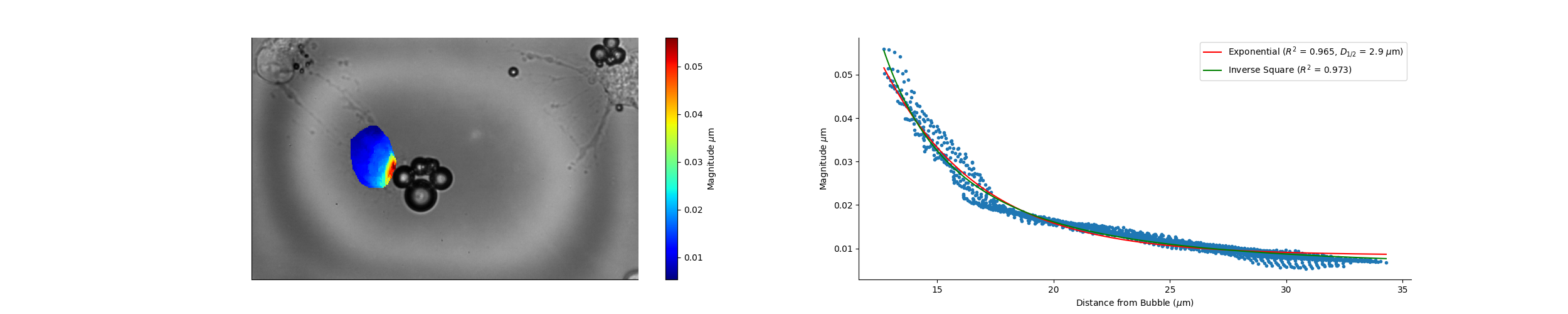

### Camera_16_57_40_Magnitude.png

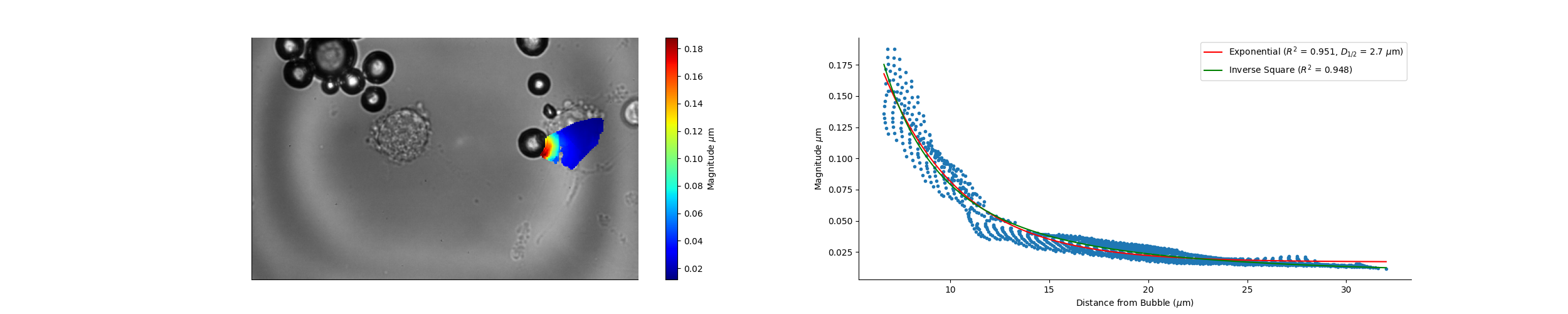

### Camera_16_58_49_Magnitude.png

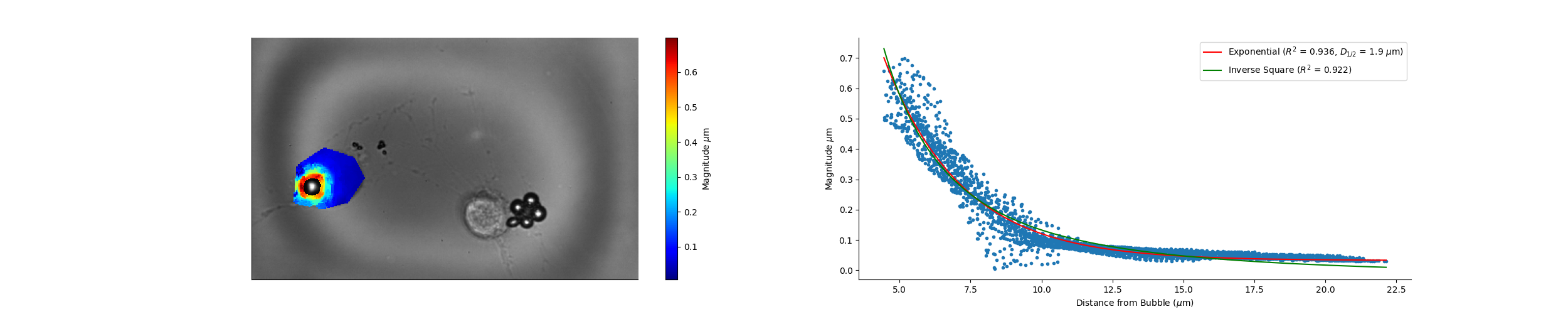

### Camera_17_02_49_Magnitude.png

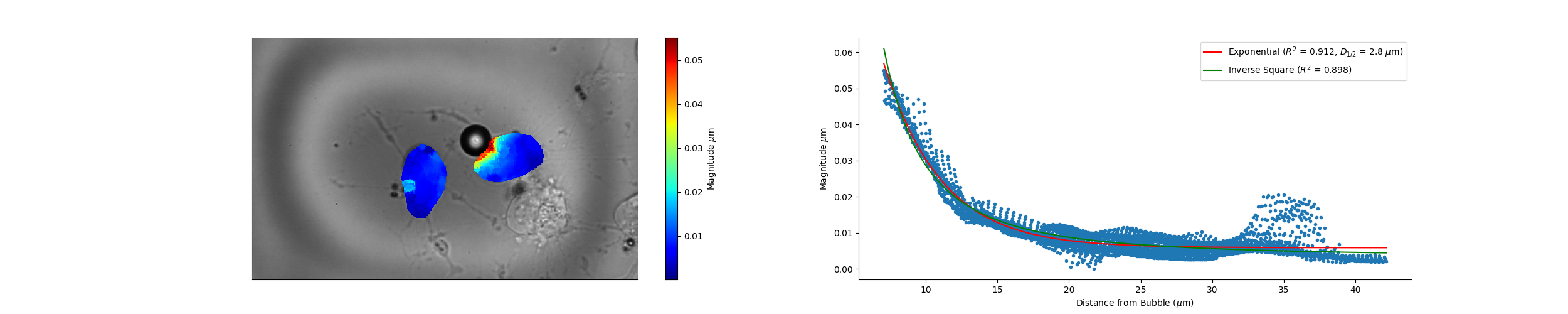

### Camera_17_15_54_Magnitude.png

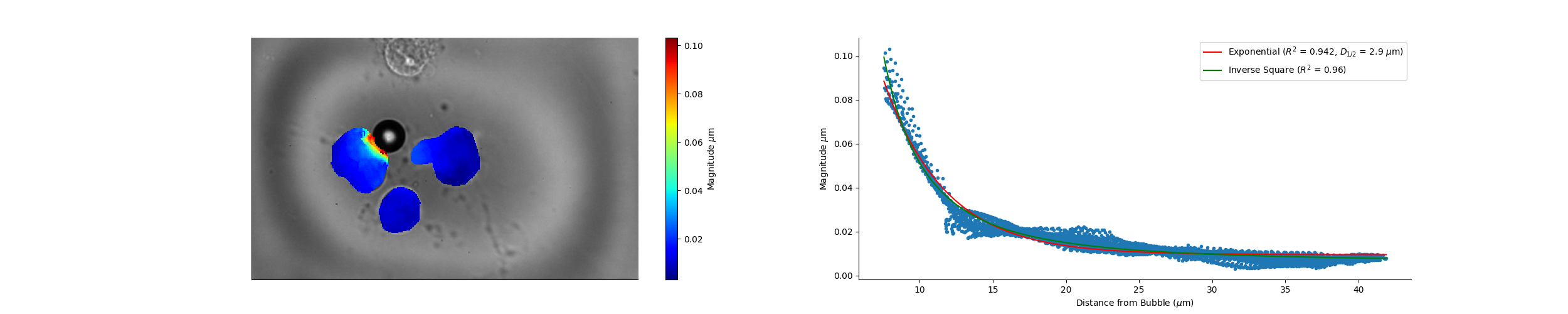

### Camera_17_17_18_Magnitude.png

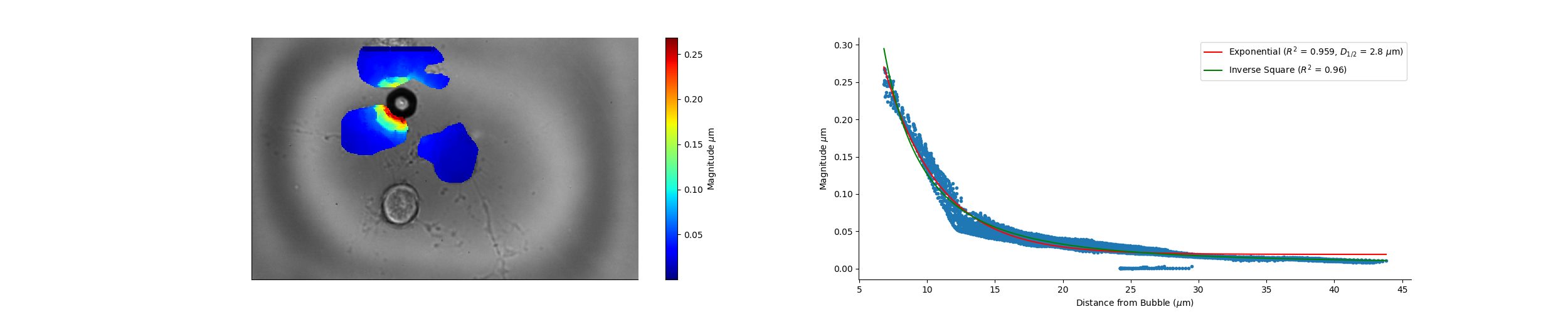

### Camera_17_21_43_Magnitude.png

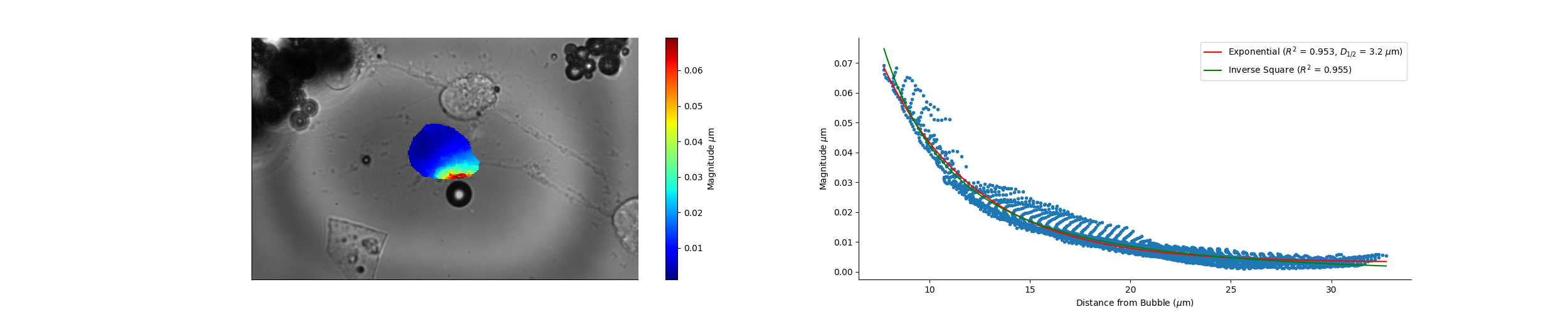

### Camera_17_33_52_Magnitude.png

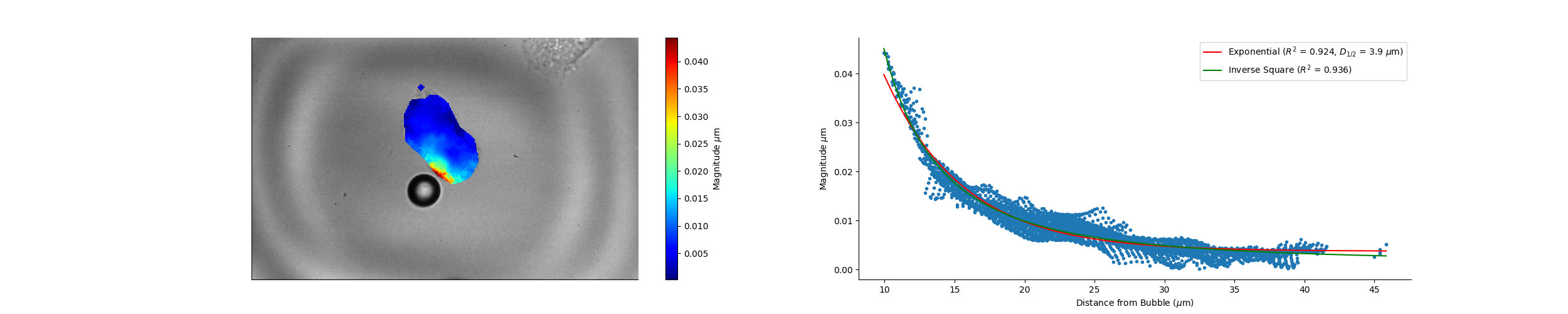

### Camera_17_33_52_Wave_Speed.png

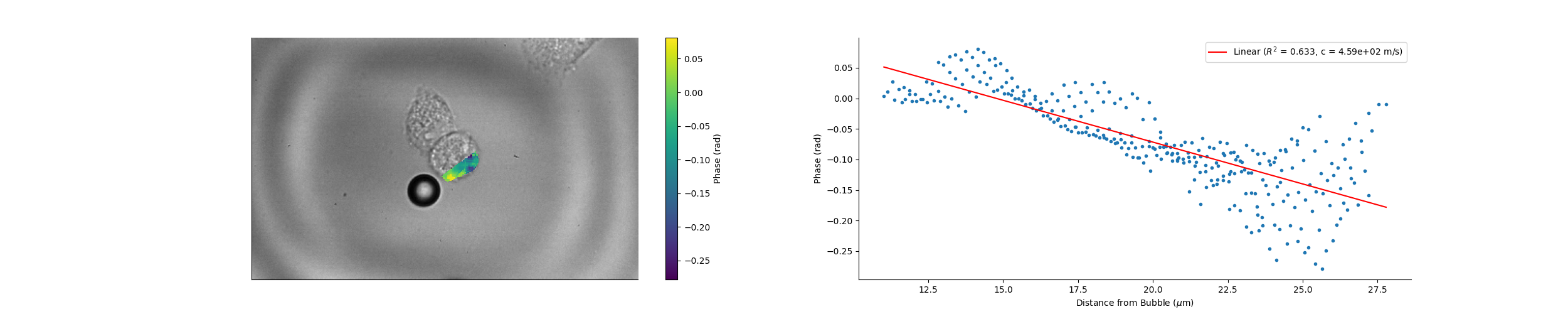

### Camera_17_39_33_Magnitude.png

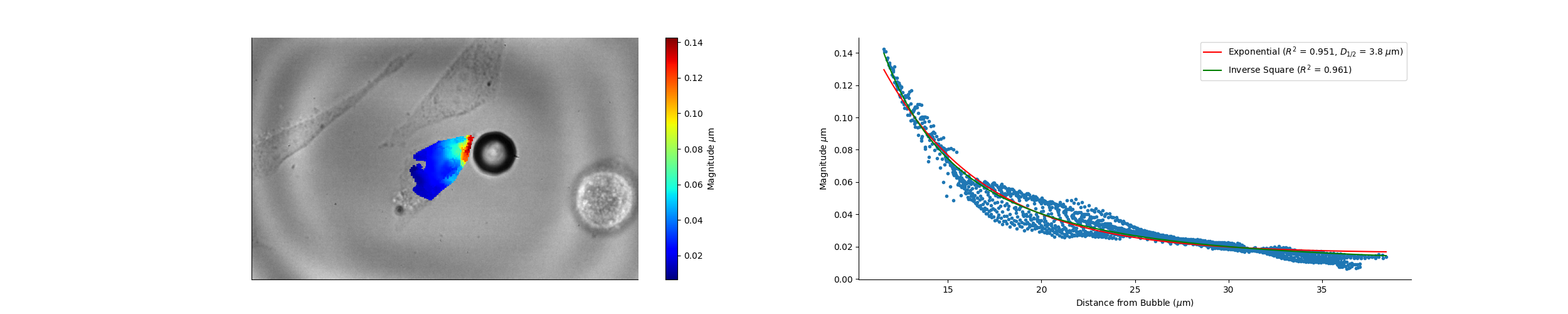

### Camera_17_39_33_Wave_Speed.png

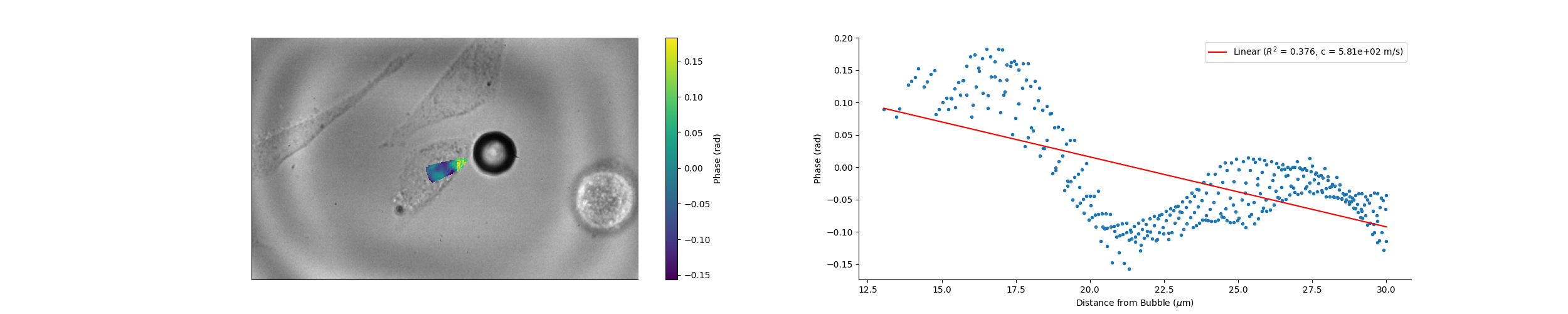

### Camera_17_41_54_Magnitude.png

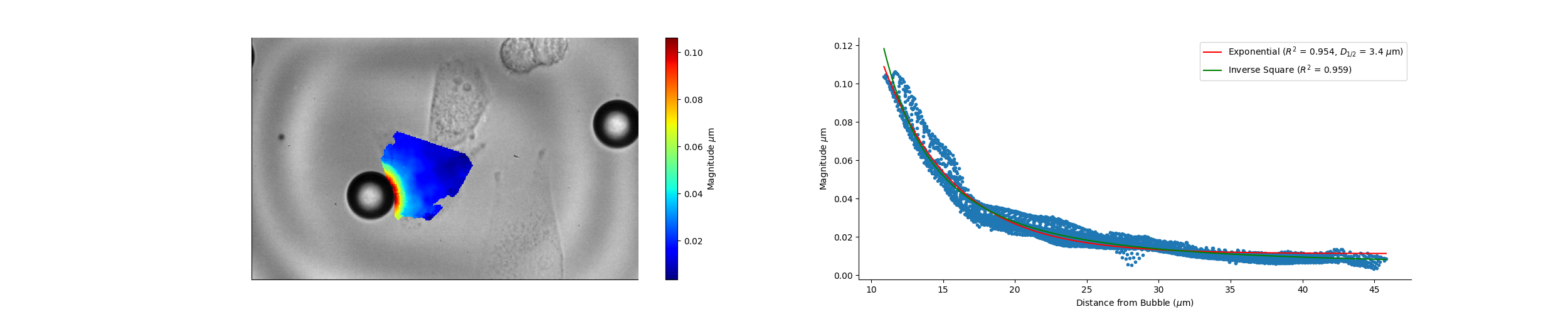

### Camera_17_41_54_Wave_Speed.png

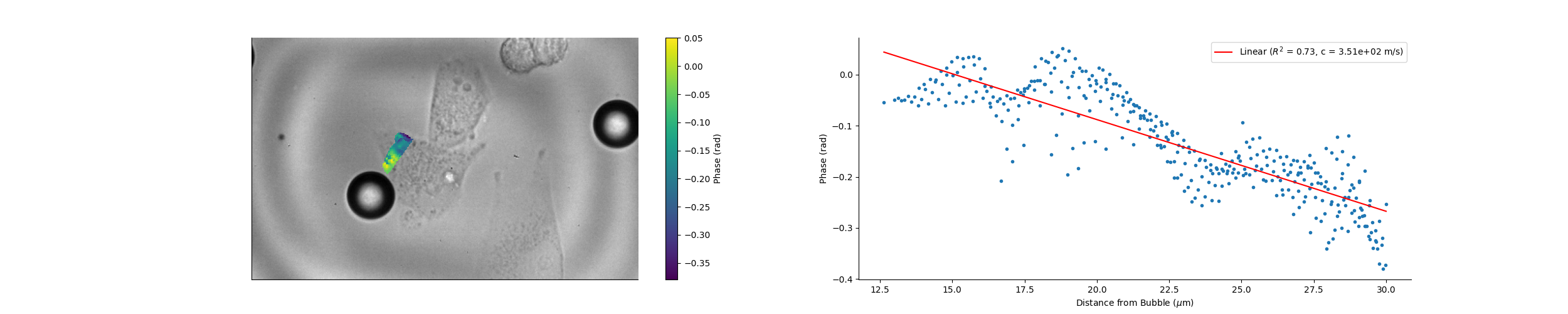

### Camera_17_42_55_Magnitude.png

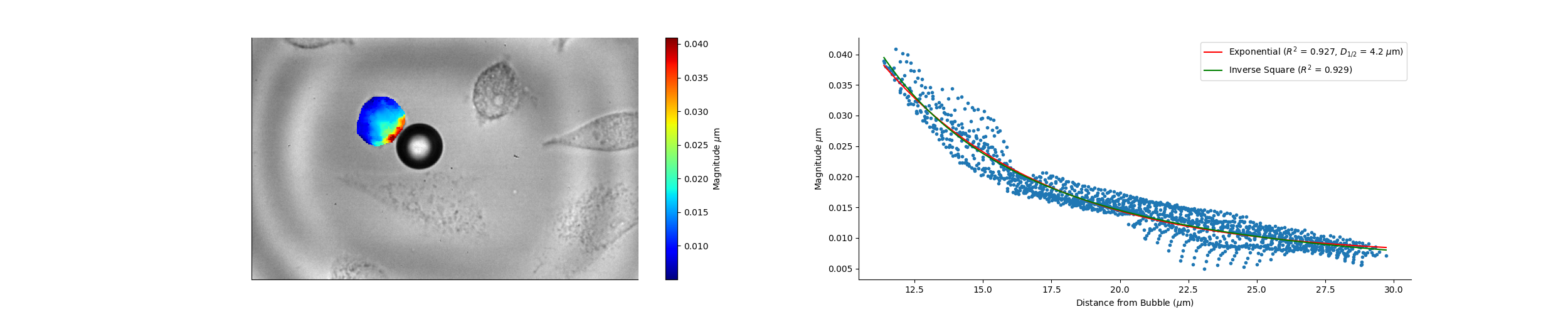

### Camera_17_42_55_Wave_Speed.png

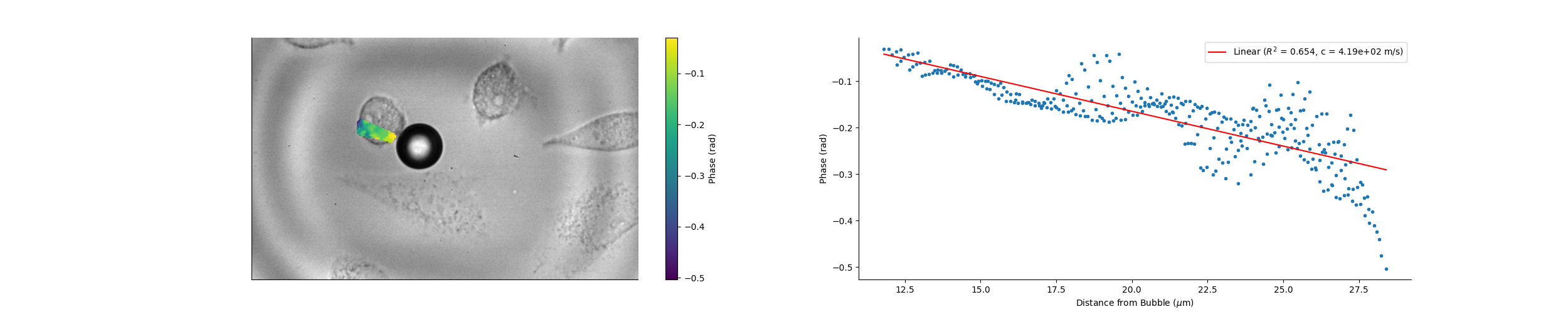

### Camera_17_43_52_Magnitude.png

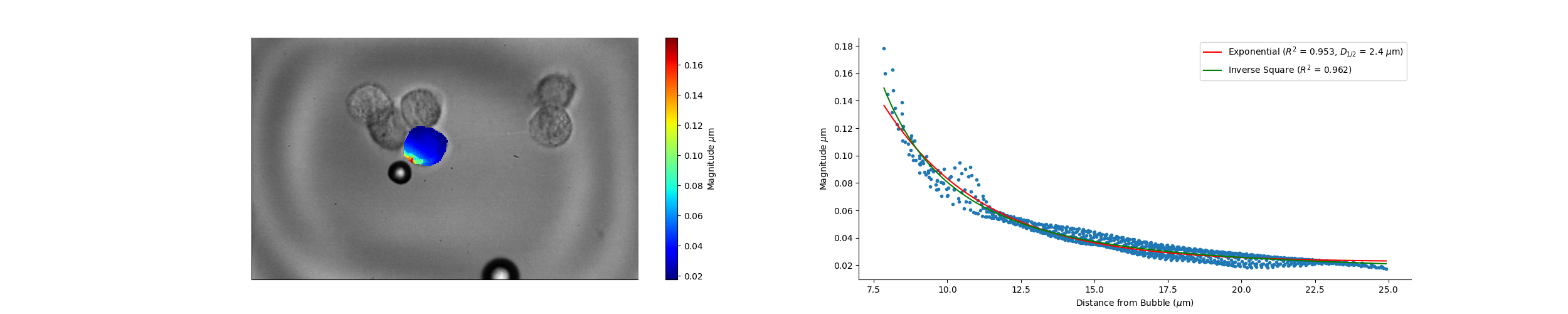

### Camera_17_43_52_Wave_Speed.png

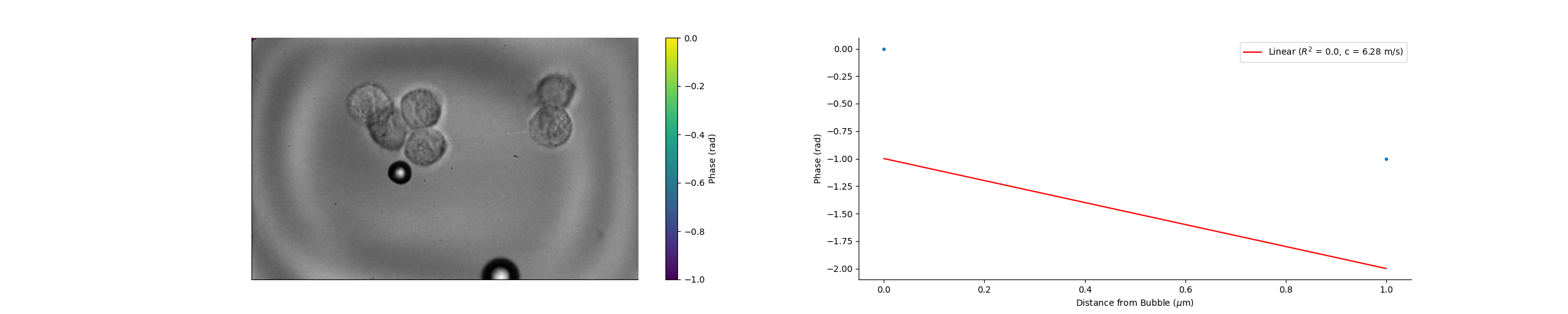

### Camera_17_45_13_Magnitude.png

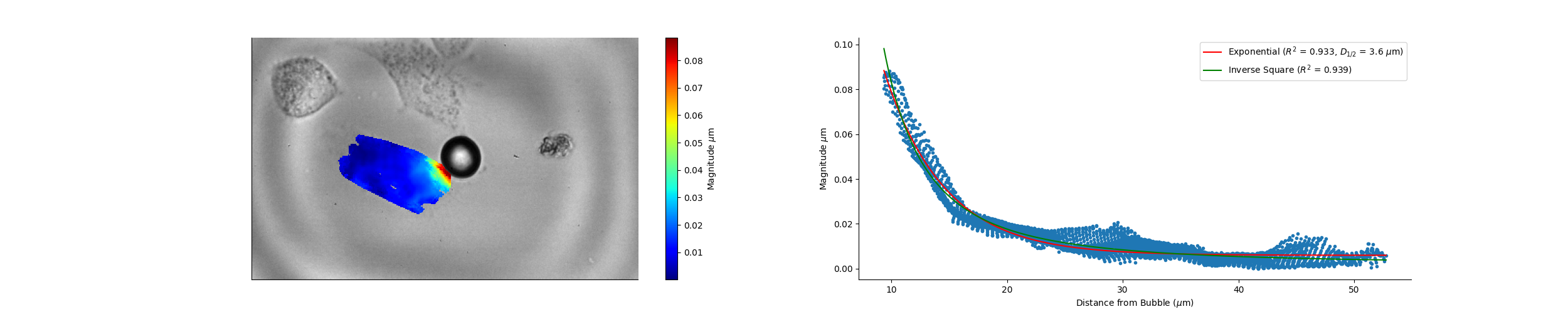

### Camera_17_45_13_Wave_Speed.png

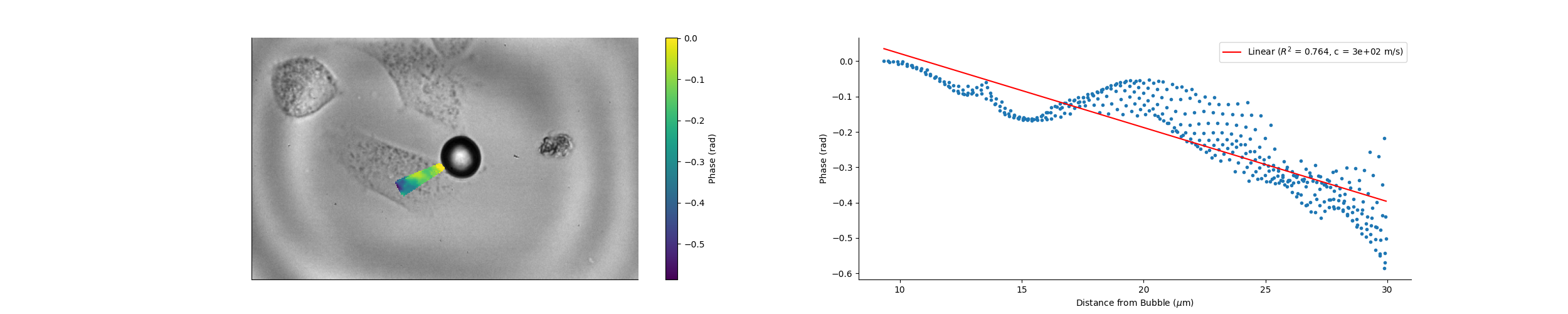

### Camera_17_45_52_Magnitude.png

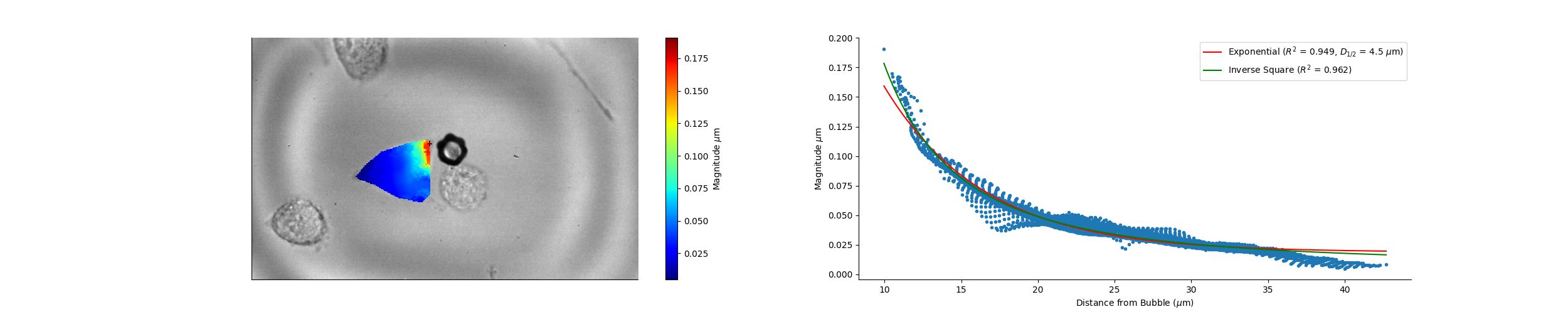

### Camera_17_45_52_Wave_Speed.png

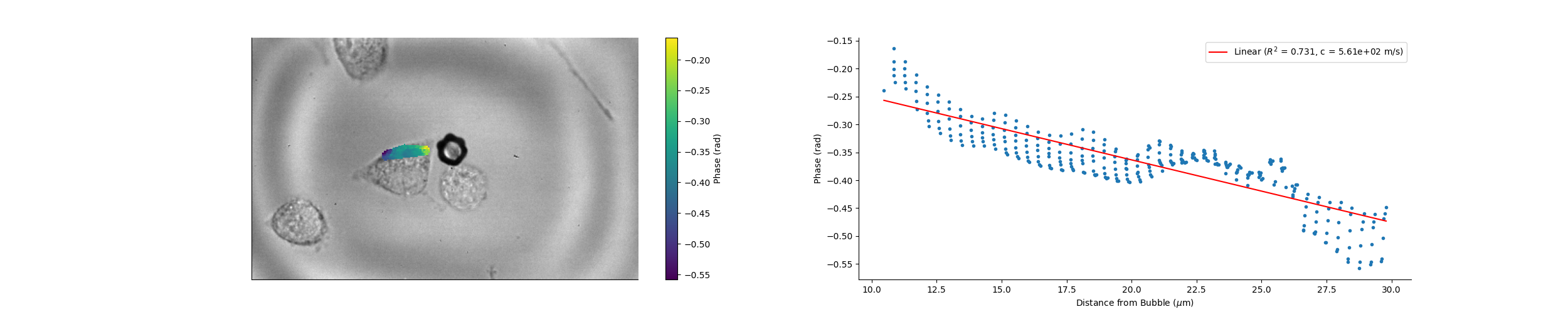

### Camera_17_46_17_Magnitude.png

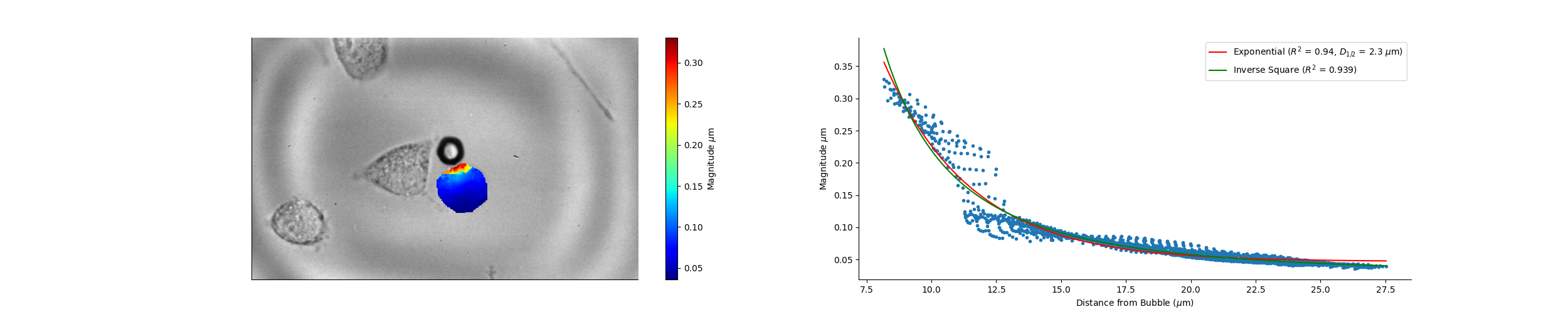

### Camera_17_46_17_Wave_Speed.png

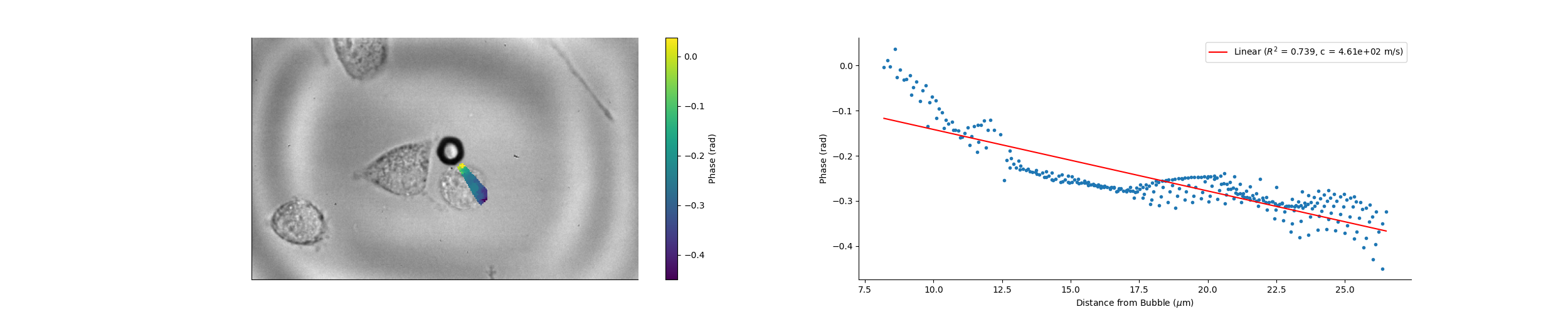

### Camera_17_47_43_Magnitude.png

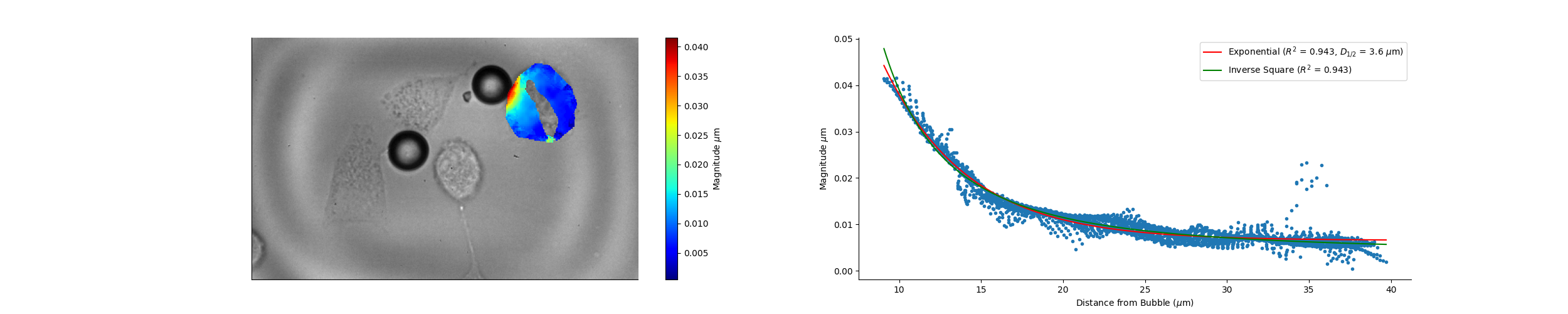

### Camera_17_47_43_Wave_Speed.png

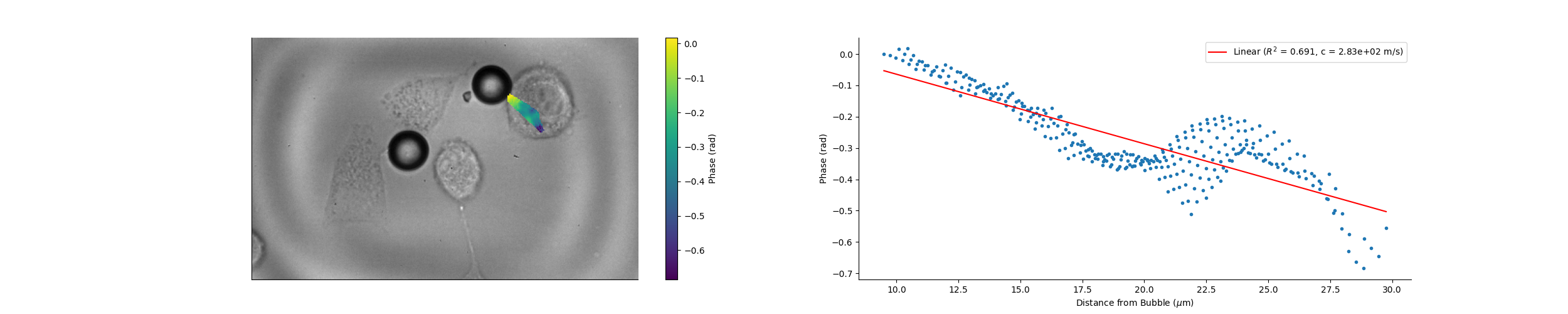

### Camera_17_50_39_Magnitude.png

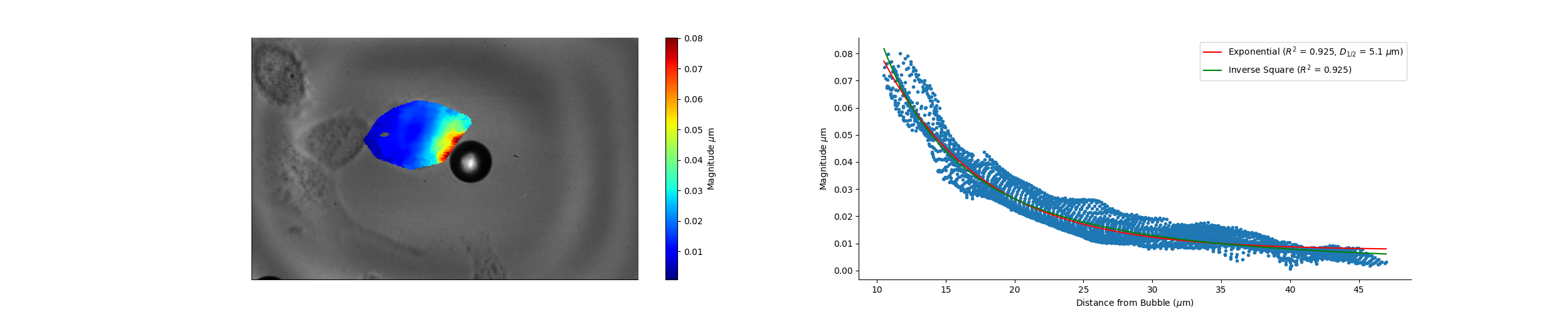

### Camera_17_50_39_Wave_Speed.png

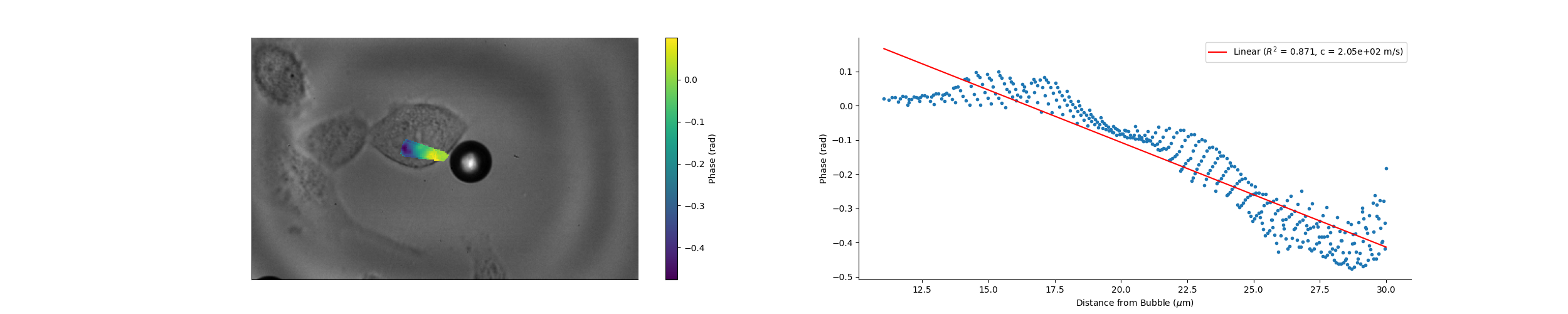

### Camera_17_51_16_Magnitude.png

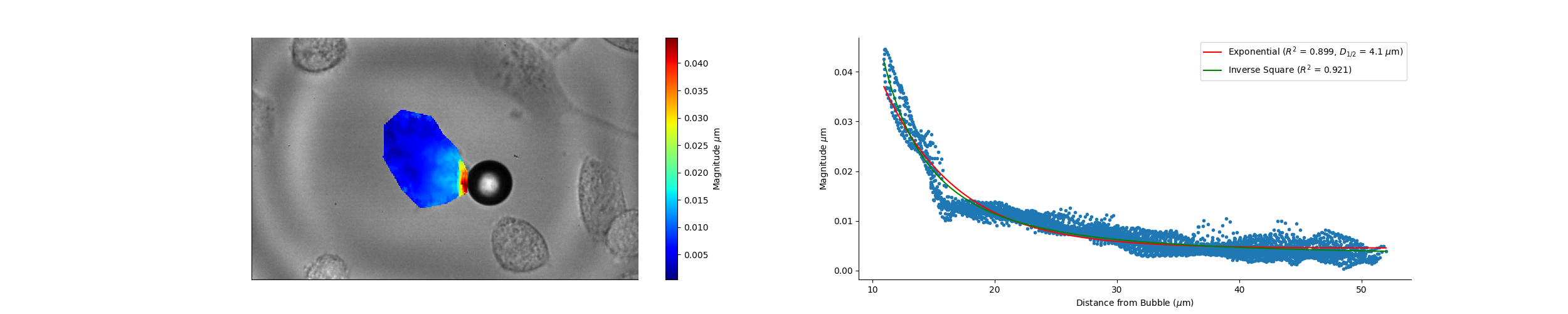

### Camera_17_51_16_Wave_Speed.png

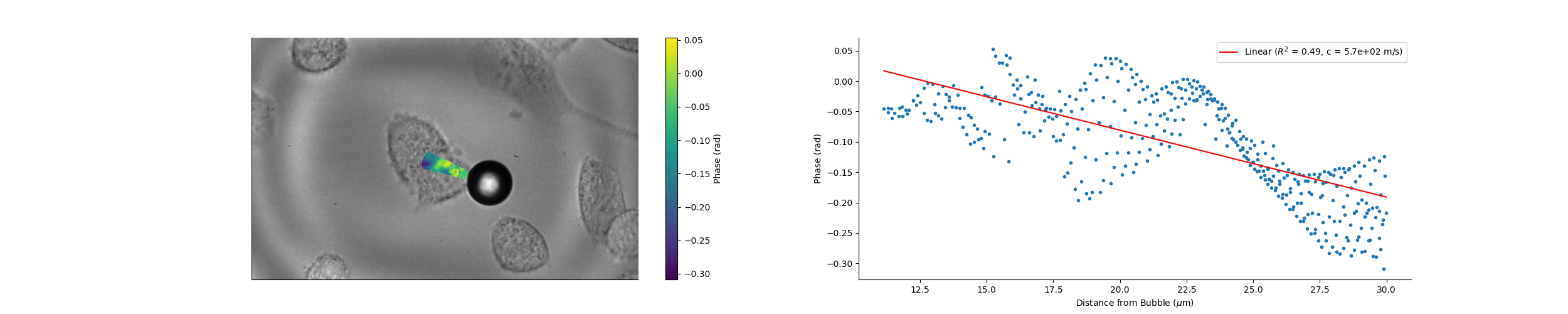

### Camera_17_58_29_Wave_Speed.png

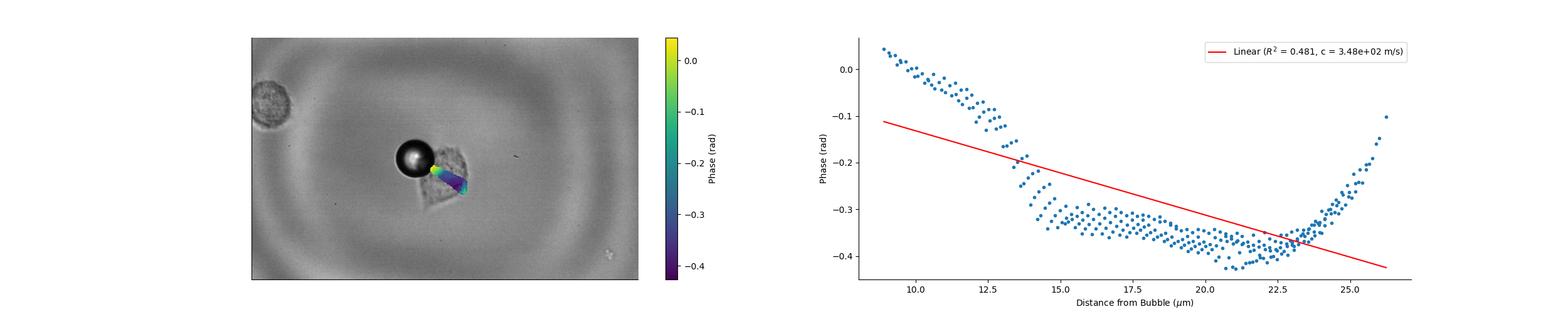

### Camera_17_58_57_Magnitude.png

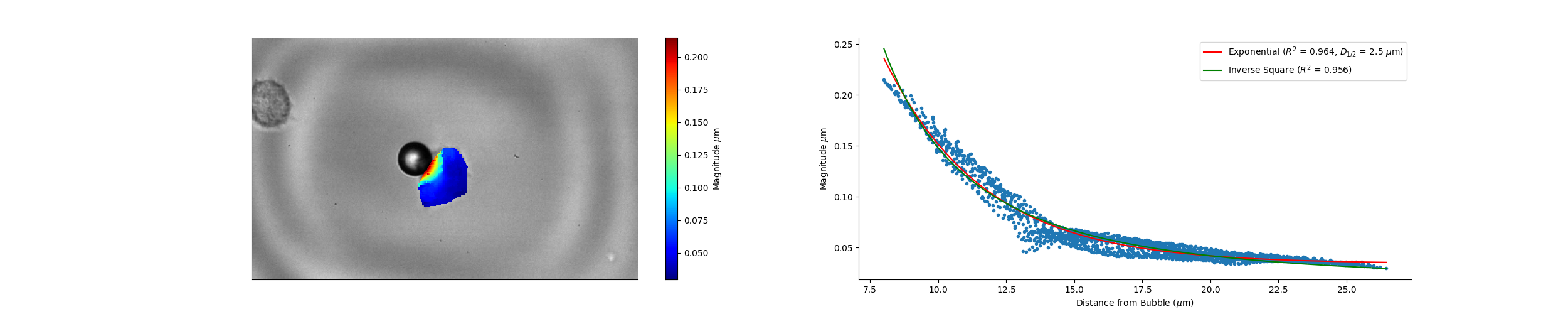

### Camera_18_15_53_Magnitude.png

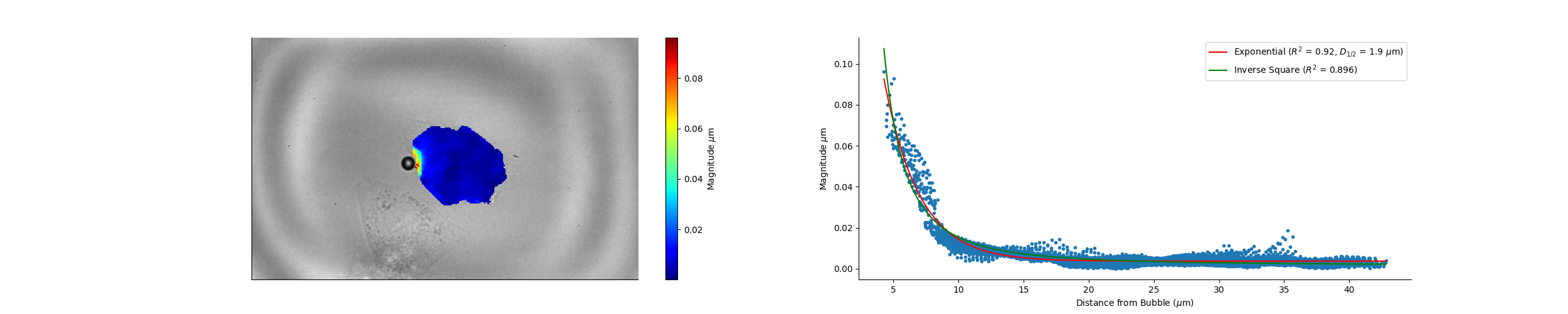

### Camera_18_15_53_Wave_Speed.png

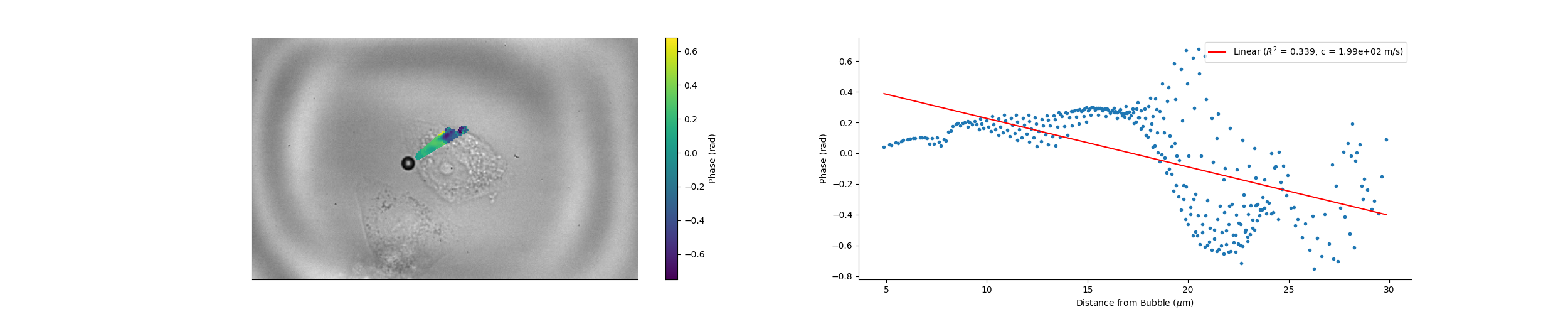

### Camera_18_16_29_Magnitude.png

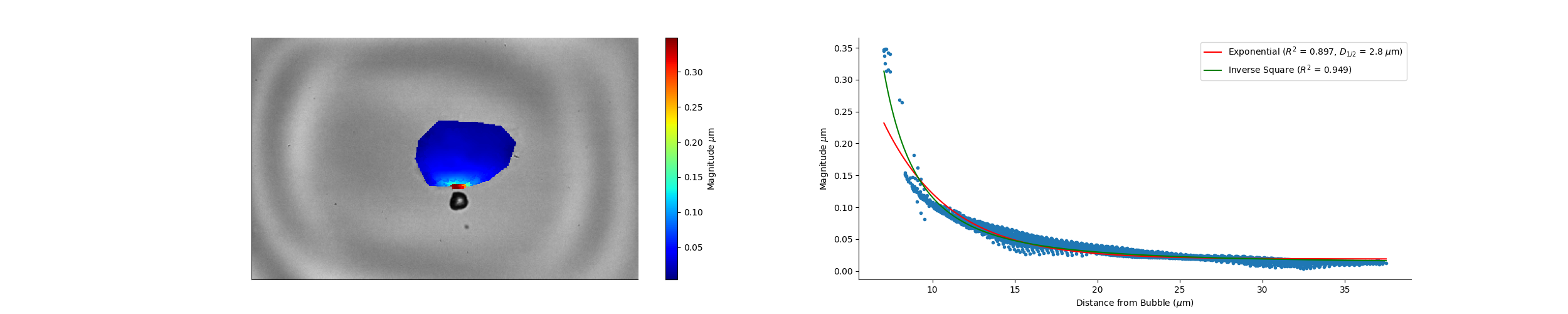

### Camera_18_16_29_Wave_Speed.png

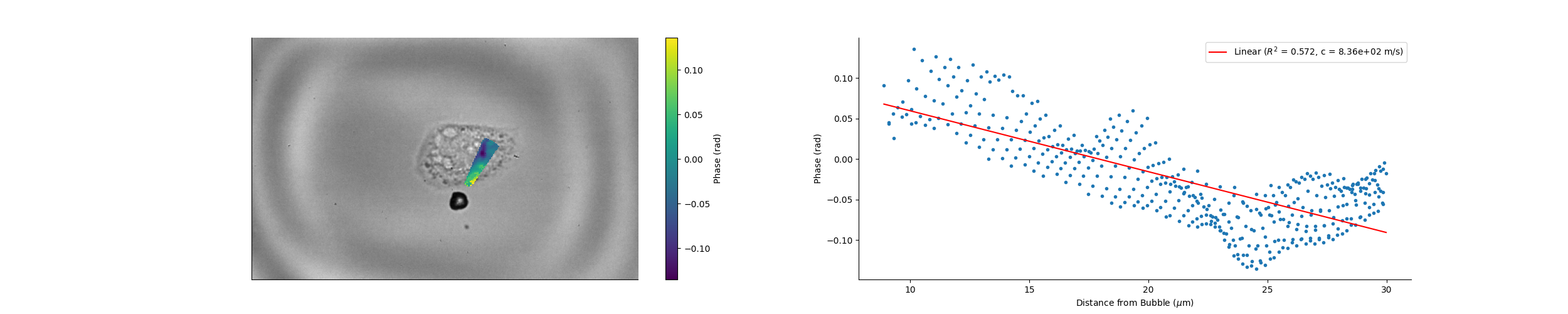

### Camera_18_18_00_Magnitude.png

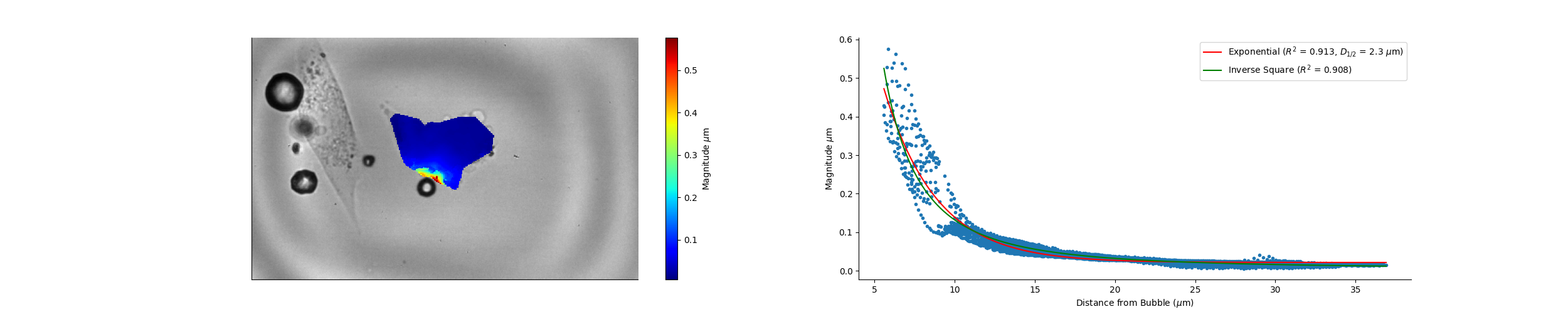

### Camera_18_18_00_Wave_Speed.png

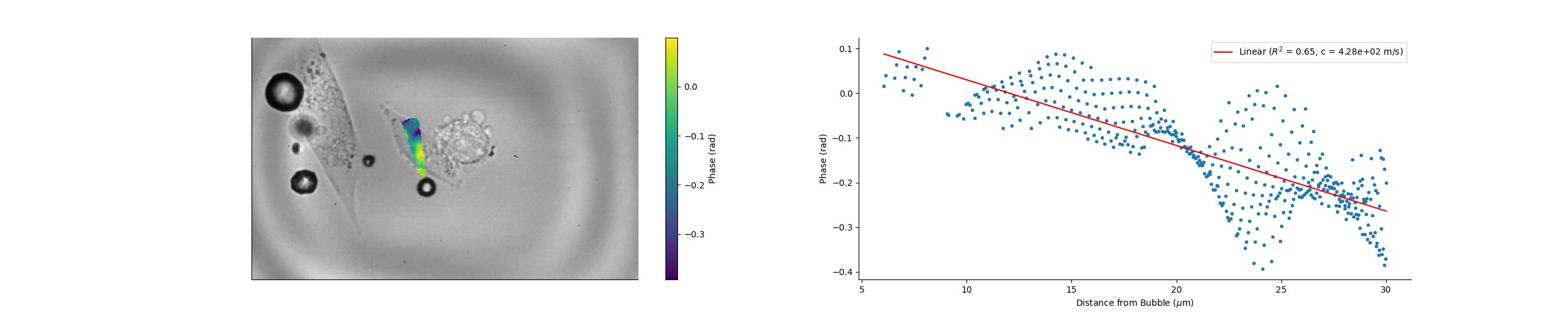

### Camera_18_18_41_Magnitude.png

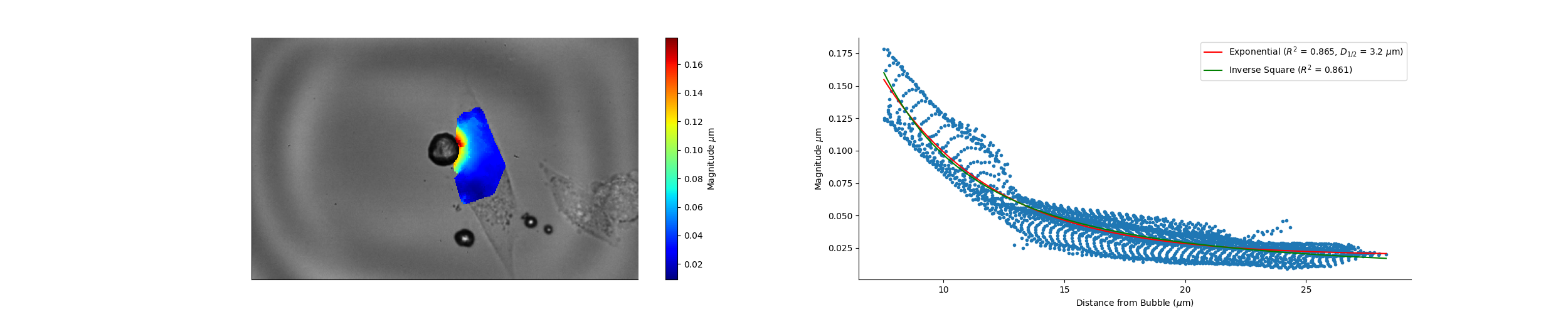

### Camera_18_18_41_Wave_Speed.png

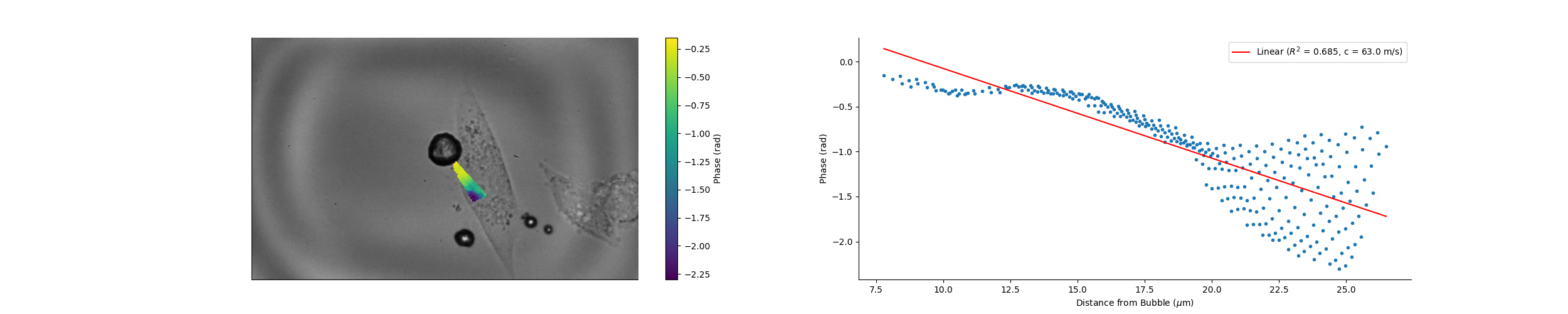

### Camera_18_19_08_Magnitude.png

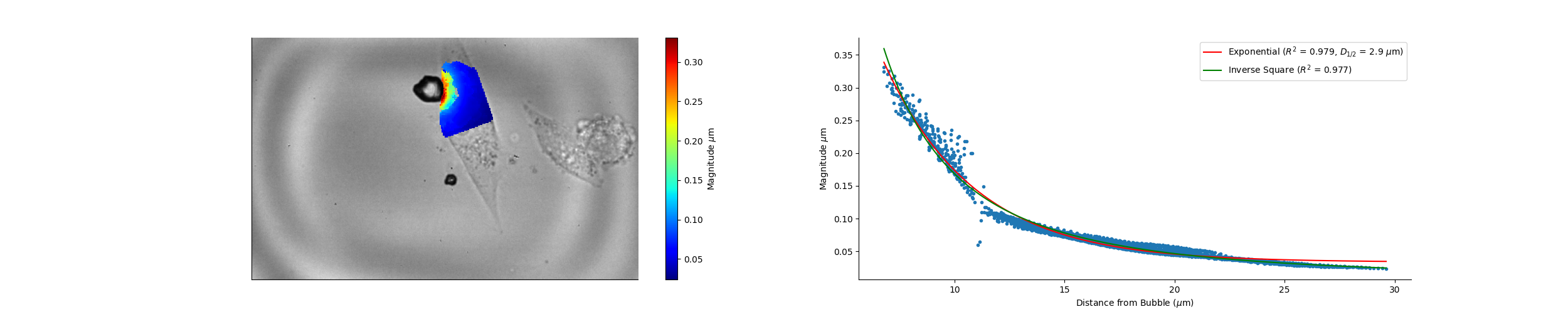

### Camera_18_19_08_Wave_Speed.png

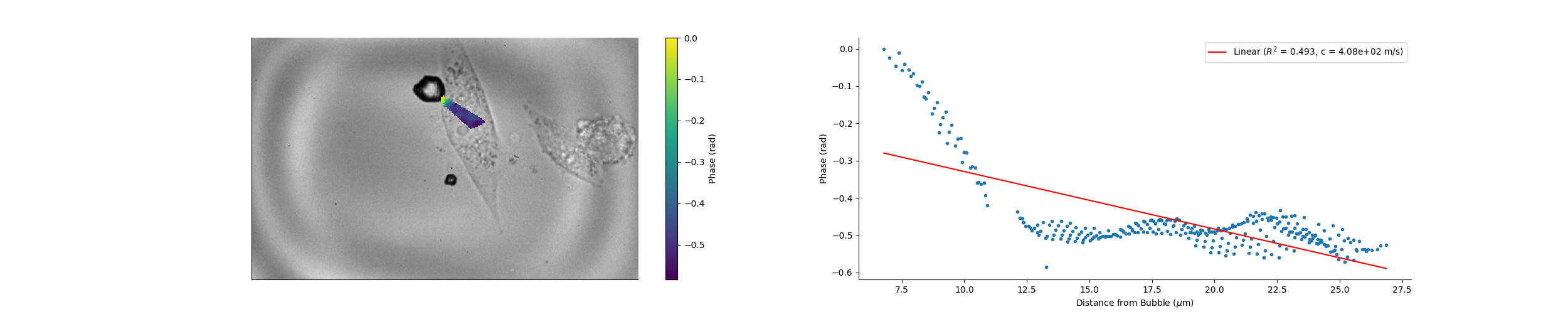

### Camera_18_56_39_Magnitude.png

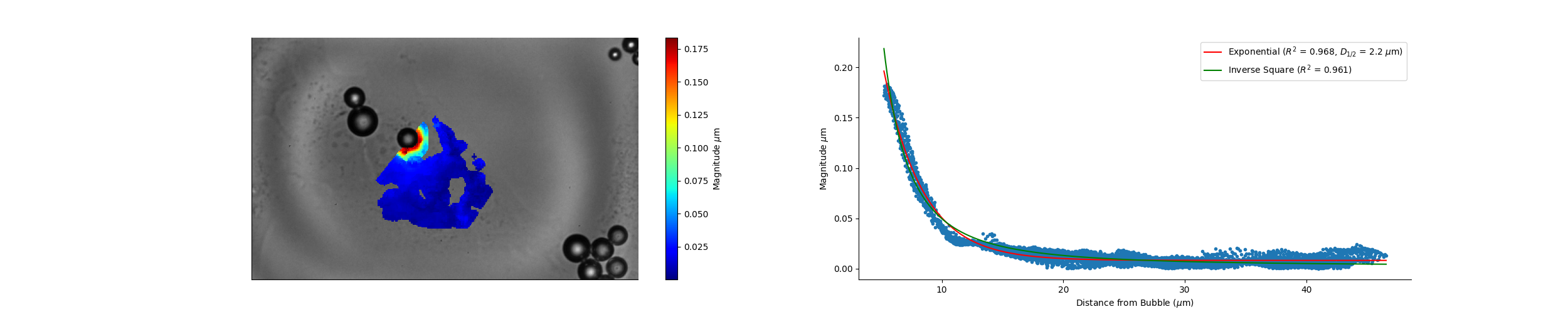

### Camera_18_56_39_Wave_Speed.png

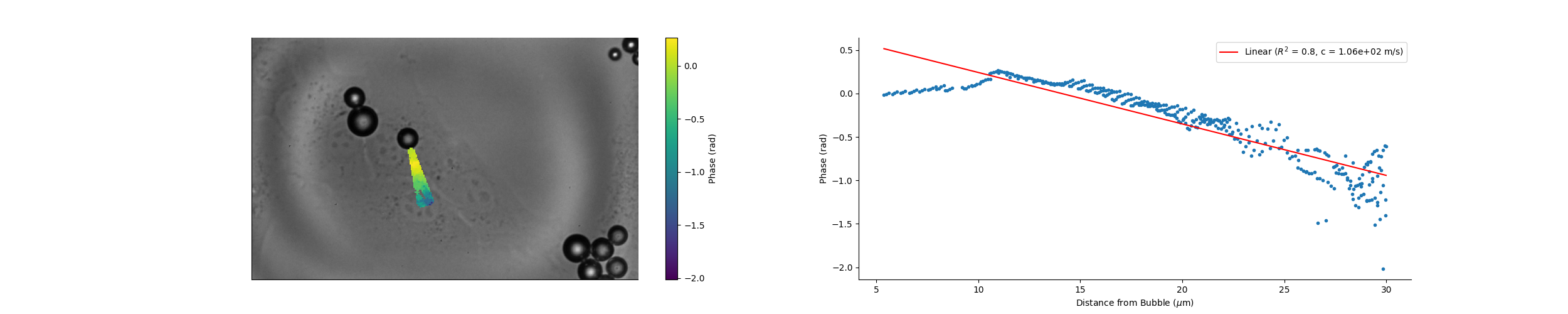

### Camera_18_58_10_Magnitude.png

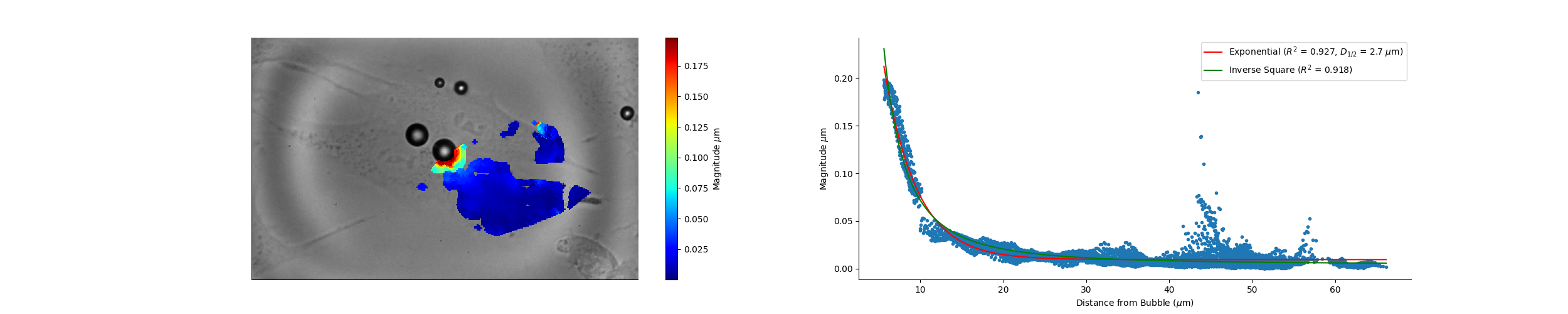

### Camera_18_58_10_Wave_Speed.png

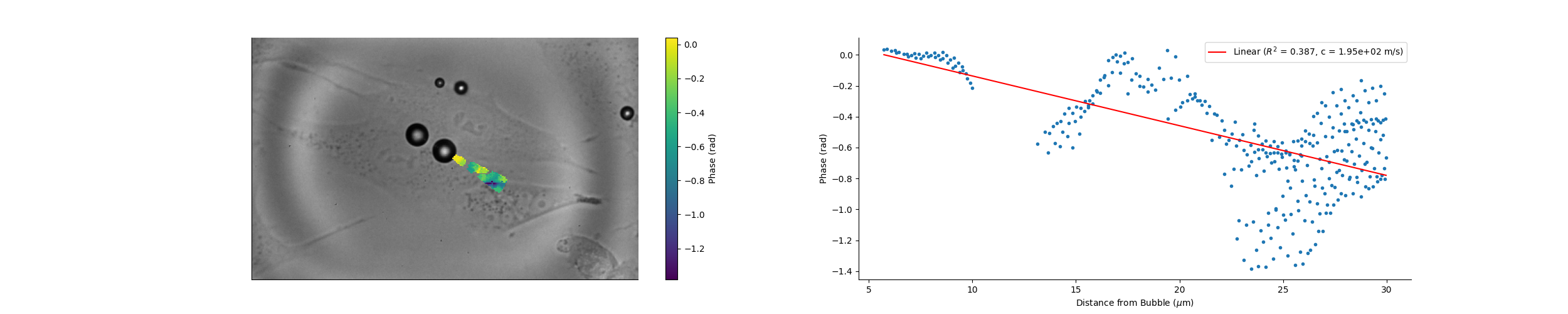

### Camera_19_05_02_Magnitude.png

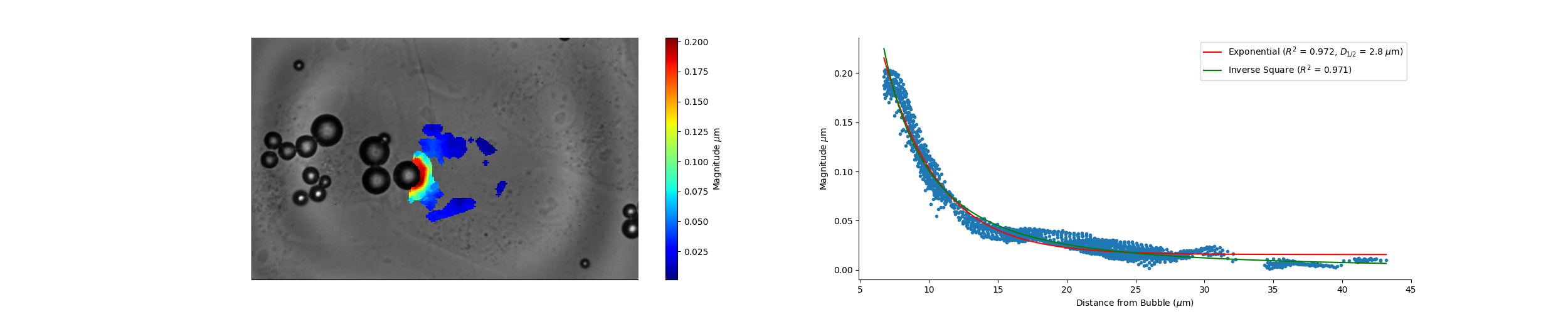

### Camera_19_05_02_Wave_Speed.png

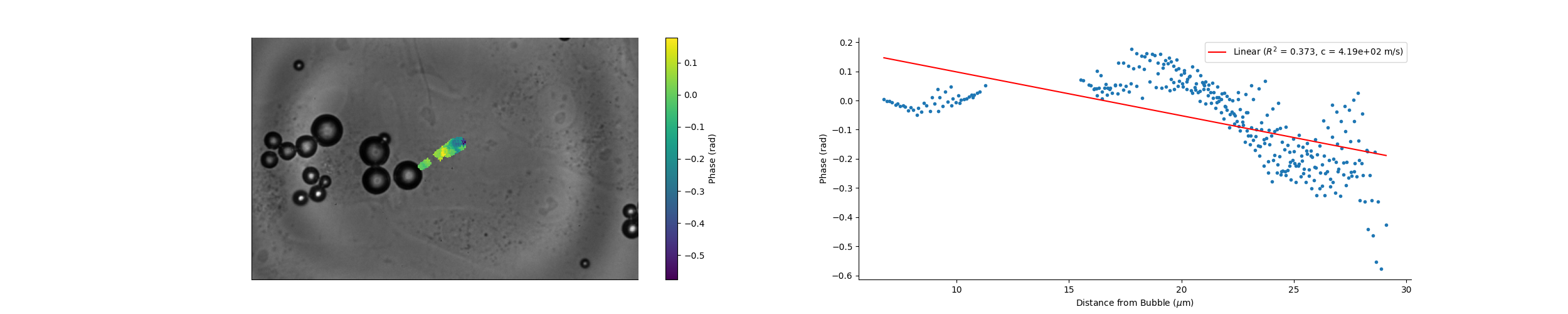

### Camera_19_21_29_Magnitude.png

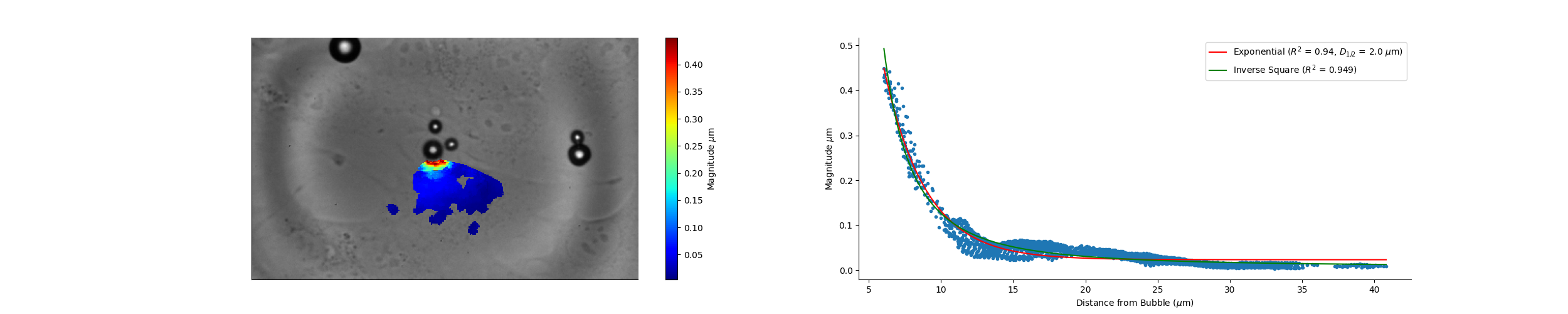

### Camera_19_21_29_Wave_Speed.png

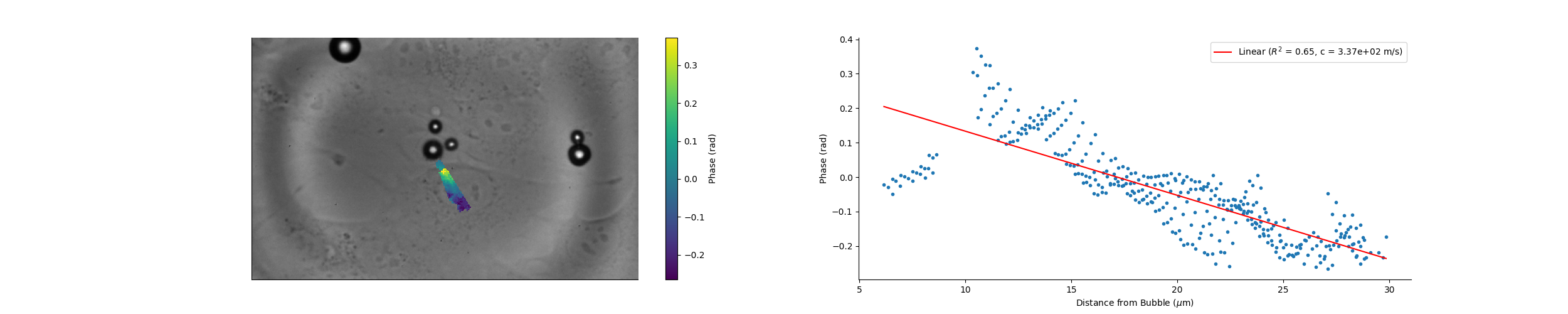

### Camera_19_22_47_Magnitude.png

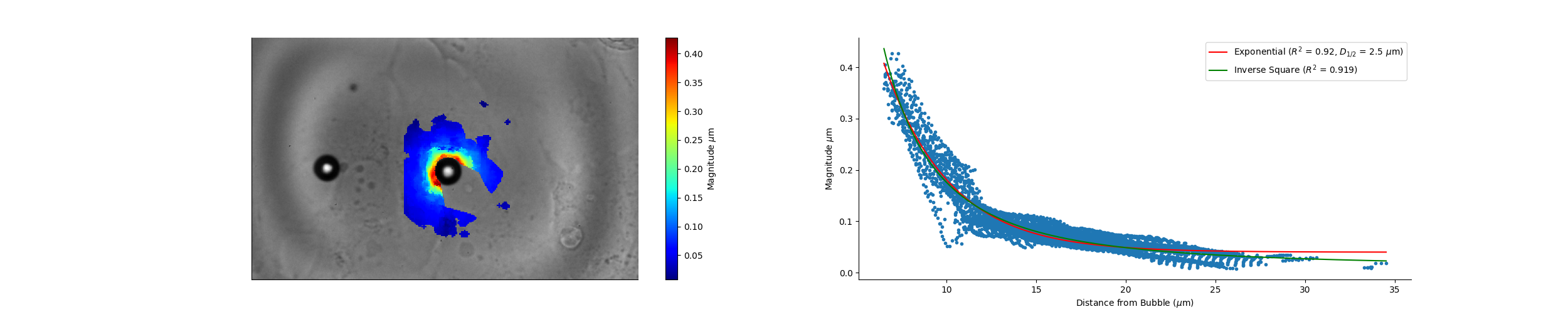

### Camera_19_22_47_Wave_Speed.png

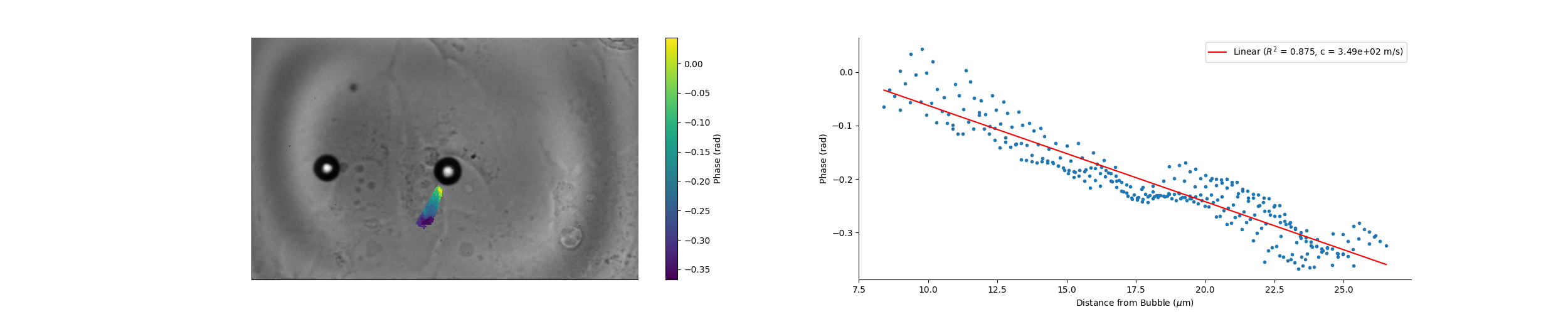

### Camera_19_32_12_Magnitude.png

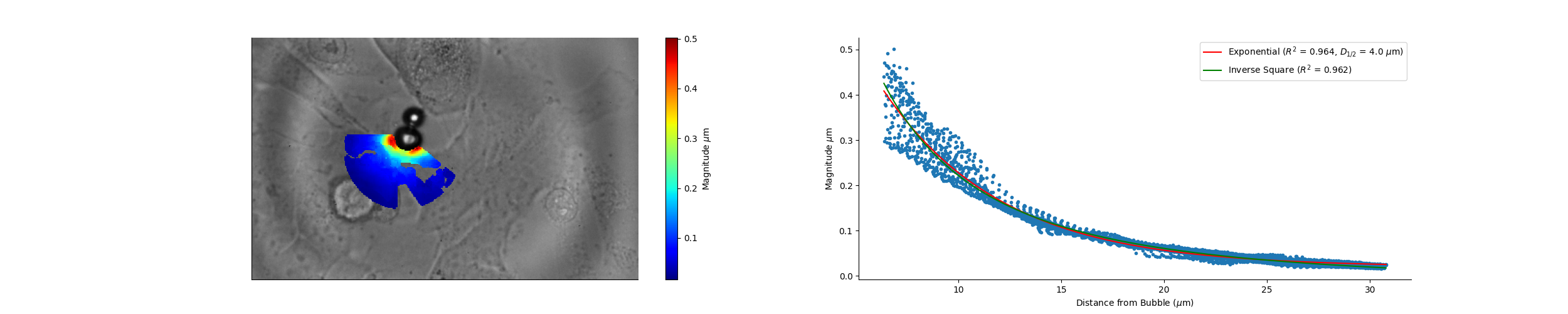

### Camera_19_32_12_Wave_Speed.png

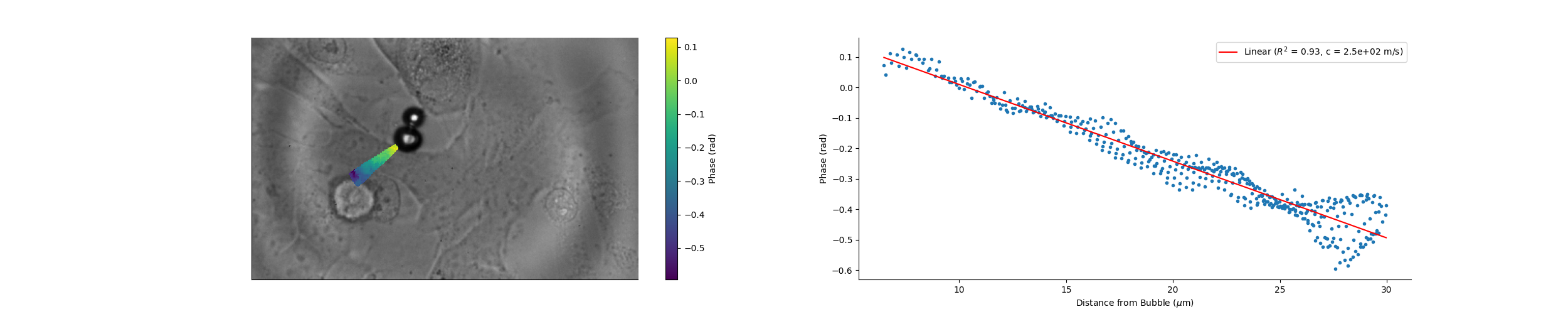

### Camera_19_32_53_Magnitude.png

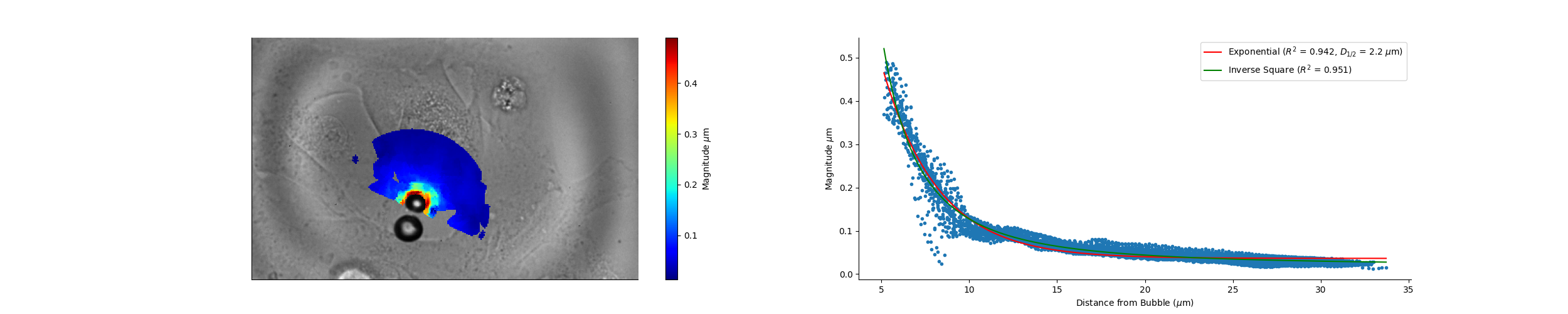

### Camera_19_32_53_Wave_Speed.png

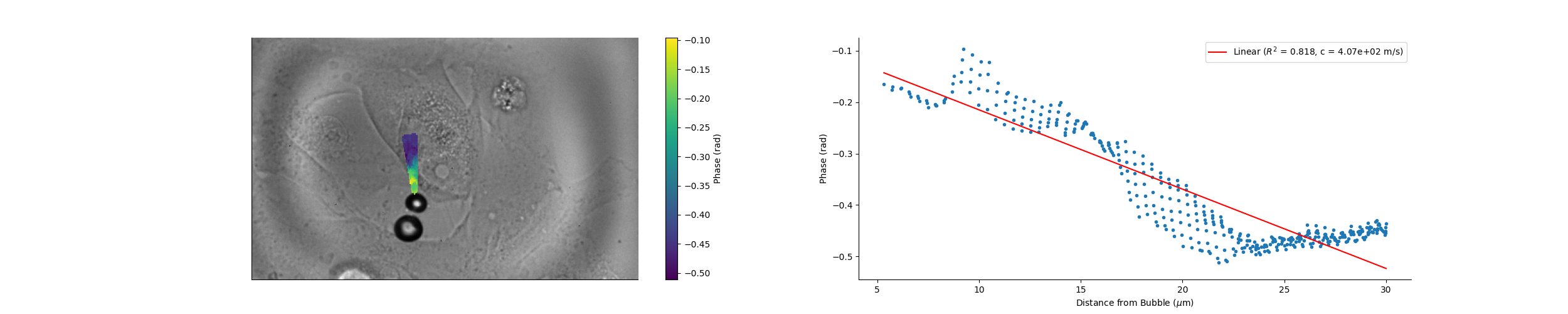

### Camera_19_41_40_Magnitude.png

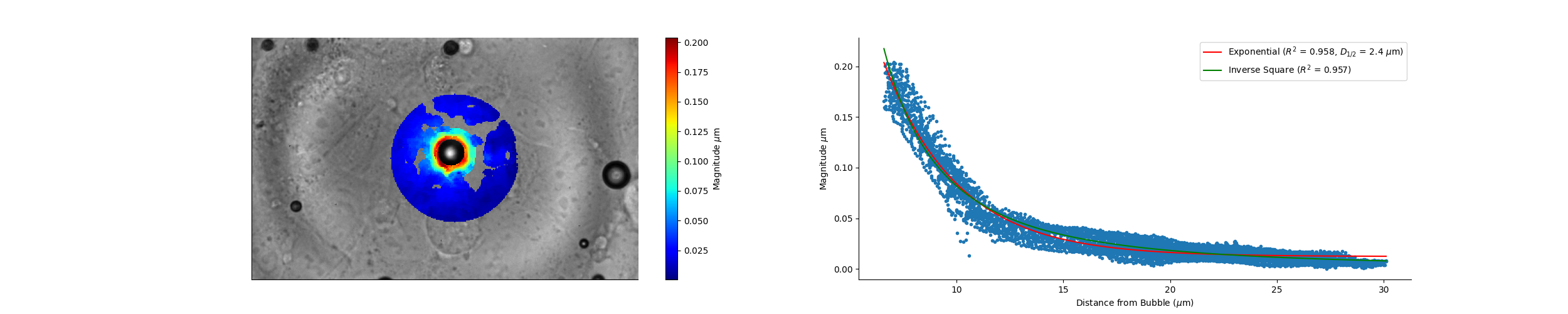

### Camera_19_48_01_Magnitude.png

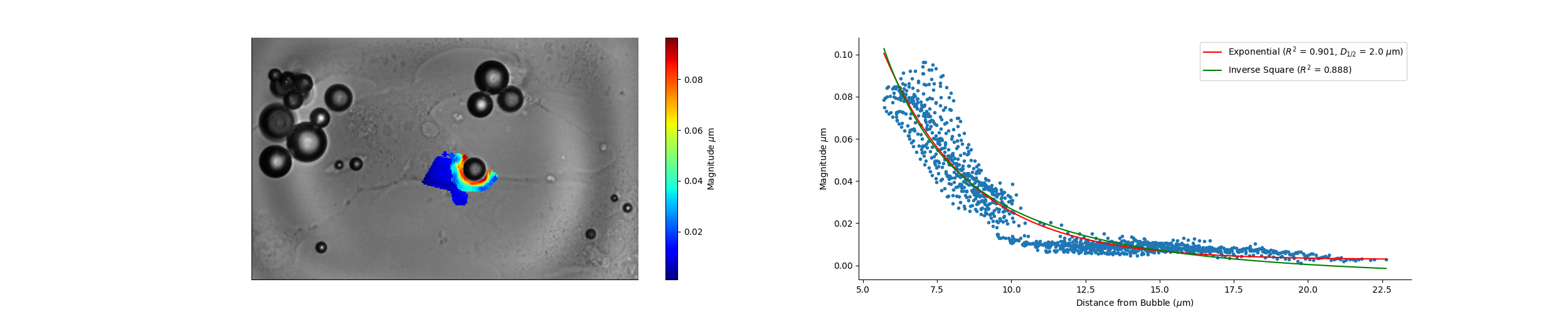

### Camera_19_51_43_Magnitude.png

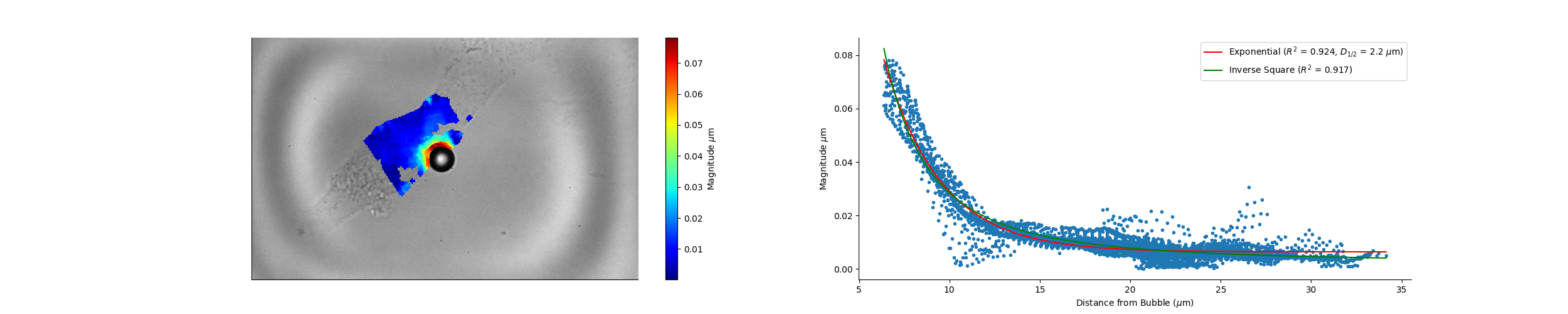

### Camera_19_51_43_Wave_Speed.png

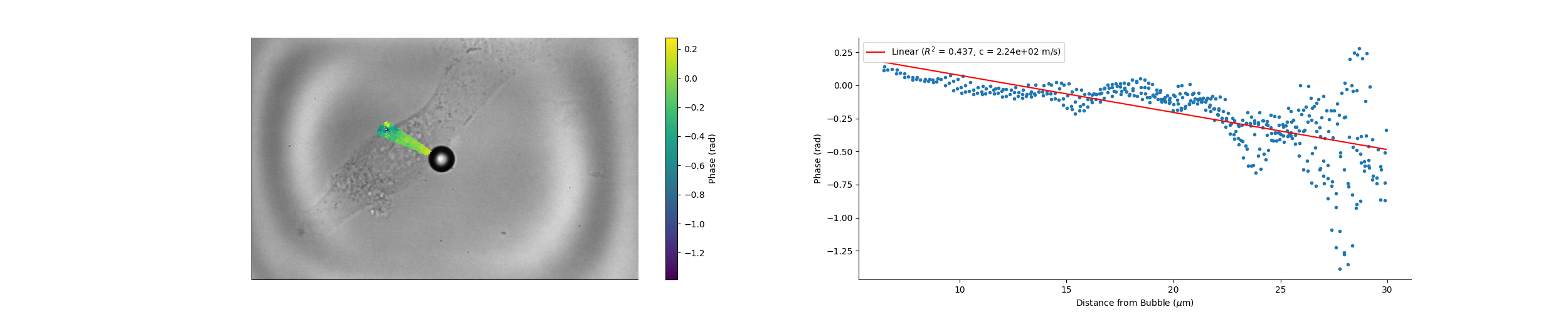

### Camera_19_54_49_Magnitude.png

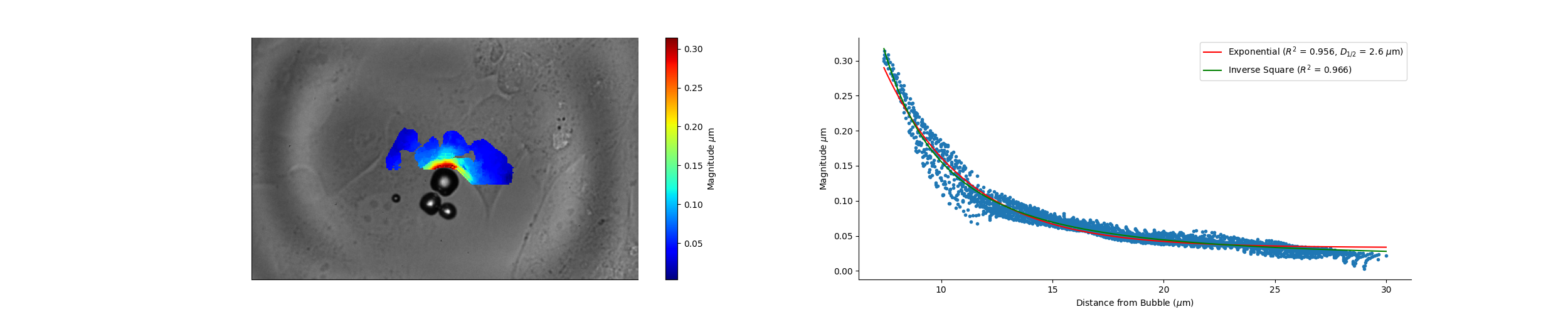

### Camera_19_54_49_Wave_Speed.png

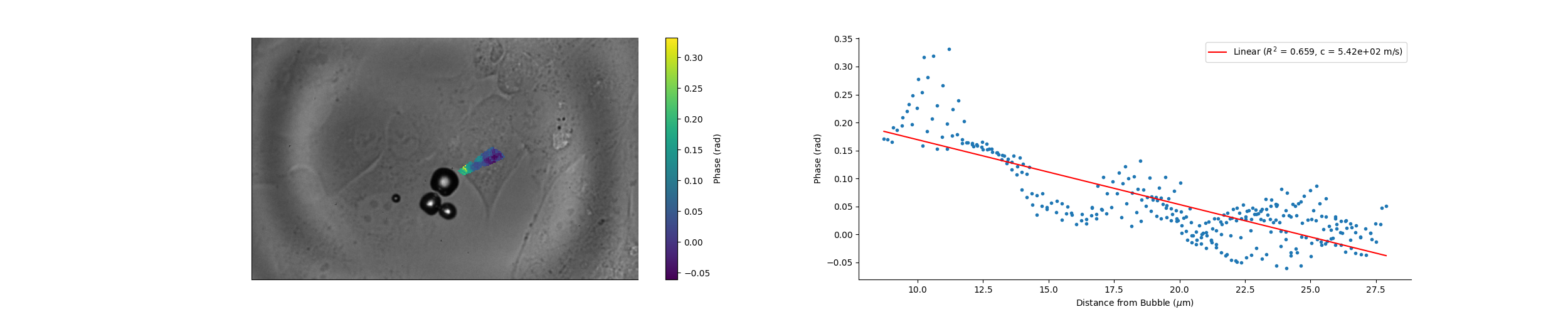

### Camera_19_55_58_Magnitude.png

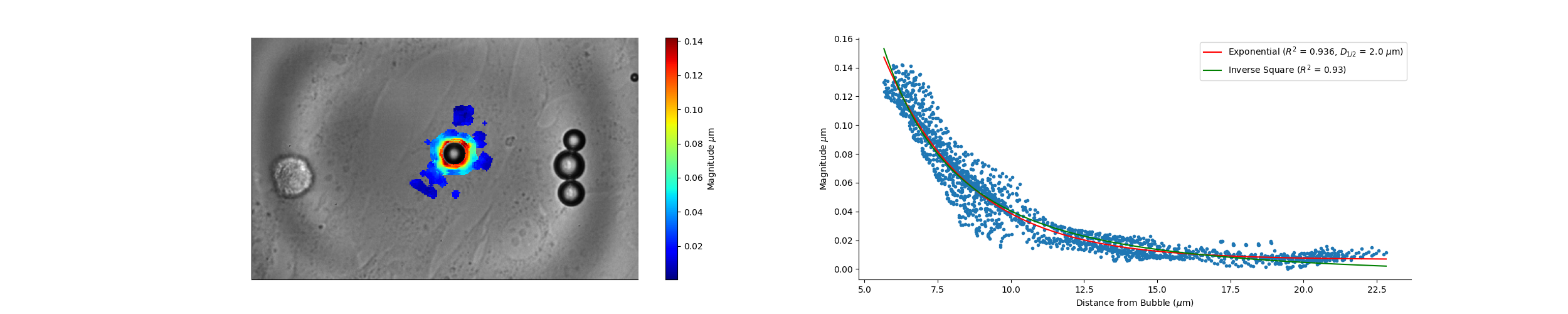

### Camera_19_57_58_Magnitude.png

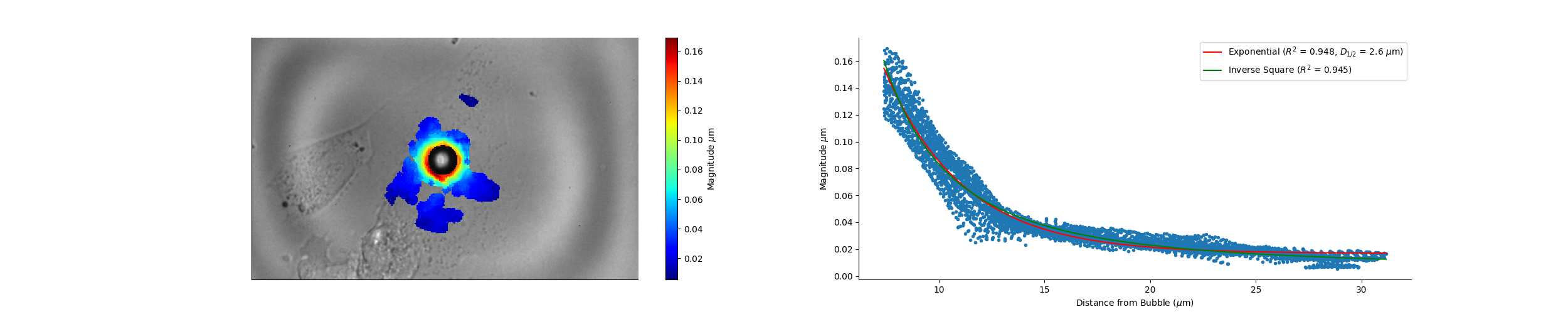
