## Supplementary Information 2 for "Cell material state determines high-frequency cell deformation and microbubble-induced permeabilization"

**Supplementary Information 2: Wave Speed Derivation**

Consider the harmonic oscillation;

$$y(t,x) = \mathrm{A(x)sin}\left(2\pi f\left(t - \partial t(x)\right)\right) = \mathrm{A(x)sin}\left(2\pi f t + \phi_{ROI}\right)$$

where $y(x,t)$ is the deformation of a point at a distance $x$ from the microbubble and at time $t$, $A(x)$ is the amplitude of the oscillation at a point at a distance $x$ from the microbubble, $f$ is the driving frequency in Hz, $\partial t(x)$ is the lag time of a point a distance $x$ from the microbubble and $\phi_{ROI}(x) = -2\pi f \partial t(x)$ is the phase change, in radians, of a point a distance $x$ from the microbubble. Therefore;

$$\frac{\partial \phi_{ROI}(x)}{\partial x} = -2\pi f \frac{\partial t(x)}{\partial x}$$

or;

$$v_p = \frac{\partial x}{\partial t(x)} = \frac{-2\pi f}{\frac{\partial \phi_{ROI}(x)}{\partial x}}$$

Assuming linearity, $\phi_{ROI}(x) = M_{\phi_{ROI}} x + C$, where $M_{\phi_{ROI}}$ is the fitted gradient of a linear regression and $C$ is a constant. You therefore find, $\frac{\partial \phi_{ROI}(x)}{\partial x} = M_{\phi_{ROI}}$, and Equation 5:

$$v_p = \frac{-2\pi f}{M_{\phi_{ROI}}}$$
