## Supplementary Figures and Tables for "Cell material state determines high-frequency cell deformation and microbubble-induced permeabilization"

### Supplementary Figure 1: Grey scale oscillations

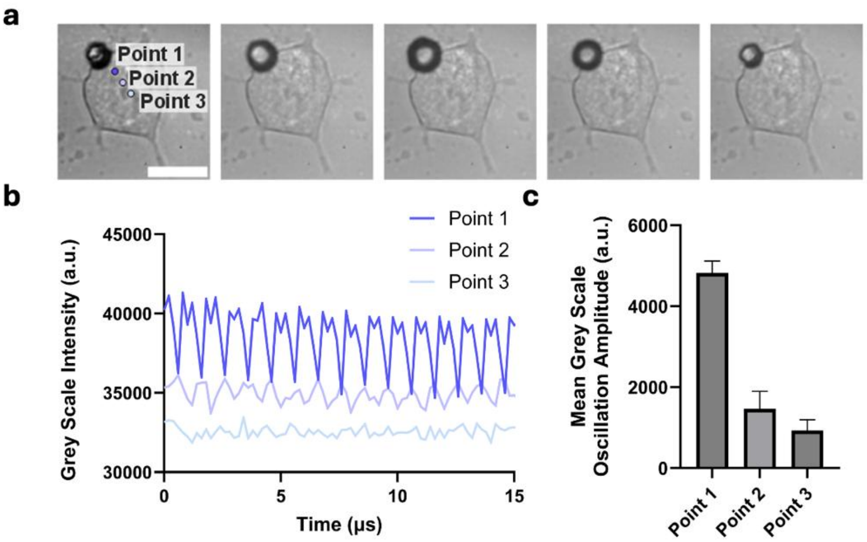


***Figure 2: Using fluctuations in the grey scale from the high-speed imaging to infer possible deformation in the cell cytoplasm.*** **a)** *5 frames from* Video 1 showing ultrasound microbubble-induced displacement on the cell. Points 1, 2 and 3 have been selected at increasing distance, and b) the oscillation in the grey scale values through the first 10 µs is shown. c) The mean amplitude of these oscillations is then plotted for each point, showing a clear decrease with distance from the microbubble.

### Supplementary Figure 2: Additional images of cells treated with cytoskeletal disrupting agents.

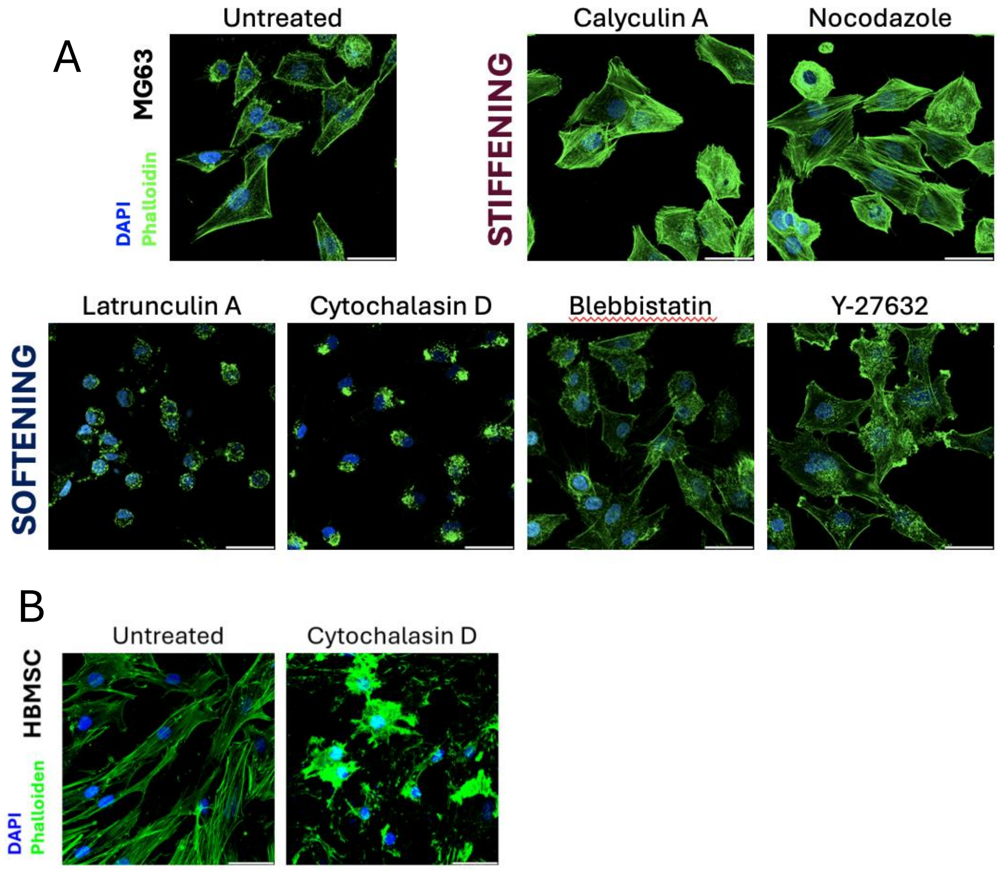


***Figure 1:*** *Confocal images with DAPI-stained nuclei (blue) and phalloidin-stained actin (green) for untreated, calyculin or nocodazole-treated (stiffening agents) and latrunculin A, cytochalasin D, blebbistatin, or Y-27632-treated (softening agents) in (A) MG63 cells and (B) BMSCs (scale bar = 50 µm).*

### Supplementary Figure 3: Sonopermeablisation data with controls (Figure 3f & h)

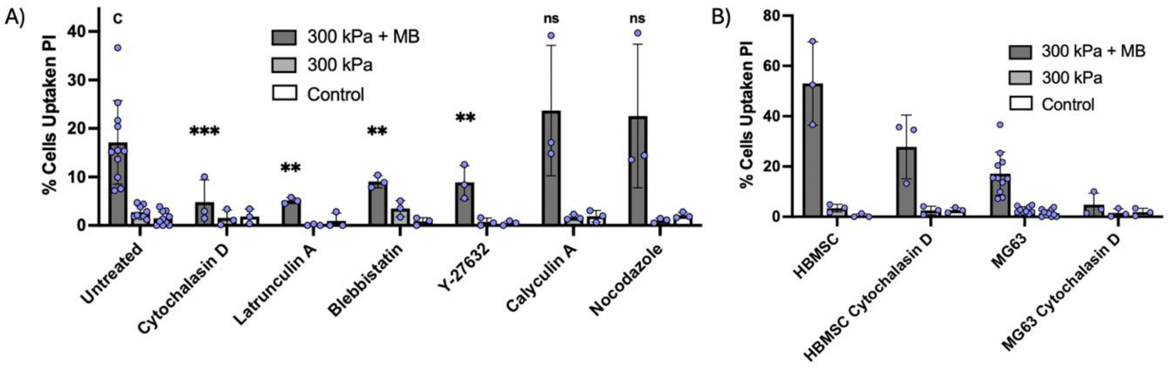


***Figure 3: Sonopermeabilisation data showing % PI uptake in;* a)** *untreated MG63 cells or treated with cytochalasin D, latrunculin A, blebbistatin, Y-27632, calyculin A, and nocodazole and* **b)** *HBMSCs compared with MG63 cells untreated and treated with cytochalasin D. Both figures contain microbubble free (300 kPa) and a microbubble and ultrasound-free (control) group.*

### Supplementary Table 1: Statistical analysis of the effect of cytoskeletal inhibitors on $D_{1/2}$

*Appendix Table 3: All P values relating to the half decay distance between treatment groups (Figure 3b)*

| | Calyculin A | Nocodazole | Untreated | Latrunculin A | Cytochalasin D | Blebbistatin | Y-27632 |
|---|---|---|---|---|---|---|---|
| Y-27632 | 1 | 0.6012 | 0.7254 | 0.4813 | 0.9798 | 0.9996 | |
| Blebbistatin | 0.1543 | 0.1056 | 0.0457 | 0.9738 | 0.9994 | | |
| Cytochalasin D | 0.0031 | 0.0049 | 0.4081 | 0.7888 | | | |
| Latrunculin A | 0.0001 | 0.0495 | 0.1403 | | | | |
| Untreated | 0.0033 | 0.0145 | | | | | |
| Nocodazole | 0.0004 | | | | | | |
| Calyculin A | | | | | | | |

### Supplementary Table 2: MatchID Hardware and Software Tables

*Appendix Table 1: DIC Hardware Parameters*

| | |
|---|---|
| ***Camera*** | Shimadzu Hypervision HPV-X Camera |
| ***Image Resolution*** | 400 × 250 pixels$^2$ |
| ***Microscope*** | Olympus IX71 |
| ***Magnification*** | 80× |

| | |
|---|---|
| ***Image scale*** | 0.42 µm/pixel |
| ***Working Distance*** | 10 mm |
| ***Frame Rate*** | 5 million frames per second (FPS) |
| ***Patterning Technique*** | Natural speckle pattern in cells |
| ***Laser*** | Cavitar CAVILUX |
| ***Exposure*** | 110 ns |
| ***Laser Pulse*** | 50 ns |

***Appendix Table 2: DIC Hardware Parameters***

| | |
|---|---|
| ***DIC Software*** | MatchID 2D 2024.2.5 |
| ***Image Filtering*** | Gaussian 5×5 kernel |
| ***Subset Size*** | 17 pixels |
| ***Step Size*** | 1 pixel |
| ***Matching Criteria*** | Zero Normalised Sum of Square Differences (ZNSSD) |
| ***Interpolation*** | Bi-cubic Spline |
| ***Camera Noise*** | 0.335% |
| ***Displacement Noise*** | 0.1 pixel (40 nm) |

### Supplementary Information 1: All data can be found in zipped and indexed folder

### Supplementary Information 2: Wave Speed Derivation

Consider the harmonic oscillation;

$$y(t,x) = \mathrm{A(x)}\sin\left(2\pi f\left(t - \partial t(x)\right)\right) = \mathrm{A(x)}\sin\left(2\pi f t + \phi_{ROI}\right)$$

where $y(x,t)$ is the deformation of a point at a distance $x$ from the microbubble and at time $t$, $A(x)$ is the amplitude of the oscillation at a point at a distance $x$ from the microbubble, $f$ is the driving frequency in Hz, $\partial t(x)$ is the lag time of a point a distance $x$ from the microbubble and $\phi_{ROI}(x) = -2\pi f \partial t(x)$ is the phase change, in radians, of a point a distance $x$ from the microbubble. Therefore;

$$\frac{\partial \phi_{ROI}(x)}{\partial x} = -2\pi f \frac{\partial t(x)}{\partial x}$$

or;

$$v_p = \frac{\partial x}{\partial t(x)} = \frac{-2\pi f}{\frac{\partial \phi_{ROI}(x)}{\partial x}}$$

Assuming linearity, $\phi_{ROI}(x) = M_{\phi_{ROI}} x + C$, where $M_{\phi_{ROI}}$ is the fitted gradient of a linear regression and $C$ is a constant. You therefore find, $\frac{\partial \phi_{ROI}(x)}{\partial x} = M_{\phi_{ROI}}$, and Equation 5:

$$v_p = \frac{-2\pi f}{M_{\phi_{ROI}}}$$
